# Tertiary-interaction characters enable fast, model-based structural phylogenetics beyond the twilight zone

**DOI:** 10.1101/2023.12.12.571181

**Authors:** Caroline Puente-Lelievre, Ashar J. Malik, Jordan Douglas, David Ascher, Matthew Baker, Jane Allison, Anthony Poole, Daniel Lundin, Matthew Fullmer, Remco Bouckert, Hyunbin Kim, Martin Steinegger, Nicholas Matzke

## Abstract

Protein structure is more conserved than protein sequence, and therefore may be useful for phylogenetic inference beyond the “twilight zone” where sequence similarity is highly decayed. Until recently, structural phylogenetics was constrained by the lack of solved structures for most proteins, and the reliance on phylogenetic distance methods which made it difficult to treat inference and uncertainty statistically. AlphaFold has mostly overcome the first problem by making structural predictions readily available. We address the second problem by redeploying a structural alphabet recently developed for Foldseek, a highly-efficient deep homology search program. For each residue in a structure, Foldseek identifies a tertiary interaction closest-neighbor residue in the structure, and classifies it into one of twenty “3Di” states. We test the hypothesis that 3Dis can be used as standard phylogenetic characters using a dataset of 53 structures from the ferritin-like superfamily. We performed 60 IQtree Maximum Likelihood runs to compare structure-free, PDB, and AlphaFold analyses, and default versus custom model sets that include a 3DI-specific rate matrix. Analyses that combine amino acids, 3Di characters, partitioning, and custom models produce the closest match to the structural distances tree of Malik et al. (2020), avoiding the long-branch attraction errors of structure-free analyses. Analyses include standard ultrafast bootstrapping confidence measures, and take minutes instead of weeks to run on desktop computers. These results suggest that structural phylogenetics could soon be routine practice in protein phylogenetics, allowing the re-exploration of many fundamental phylogenetic problems.

## Introduction

Protein structural phylogenetics is the “missing chapter in molecular evolutionary biology” (Lai 2020). It is well-known that protein structure is often remarkably conserved (Chothia & Leak 1986; Flores et al. 1993) even in the “twilight zone” (Doolittle 1986; Chung & Subbiah 1986), where amino acid (AA) sequence similarity decays below 30%. Very low sequence similarity limits our ability to detect homology, accurately align sequences, and resolve deep relationships. It would therefore be advantageous to incorporate protein structural information into phylogenetic inference.

### Structural phylogenetics: progress and limitations

Researchers have taken a variety of approaches to link structure to phylogenetic inference. Lake (1984) used visual interpretation of low-resolution quaternary structures of the ribosome to identify gross characteristics that seemed to support the placement of eukaryotes within archaea. More generally, except at the coarsest levels, structural classifications of proteins suggest common ancestry (e.g. SCOP Superfamily, CATH Homology; Andreeva et al. 2020; Sillitoe et al. 2019), but they do not resolve within-group relationships, nor do they constitute a formal estimate of a phylogenetic tree (Lundin et al. 2012, Malik et al. 2020).

Secondary structure (SS) and residue accessibility have been used to modify amino acid substitution processes (Thorne et al. 1996; Le & Gascuel 2010), including joint estimation of the secondary structure (Goldman et al. 1996; Goldman et al. 1998). A probabilistic model of amino acid dihedral angle evolution has been used to model pairwise evolution between structures (Golden et al. 2017), but has not been generalized for phylogenetic inference. Modifications to substitution rates based on tertiary structure have also been explored (Choi et al. 2007). Lai et al. (2020) extended this work to include seven DSSP-defined (Kabsch & Sander 1983) secondary structure (SS) states as character data, estimating substitution matrices from prealigned PDB structures.

An alternative approach to these methods is to use 3D structural superposition of structures. Structural superposition is already popular in the related problem of structure-informed sequence alignment (e.g., DALI, Holm & Sander 1995; TM-Align, Zhang & Skolnick 2005; SSM, Krissinel & Henrick 2004; US-align, Zhang et al. 2022; SWAMPNN, Trinquier et al. 2022). For phylogenetics, a matrix of pairwise distances between all structures is calculated using a metric such as root mean square positional deviation (RMSD), or a welter of others (Hasegawa & Holm 2009), including more robust metrics such as TM-Score (Zhang & Skolnick 2005), LDDT (Mariani et al. 2013), or Q_score_ (Krissinel & Henrik 2004; Malik et al. 2020). Following earlier papers using small numbers of structures (≤18; Bujnicki 2000; Breitling et al. 2001; Garau et al. 2005), Lundin et al. (2012) used distance matrices to estimate a phylogenetic network using NeighborNet with 83 PDB structures of the ferritin-like superfamily and broadly replicated previous sequence-based results, as well as structural database classifications and functionally-delineated groups.

Structural phylogenetics with distance methods is a simple approach, and thanks to the AlphaFold2 revolution (Jumper et al. 2021; Skolnick et al. 2021), is no longer hindered by a shortage of structural data. However, such approaches are restricted by the use of distance methods to create trees, such as Neighbor Joining (NJ). For phylogeneticists, distance methods are often considered rather “quick and dirty”: NJ provides a reasonable first approximation, but its statistical consistency is contingent on the accuracy of the distance matrix (Bryant 2005, Gascuel and Steel 2006). The accuracy of distances is well-defined for nucleic and amino acids, where it measures the number of nucleotide or amino acid substitutions. However, for structural distances, it is not known whether LDDT (Mariani et al. 2013), TM-score (Zhang & Skolnick 2004), or others are fair metrics of structural change, where the complex true process would include not only changes in AAs, but also other changes such as binding partner or environment. Another issue is that protein conformations vary, sometimes substantially, on nanosecond timescales. Nevertheless, work to date (Chothia & Lesk 1986; Flores et al. 1993; Lundin et al. 2012; Lai et al. 2020; Malik et al. 2020) suggests that structural distances correlate with the better understood metric of sequence divergence.

A further fundamental limitation of NJ used on structural distances (unlike nucleic or amino acid sequences) is the lack of probabilistic models describing generative processes. Probabilistic models have many advantages, including statistical model comparison and selection, and standard procedures to quantify statistical uncertainty, such as bootstrapping and posterior probabilities. Moreover, generative models are explanatory and capture real biological processes, allowing use of *in silico* simulations for model validation. Lastly, probabilistic models open the pathway towards “total-evidence” approaches (Hulsenbeck et al. 1996) allowing multiple sources of data to be combined into a single analysis. It is challenging to conceptualize a generative model that realistically describes how secondary, tertiary, and quaternary structure co-evolves over evolutionary time, but models of some aspects exist (Boomsma et al. 2008)

A method to ameliorate some of the above issues was proposed by Malik et al. (2020). Here, Molecular Dynamics (MD) simulations were used to sample different conformations of each leaf structure. Superposition, distance calculation, and NJ trees were calculated for each sample of tip conformations, and MD-based bootstrap scores were calculated for each branch, providing a measure of the effect of structure variability on the stability of the estimated relationships. The method was used on a subset of Lundin et al.’s ferritin-like superfamily dataset (53 PDB structures), retrieving similar relationships but with the advantage of reporting bootstrap supports. Disadvantages of this approach include the slowness of the MD simulation step, its reliance on distance methods, and the fact that sources of uncertainty other than conformational variation are not addressed.

In summary, research to date indicates that structural data could have great utility in phylogenetics, in particular beyond the twilight zone, if the challenges of probabilistic modeling, uncertainty estimation, and computational runtime can be overcome (Bujnicki 2000; Breitling et al. 2001; Garau et al. 2005; Lundin et al. 2012; Lai et al. 2020; Malik et al. 2020). Combined with reasonably accurate AlphaFold structures now available for many proteins, such a technique could rapidly make structural phylogenetics a routine part of protein phylogenetics.

### 3Di characters

A potential solution to the above challenges is provided by Foldseek (van Kempen et al. 2023), a program designed for efficient structure-informed homology searches deep in the twilight zone. Foldseek’s 3Di alphabet encodes 3-D tertiary structure interactions in a 1-D string of discrete states, enabling deployment of highly efficient dynamic programming algorithms for alignment and search.

A summary of Foldseek’s procedure classifying residues into 1 of 20 3Di states is given in Figure 1. For a full description of the classification procedure, and the derivation and training of the 3Di alphabet, see Van Kempen et al. (2023). The 3Di alphabet was originally designed to enable fast and sensitive homology search and structural alignment, and the alphabet was trained for that purpose. The encoding of tertiary structure interactions seems to retain the signal of core conserved folds in a protein, and to be robust to changes outside the core (such as indels) that challenge traditional superposition metrics. Sensitivity in homology search suggests that the 3Di states could be interpreted as highly conserved, phylogenetically informative *characters*. In combination with the AlphaFold database, this is an immediately appealing idea for phylogeneticists, as the Foldseek encoder could then be used to create a 3Di character dataset to compare to, or supplement, any traditional amino acid dataset. The fact that the 3Di alphabet has 20 states is particularly convenient, as it allows 3Di characters to be input into traditional bioinformatic and phylogenetics programs. It is tempting to think this brings us a step closer to the “Holy Grail”: a set of easily-obtainable, highly conserved characters that can be added to any traditional protein phylogenetics project, enabling greater resolution deeper in the tree.

**Figure 1.**
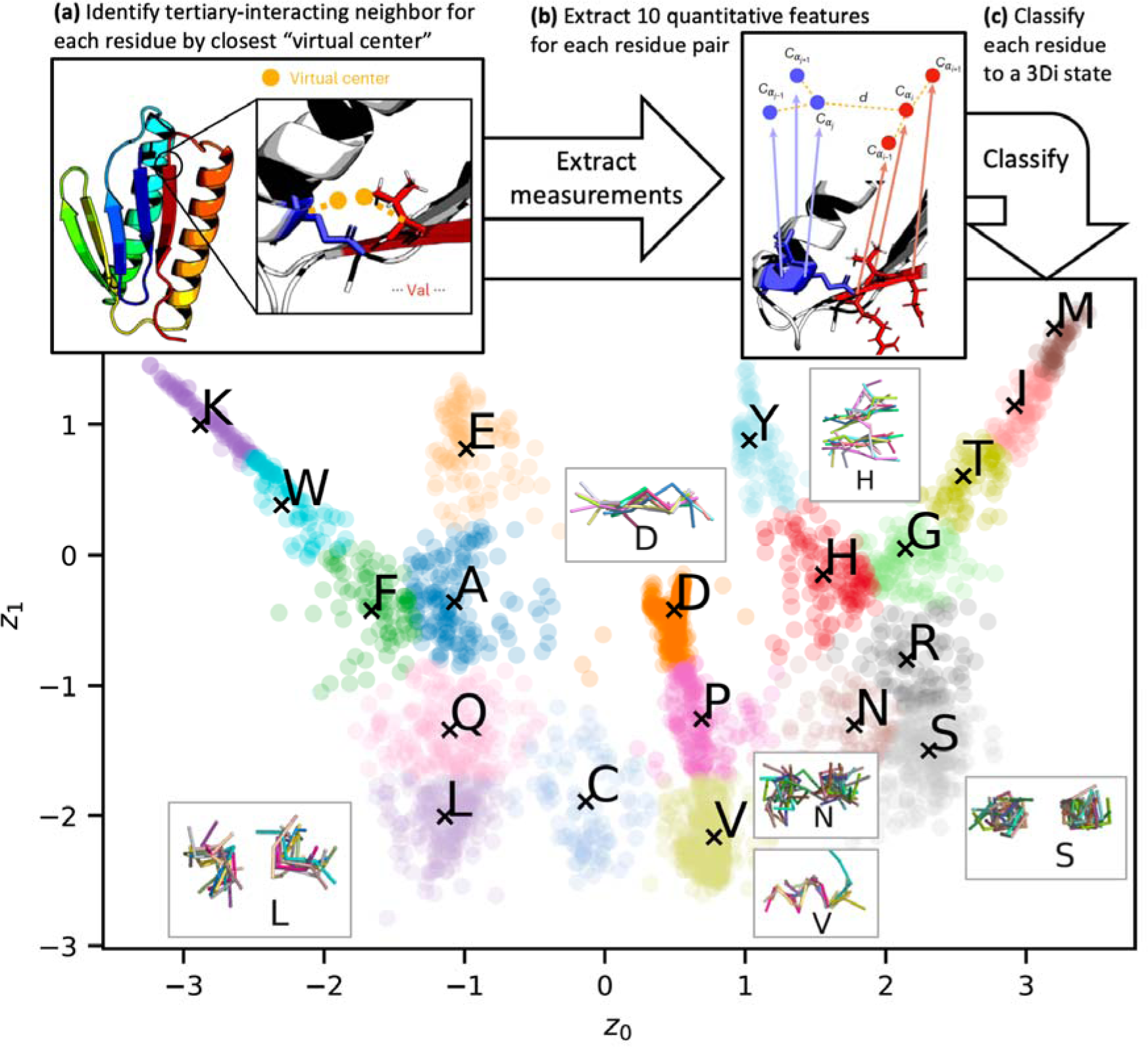
Summary of Foldseek’s system for classifying tertiary interactions into the 20-state 3Di alphabet. For each residue in a protein structure **(a)** a virtual center is defined, and each residue is assigned a tertiary neighbor based on the closest virtual center. **(b)** 10 quantitative traits are collected for each residue+neighbor interaction, and then **(c)** assigned to the best-matching 3Di state using the classifier pre-trained by Van Kempen et al. 2023. The main panel shows a two-dimensional representation of the 20 states. Examples of 3Di states are inset; each of these represents 10 pairs of backbone fragments showing the typical arrangement for a 3Di state. Graphics modified from Van Kempen et al. 2023.

The creators of Foldseek have already taken one step in this direction with FoldTree, which uses Foldseek’s 3Di alphabet to structurally align protein structures and AA sequences, and then calculate a “Fident” (fraction identical) distance between the AA sequences between pairs of structures (Moi et al. 2023). Although they obtain good results with this approach (higher taxonomic congruence and more ultrametricity), trees are estimated from distance matrices using FastME (NJ followed by NNI/SPR optimization for the minimum evolution tree; Lefort et al. 2015), and therefore subject to the limitations discussed above.

We propose that 3Di codes can be interpreted as conventional phylogenetic characters, and thus can be readily included in the suite of standard model-based phylogenetic estimation procedures, including statistical model comparison, maximum likelihood phylogenetic estimation, and bootstrapping. If it is true that 3Di characters are indeed the “Holy Grail,” a source of slowly-evolving characters, then inferences including 3Di characters should outperform standard, structure-free AA-based inferences in the twilight zone.

As a proof-of-concept, we use both PDB and Alphafold2-predicted structures from the ferritin-like superfamily, a group that shares the ferritin fold. From the perspective of evolutionary enzymology, and hence phylogenetics, the ferritin fold is interesting. It likely evolved to control the Fenton reaction (Fenton, 1894), in which ferrous iron reacts with hydrogen peroxide to form highly reactive hydroxyl radicals. From this starting point it evolved into several major types (Lundin et al. 2012): rubrerythrins, and similar small proteins, control reactive oxygen species and are hence likely close to the ancestral function; the ferritins, bacterioferritins and “DPS” proteins store insoluble ferric iron in the cell or protect from radicals; and lastly, a large group of enzymes utilize highly reactive oxygen species for catalysis (fatty acid desaturases, bacterial multicomponent monooxygenases, BMMs) or radical formation (ribonucleotide reductase R2 components, RNR R2). This superfamily has previously been used as a test dataset for structural phylogenetics (Lundin et al. 2012; Malik et al. 2020) as homology is likely, but inter-family relationships are in the twilight zone.

## Methods

### Structure Data

The 53 PDB structures used by Malik et al. (2020; henceforth M20) were downloaded from RCSB.org (Berman et al. 2000; RCSB Protein Data Bank 2023). They were clipped to the single chains used by Malik et al. using *bio3d::pdbsplit* (Grant et al. 2006; Grant et al. 2020). The list of PDB chain files was processed to AA and 3Di FASTA files using the custom R function *str_to_3di* (SI), which runs Foldseek’s *structureto3didescriptor* command and parses the output to FASTA files.

### Reference tree

The M20 structural distances NJ tree was replicated from the original distances matrix using SplitsTree (Huson & Bryant 2006). We used the M20 tree as the comparator for the trees estimated by Maximum Likelihood (ML). While there is no guarantee that the M20 tree is the exact true tree, it at least constitutes the state-of-the-art published estimate which represents the tree obtained from the traditional superposed structural distances approach, its clades reproduced previously-hypothesized groups recognized in structural databases, and bootstrap confidence estimates were available for the deeper branches. M20’s clade labels and MD-based bootstraps were manually transferred to edited Newick files for automated comparison with ML trees.

### AlphaFold-predicted structures

To test whether 3Di characters derived from AlphaFold structures would result in similar phylogenies to empirical structures, AlphaFold structures were downloaded corresponding to each M20 structure. This was accomplished by manually searching AlphaFold DB (https://alphafold.ebi.ac.uk/; Jumper et al. 2021; Varadi et al. 2022; Varadi et al. 2023) against the AA sequence for each M20 structure. All but 9 AA sequences had 100% matches in AlphaFold DB; when no 100% match was found, the closest-hits AlphaFold structure was chosen. The most distant match was 98% identical to the target AA sequence. While 100% identical sequences would be preferable, close-but-inexact matches represent a realistic situation for researchers with a list of study sequences who are aiming to add 3Di characters to their dataset. The average AlphaFold structure has a mean pLDDT of 94.1%, with 90% of all sites scoring over 85.1%, suggesting that structural predictions have high levels of confidence (likely reflecting the existence of close homologs in the protein data bank). See SI for a breakdown of pLDDT summaries.

### Phylogenetic analyses

To test the effect of various analysis decisions, and to see whether or not 3Di codes function like standard phylogenetic characters, we ran a total of 60 maximum likelihood inference runs in IQtree v. 2.2.0 (Minh et al. 2020). Analysis variations used at each step (structures, data, alignment, trimming, models, partitioning) are described below.

### Structures and Data

Analyses (alignment + phylogeny) were run using either no structures (AA-only alignment and phylogeny, matching standard phylogenetic analysis), PDB structures, or AlphaFold structures. For a given alignment, phylogenetic analyses were run on AAs-only, 3Dis-only, or AA+3Di datasets.

### Alignment

Alignment for AA-only, structure-free runs was conducted with FAMSA (Deorowicz et al. 2016), which uses an MIQS substitution matrix useful for distantly-related proteins (Yamada & Tomii 2014). The first set of structure-informed runs used the Universal Structural Alignment (US-align; Zhang et al. 2022) program’s Multiple-Structure Alignment (MSTA) mode to align AAs using the 3D structures themselves, with no involvement of the 3Di characters. The second set of structure-informed runs used FAMSA3di, a version of FAMSA recompiled with Foldseek’s substitution cost matrix for the 3Di alphabet. FAMSA3di was used to align the 3Di characters. As Foldseek produces 3Di sequences of identical length to the AA sequence of a structure, a 3Di alignment can be transferred position-by-position to unaligned AA characters and vice versa, a procedure we implemented in the custom R function *align_3dis_to_AAs*.

### Trimming

The alignment trimmer trimAl 1.4 (Capella-Gutierrez et al. 2009) was used to trim all columns of AA and 3Di alignments that were <35% data. This tended to cut regions of the alignment that were poorly-aligned regions (which were often low-confidence regions of the AlphaFold structures) or represented elements present in only a few structures. ML analyses were run on trimmed as well as full, untrimmed alignments.

### Substitution models

ML analyses of AA, 3Di, and AA+3Di datasets were run under “allmodels”, using IQtree’s extensive default model set, or under a custom list which contained models *a priori* likely to be relevant to AA or 3Di data as well as computationally efficient. The custom list included a custom 3Di rate matrix, which we calculated from Foldseek’s 3Di substitution cost matrix, following the method described by Tammi (2021). The custom models list (IQtree *-mset* input) was: Blosum62, Dayhoff, DCMut, JTT, JTTDCMut, LG, Poisson, PMB, WAG, EX2, EX3, EHO, EX_EHO, 3DI (non-3DI model descriptions available in Minh et al. 2022). The base frequency options used were *-mfreq* FU,F (base frequencies specified by the protein matrix, or empirical base frequencies), and the among-site rate variations used were *-mrate* E,G,R (Equal, Gamma, or FreeRate models of sitewise variation).

### Partitioning

*A priori* it is extremely likely that AA and 3Di characters will evolve under different models, that AA models will be poor fits to 3Di data, that the 3Di matrix will be a superior fit, and that AA+3DI datasets should be partitioned so that each partition can have an appropriate model. However, because this is the first ML analysis using 3Di characters, and as a sanity check of the entire pipeline described above, we decided to run both partitioned and unpartitioned analyses of the AA+3Di alignments. Tree topology and relative branch lengths were shared across partitions.

All IQtree runs were conducted with 1000 ultrafast bootstraps (UFBoot), which approximate Bayesian posterior probabilities, and 1000 replicates of SH-like approximate likelihood ratio test (Guindon et al., 2010). Bootstraps were run with the *-bnni* option, which uses a hill-climbing Nearest-Neighbor Interchange (NNI) on each bootstrap tree to reduce the risk of overestimating UFBoot supports when the data severely violates the model (Minh et al. 2022). All runs were conducted on a desktop 4.2 GHz Quad-Core Intel Mac.

### Analysis of phylogenies

Using custom R scripts (Supporting Information) we extracted from each IQtree run the best fit models and corresponding log-likelihood, AIC, AICc, BIC, number of data and number of parameters. Bootstrapped consensus trees were extracted from each run, midpoint-rooted and ladderized for visualization, and compared to the M20 NJ tree by a variety of metrics. These included: number of matching bipartitions (51 possible); number of matching “deep” bipartitions (20 possible, corresponding to named clades of the M20 tree displayed in Figure 10b of Malik et al. 2020), and groupings of those; number of “shallow” bipartitions (all other M20 bipartitions); total tree length (sum of all branch lengths); sum of bootstrap scores across all branches; and tree-to-tree distances available in phangorn (*treedist* function; Schliep 2010), including Robinson-Foulds distance (symmetric difference), Subtree Pruning-Regrafting (SPR) distance, approximating the number of edits required to match tree topologies; and branch-score distance (calculated on trees where branch lengths had been divided by the total tree length, to avoid artifacts due to comparing trees of very different total lengths). RF and SPR distances consider only topology and ignore branch lengths, unlike the branch-score distance. Nonmetric multidimensional scaling was used to visualize the tree-to-tree distances (Matzke et al. 2015).

## Results

### Alignment

AA and 3Di alignments were visualized and compared using Jalview’s percent-identity color scheme (Waterhouse et al. 2009). 3Di characters show substantially more conservation (116 sites are over 50% conserved) than AAs (only 17 sites) (Figure 2). The conservation pattern is particularly clear in the ferritin superfamily’s 4 core alpha helices (Lundin et al. 2012).

**Figure 2.**
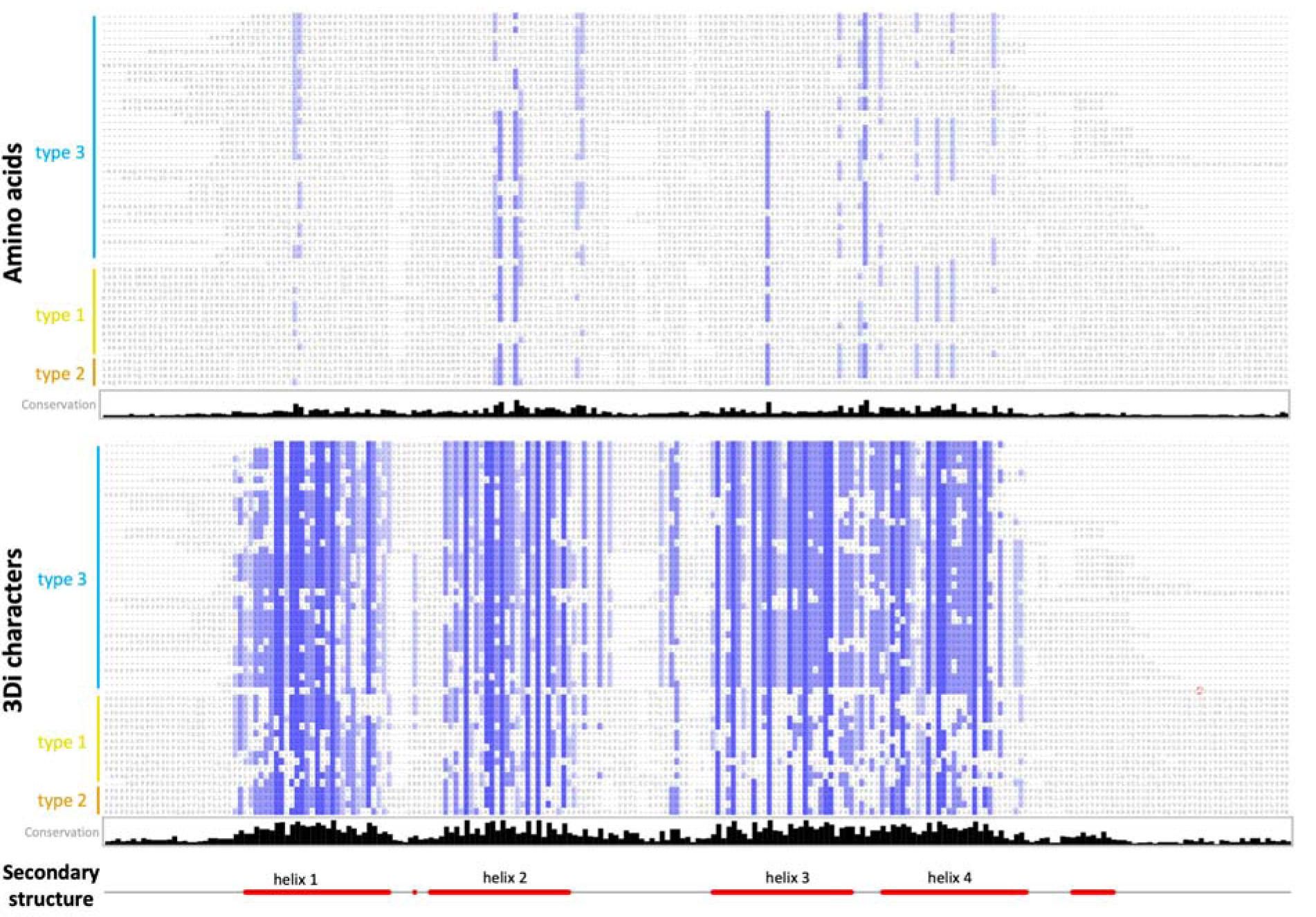
AA and 3Di alignments for 53 AlphaFold structures of the ferritin-like superfamily, displayed with Jalview’s pairwise identity color scheme, where characters matching the majority character (if any) are colored, and darker colors are closer to 100% agreement. The 3Di alignment was generated with famsa3di; the AA alignment was generated by one-to-one replacement of 3Di states with AAs. Sites with <35% data were trimmed. FASTA files are available in SI.

While both alignments have some repetitive structure in the depicted alpha-helical region, the 3Di characters are more conserved and contain information that is distinct from the AAs (Figure 3). For example, in the section of the complete alignment shown in Figure 3, sites 70-71 are both hydrophobic AAs, but do not show obvious grouping information between type 3 and type 1 groups. The corresponding 3Di states show a clear fixed difference at position 70 (V/L vs. S), and invariance at 71 (S). Sites 79-80 show little AA conservation, but the corresponding 3Di sites show fixed differences between the type 1 and type 3 groups. These “fixed” differences do not strictly hold for the full alignment, but type 3 vs. type 1/2 grouping is evident in the 3Di alignment (Figure 2).

**Figure 3.**
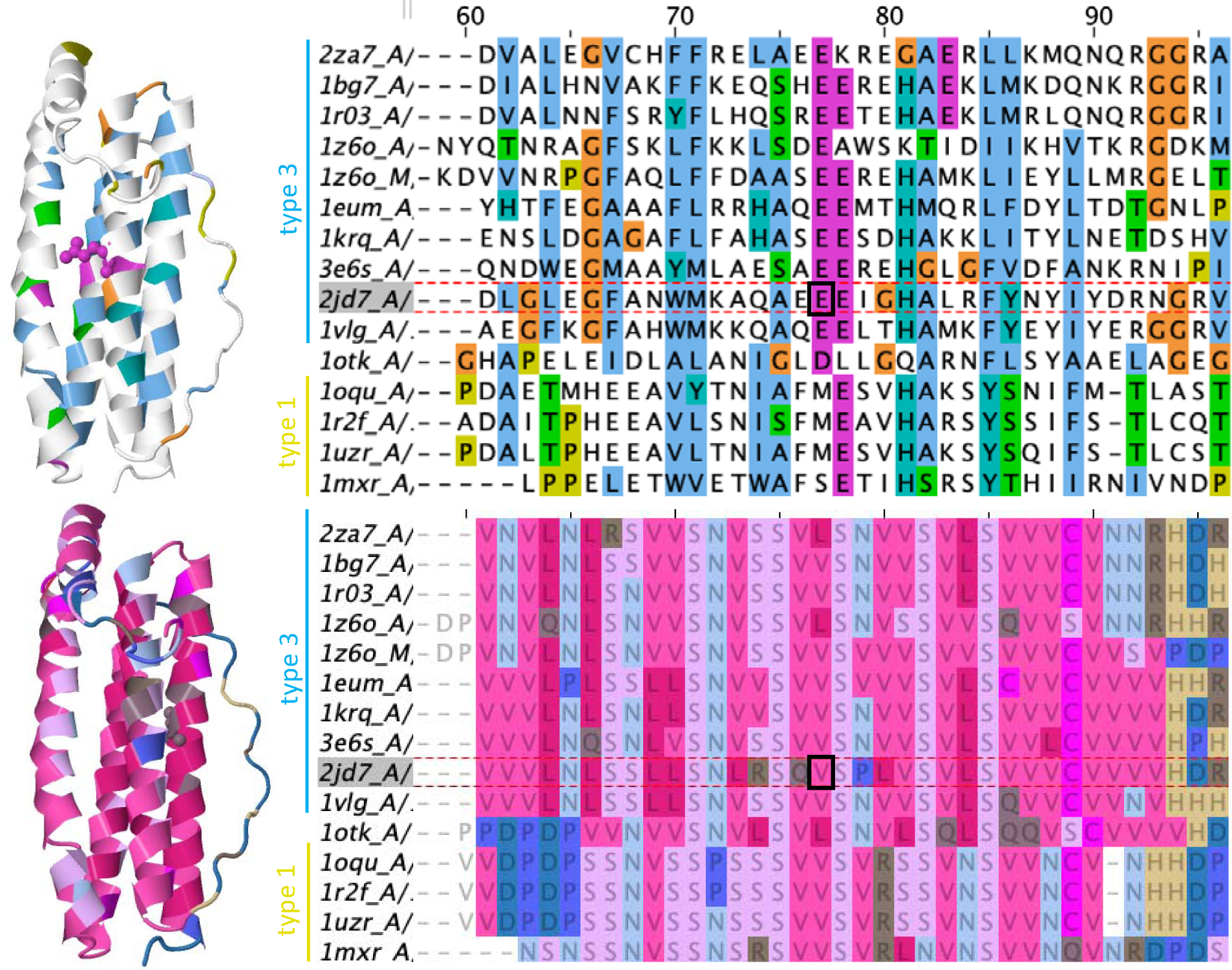
Illustrating the differences between AA and 3Di characters by zooming in on a portion of the alignment in Fig. 2, corresponding to the end of helix 2 and beginning of the subsequent loop. Glutamic acid E77 is displayed on the structure for reference. **Top:** AA alignment next to a ribbon diagram of PDB entry 2JD7:A (Tatur et al. 2007), both colored with the ClustalX color scheme (Jeanmougin et al. 1998). **Bottom:** 3Di alignment using Jalview’s gecos-Sunset color scheme.

### Model selection

AA data are typically best fit by Blosum62 or LG, standard matrices for highly diverged AA sequences, or by EX3+FU, a mixture model containing three matrices modeling different transition rates and AA frequencies for exposed, intermediate, and buried sites (Le et al. 2008). Interestingly, LG is selected only for AA data aligned by a structure-free method. All 3Di alignments are best-fit by the 3Di substitution matrix with empirical base frequencies, with minor variations in the treatment of among-site rate variation (G4, R3-R5) (Table 1, Supporting Information).

**Table 1.**
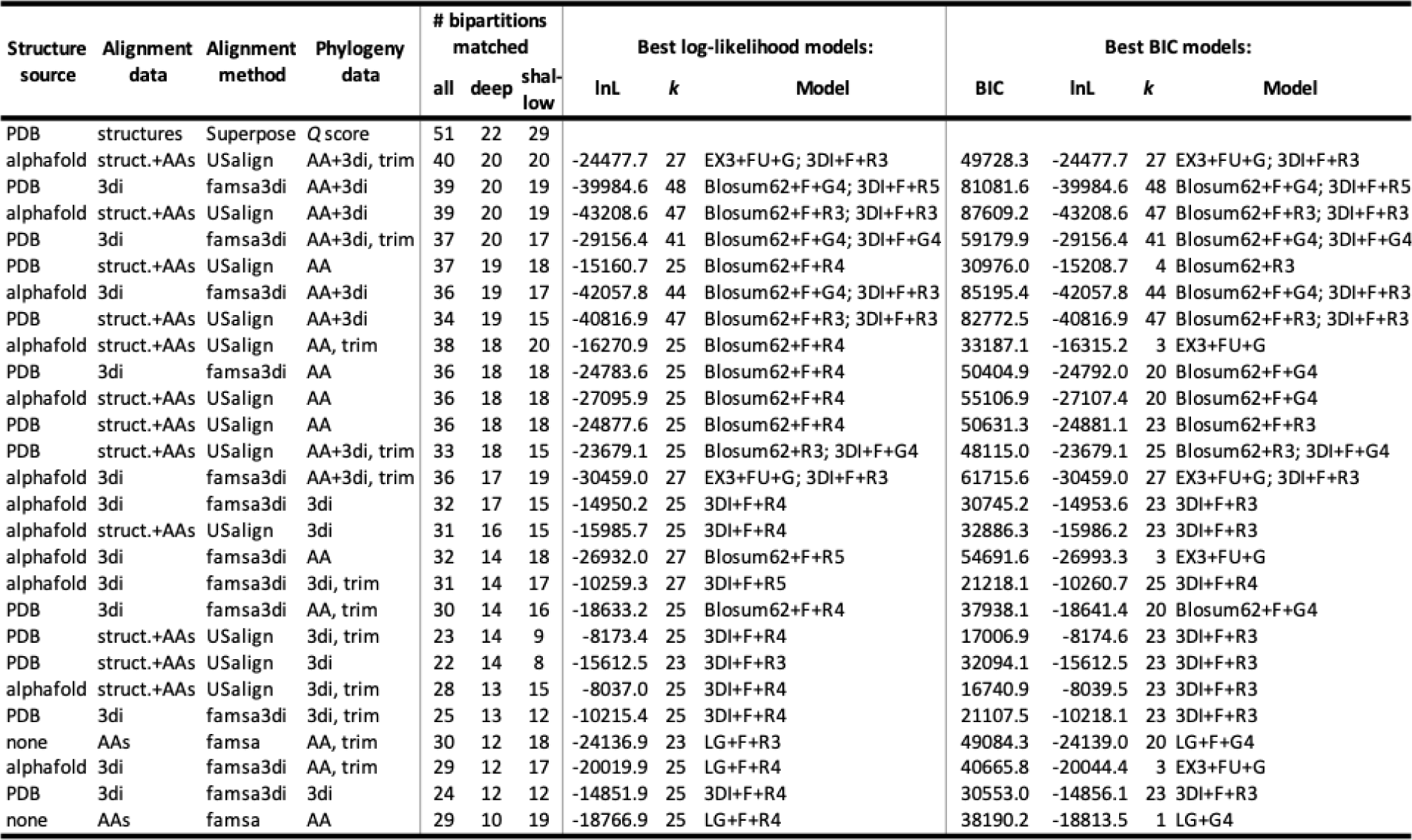
Best-fit substitution models for IQtree runs, according to log-likelihood (lnL) and BIC. IQtree inferences were conducted using the BIC-best model. Runs are sorted by deep & shallow similarity to the M20 tree (first line). Results are shown only for IQtree runs using the custom model list (with partitioning for AA+3Di datasets) which included the 3Di substitution matrix. *k* represents the number of free parameters in the substitution model; all runs also included 103 free branch-length parameters in the BIC calculation. Note: lnL & BIC values can only be compared within lines, not between them, as they are only comparable for identical alignments.

Standard AA models fit best to AA data, with comparatively small differences between Blosum62, LG, and EX3. A much worse fit is the Poisson model, a Jukes-Cantor-style model which assumes that transition rates are equal between all residues. Finally, the least fitting model for AA data is the 3Di rate matrix. On the other hand, for 3Di data, the pattern is reversed, where the 3Di model dominates, followed by Poisson and then the AA models (Table 2).

**Table 2.**
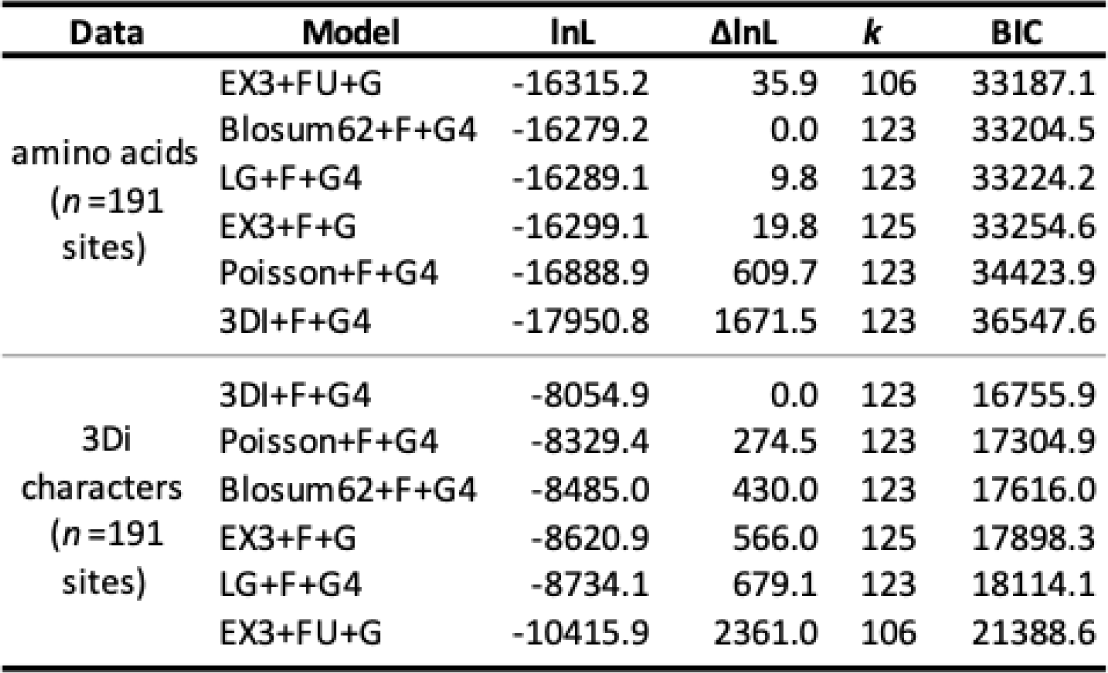
Illustrative models compared on AA and 3Di datasets. Alignment source: USalign on AlphaFold structures+AAs, trimmed sites with <35% data; phylogeny estimated with IQtree on custom model list including 3DI substitution matrix.

### Phylogenetic analyses

IQtree custom-model runs by the number of deep and shallow bipartitions that match the reference M20 tree are shown in Table 1. Results for all 60 ML runs are shown in Figure 4. Our results indicate that runs using both AA+3Di do best at matching the M20 tree topology, although with substantial overlap with structure-aligned AA runs. 3Di-only runs are consistently lower ranked.

**Figure 4.**
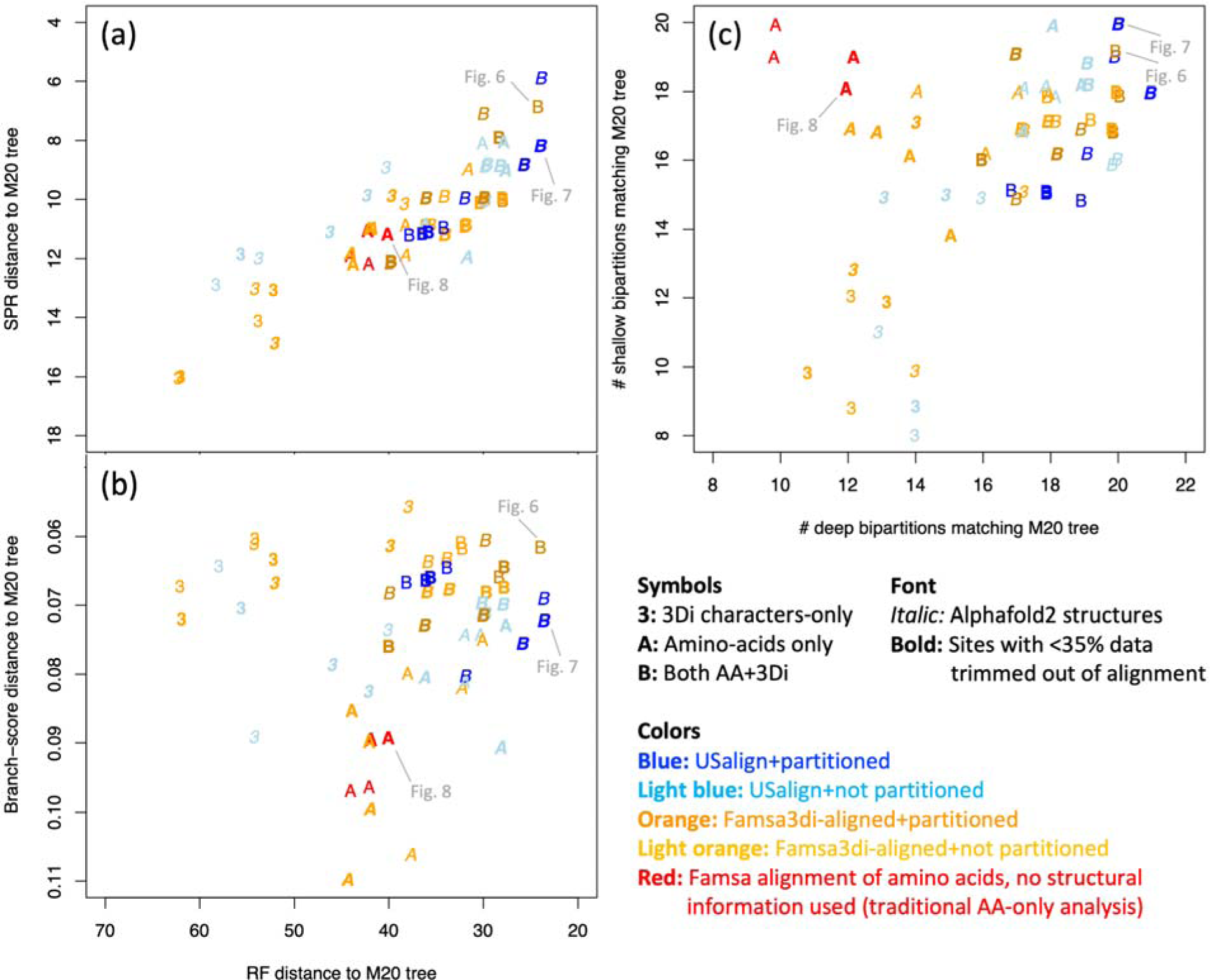
Similarity of all 60 IQtree runs to the Malik et al. (2020) structural-distances NJ tree (M20) for the ferritin-like superfamily. **(a)** Robinson-Foulds (RF) distance vs. Subtree Pruning-Regrafting (SPR) distance. SPR approximates the number of edits required to match the M20 topology. The closest matches to M20 on both metrics tend to be AA+3Di analyses with partitioning. AA-only and 3Di-only analyses tend to fare worse. **(b)** RF distance vs. branch-score distance, which takes branch lengths into account, unlike RF and SPR. 3Di-only match M20 well on the branch-score measure, but poorly on RF; however, AA+3Di performs well on both. **(c)** Number of M20 deep bipartitions matched vs. shallow bipartitions matched. “Deep” bipartitions are those annotated by Malik et al. to indicate major well-recognised categories, and groupings of those. Shallow bipartitions are all others, e.g. groups of 2-5 structures near the tips of the unrooted tree. AA-only runs retrieved shallow bipartitions but not deep ones; 3Di-only was weak on both, but AA+3Di retrieved both shallow and deep. *Note:* Points have been jittered for visibility.

Phylogenetic analyses using 3Di-only tend to be competitive in branch score difference. Structure-aligned AA-only runs are somewhat competitive in RF distance; only both AA+3Di runs are competitive in both (Figure 4b).

Amino-acid-only (structure-free) analyses are competitive for resolving shallow bipartitions, but are systematically poorer for deep bipartitions. 3Di-only analyses are poor for both. However, in combined analyses, the AA and 3Di data together seem to mutually compensate for the weaknesses of each individually and produce the best matching of both shallow and deep bipartitions (Figure 4c).

Also of interest are cases where differences in method do not seem to produce major differences in performance. It appears that PDB and AlphaFold structural data are equivalent in terms of phylogenetic results matching the M20 tree. Trimming, or not, also does not appear to make a major difference. Model partitioning did not seem to produce a consistent difference in terms of tree similarity to the M20 tree. However, lack of partitioning produces higher bootstrap supports. When bootstrap percent scores are summed across the tree for all 60 models (see SI), all 12 of the AA+3Di runs without partitioning were ranked in the top 15 (sum of bootstraps: 4219-4374). The next 16 runs contain 13 AA+3Di partitioned runs and 3 structure-aligned AA-only runs (sum of bootstraps: 4091-4216). The following 15 runs are all AA-only (3815-4087), with structure-free AA runs representing the last 4 (3815-3924), and the final 14 runs are all 3Di-only (3471-3799). While declining bootstraps for the noisier AA-only and 3Di-only datasets accords with expectations, elevated bootstraps for non-partitioned AA+3Di datasets probably represents artificial elevation of confidence due to a single model poorly fitting two very different partitions.

The nonmetric multidimensional scaling (NMDS) analysis of the RF distances between all ML trees and the M20 tree (“+”) is shown in Figure 5. An NMDS stress value of 0.12 indicates that AA-only and 3Di-only runs clearly explore different parts of tree space and supports the contention that they are not redundant data sources. The structure-free AA-only runs (Figure 5, red “A”) cluster apart from the structure-aligned AA-only runs, suggesting that AA alignments based on structure may pick up phylogenetic signal that structure-free AA alignments miss. 3Di-only analyses are much more scattered than other runs, suggesting larger variation in inferred tree topology, perhaps due to the fact that some 3Di runs were using the default IQtree model list, which does not include a 3Di-specific model. The AA+3Di runs (Figure 5,“B”) appear to combine both signals and cluster more tightly than AA-only or 3Di-only runs. The NMDS plot also indicates a difference between alignment methods. USalign-based AA and 3Di alignments (Figure 5, light blue “A” and “3”), all tend to be closer to the M20 tree than famsa3di-based alignments (light orange). This may indicate that USalign, by simultaneously aligning AAs and the 3D backbone of the structures, without considering 3Di states, is more directly capturing structural distance information that is also used by the M20 tree. However, this difference between USalign and famsa3di methods appears to be much reduced in the AA+3Di analyses (Figure 5, “B”).

**Fig. 5.**
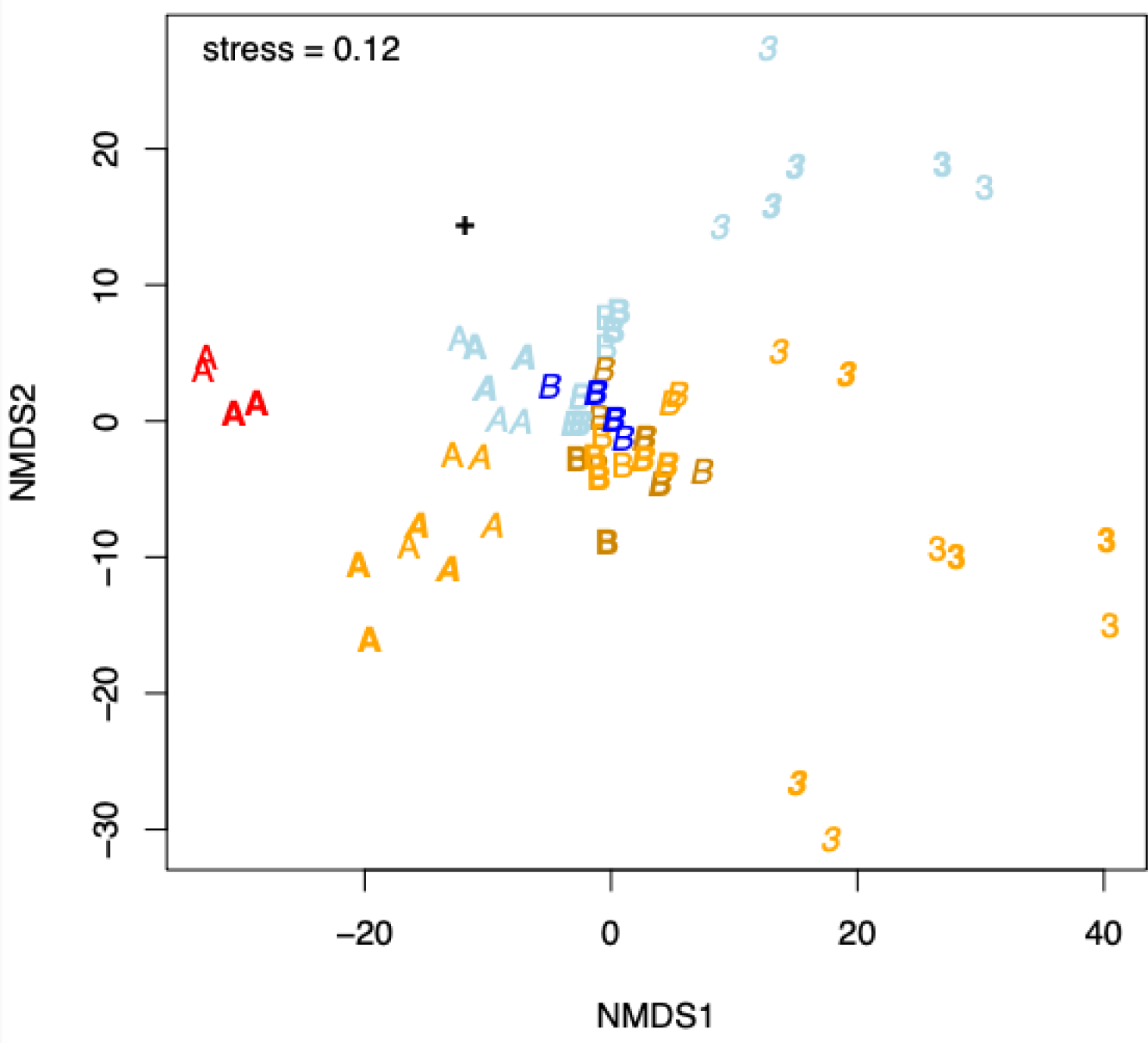
Nonmetric Multidimensional Scaling (NMDS) representation of the 61x61 matrix of RF distances between all ML trees (60 IQtree runs + the M20 tree). Values <0.1 are typically taken to indicate “excellent,” and <0.2 “good,” 2-dimensional approximations of the original distances (Clarke 1993). Legend as in Figure 4. M20 tree = **+**. The trees using both AA+3Di data (“B”) show a merger of both signals and cluster tightly, compared to the variability between the AA-only or (particularly) 3Di-only trees.

For PDB structural data, the phylogenetic tree that best matched the M20 tree (Figure 6) was generated using famsa3di on 3Di characters generated from the PDB structures. The BIC best-fit models for the AA and 3Di partitions were Blosum62+F+G4 and 3DI+F+R5. This tree shared 19 deep and 20 shallow bipartitions with the M20 tree. Similarity is apparent in both the unrooted (top) and midpoint rooted/re-rooted (bottom) depictions (Figure 6). The most notable topology differences are in the placement of two structures on long, isolated branches in M20: 2fzf_A, a *Pyrococcus furiosus* rubrerythrin domain-containing protein, and 1z60_M, a ferritin chain from the cabbage looper butterfly *Trichoplusia ni* which interacts with 1z60_A from the same species. In the PDB AA+3Di tree, 2fzf_A groups with the other rubrerythrins (albeit with low support, 57%), and 1z60_M is sister to 1z60_A (93% support), suggesting within-structure gene duplication. These are both highly plausible placements, improving on the M20 tree.

**Figure 6.**
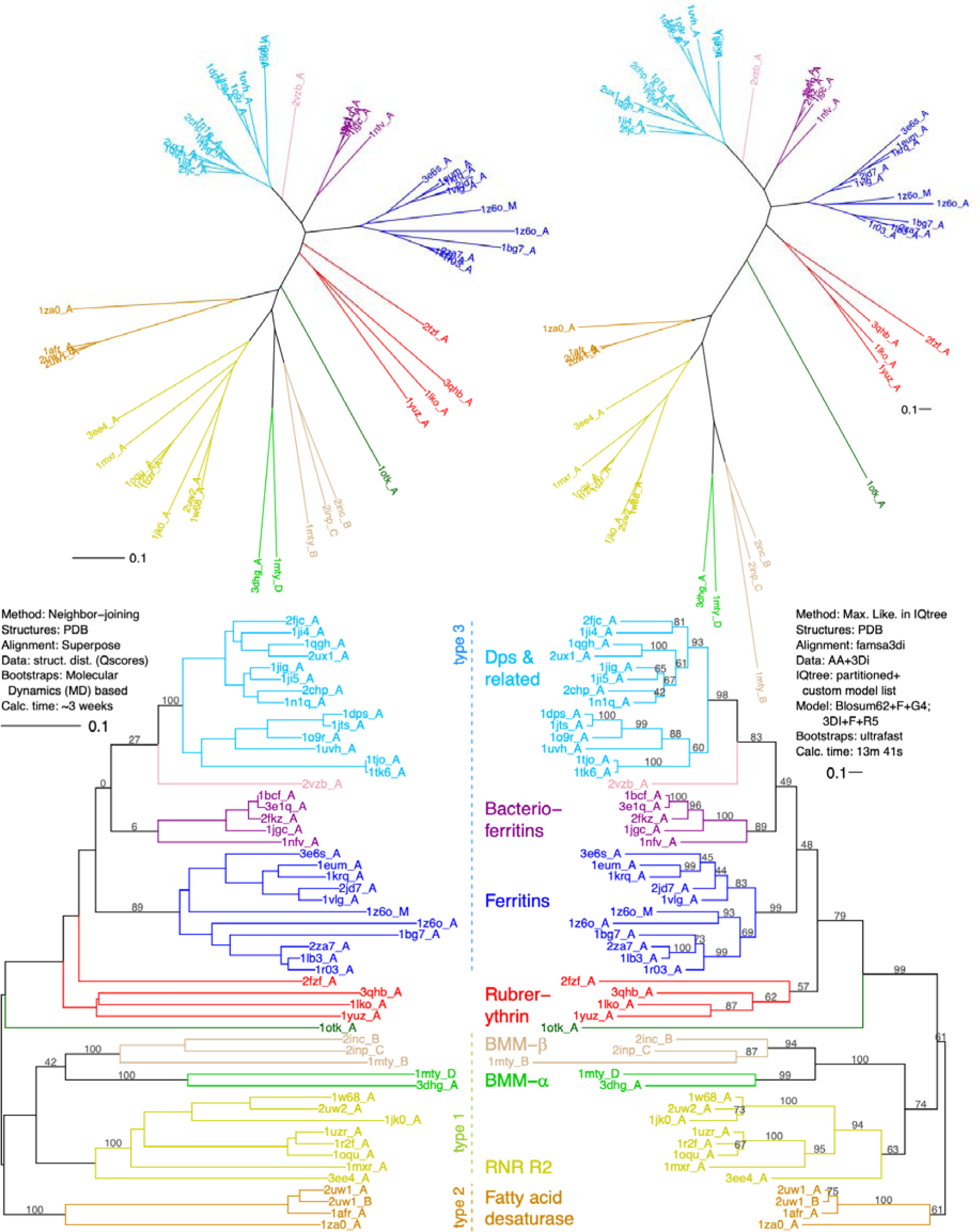
Comparison of trees using PDB structures. **Left:** M20 structural distances tree, with MD-based bootstraps where available in Malik et al. 2020. **Right:** the closest-match PDB-based IQtree analysis using AAs+3Dis from PDB structural data. This tree shares 20 deep and 19 shallow bipartitions with M20. **Top row:** unrooted trees. **Bottom:** arbitrary rooting for visual comparison. Branch length units for the M20 tree are *Q*_score_-based distances; for the ML tree they are expected substitutions per site. The types refer to dimer geometries, likely slowly evolving structural characters (Lundin et al. 2012).

The best-match IQtree analysis overall (Figure 7) used AlphaFold structure predictions, jointly aligned structures and AAs using USalign, with Alphafold-derived 3Di characters then mapped to the aligned AA positions. The alignment was trimmed, and the BIC best-fit models for the AA and 3Di partitions were EX3+FU+G and 3DI+F+R3. This tree shared 20 deep and 20 shallow bipartitions with the M20 tree. This analysis also groups 2fzf_A with other rubrerythrins (similar low support, 60%) and 1z60_M with 1z60_A (70% support). The other major difference is that the dimer geometry type 2 group of fatty acid desaturases is nested within the type 1 group, as sister to RNR R2, although with low support (69%).

**Figure 7.**
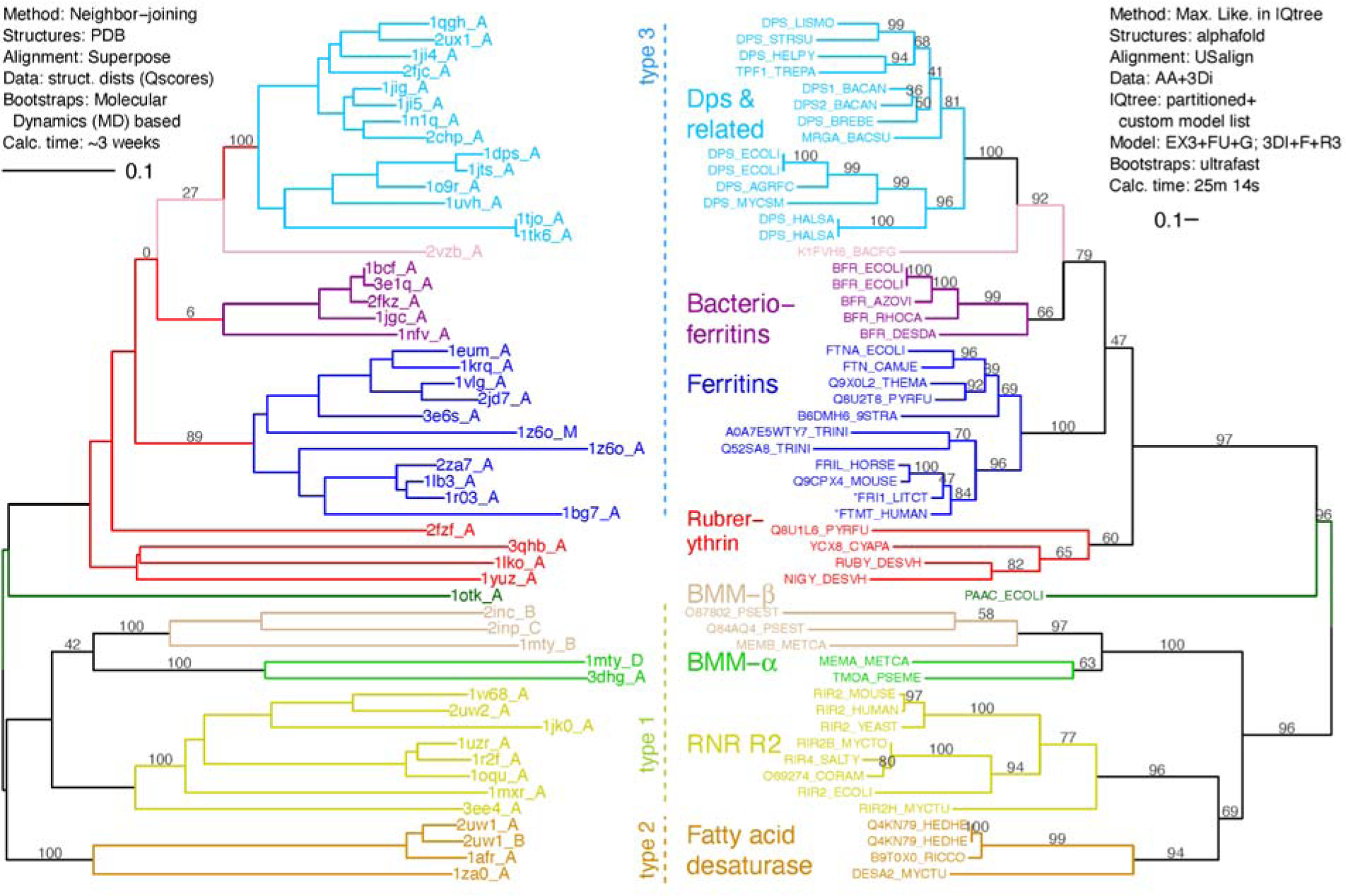
AlphaFold-based ML AA+3Di tree (right) compared to the M20 structural distances tree (left). This ML run was the closest overall topology match to the M20 tree, and shares 20 deep and 20 shallow bipartitions with M20. Tip labels are Uniprot IDs for which AlphaFold predictions were downloaded. Each Uniprot label on the right represents the closest AA match to the corresponding PDB structure on the left, with the exception of two tips marked with asterisks; here, FRI1_LITCT corresponds to 1bh7_A, and FTMT_HUMAN corresponds to 1r03_A. The types refer to dimer geometries, likely slowly evolving structural characters (Lundin et al. 2012).

The resulting topology from a “standard”, sequence-only, structure-free alignment (Figure 8) shares 19 shallow bipartitions with M20, but only 12 deep ones. Severe long-branch attraction effects are evident; instead of 2fzf_A grouping with the rubrerythrins and 1z60_A with 1z60_M and other ferritins, they instead form a group of oddballs along with the long-branched 1otk_A. The BMM-β group breaks apart and renders fatty acid desaturases and ferritins paraphyletic.

**Figure 8.**
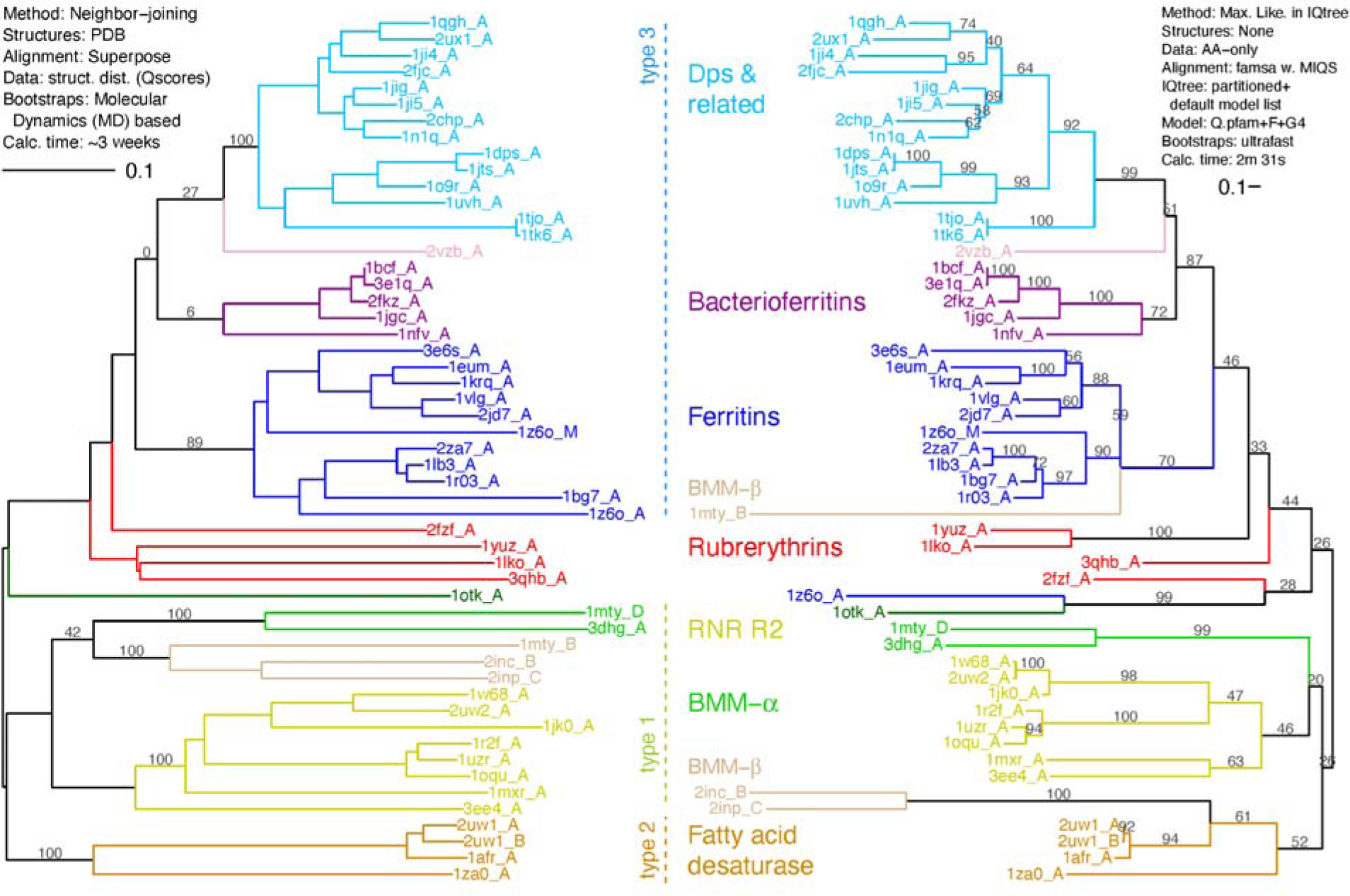
Structure, AA-only ML tree (right) compared to the M20 structural distances tree (left). ML tree is from the closest match “standard” AA-only analysis. The tree shares 12 deep and 19 shallow bipartitions with M20. The types refer to dimer geometries, likely slowly evolving structural characters (Lundin et al. 2012).

In terms of computational run-time, these results are a dramatic improvement. The Malik et al. 2020 MD-based bootstrapping of ferritins took approximately 3 weeks of wall-clock time to run, even making use of parallel processing, as each of the 53 structures had simulate 100 nanoseconds (ns) of conformational change with Molecular Dynamics. Even longer simulations (>500 ns) would be preferred in a new analysis, which would imply a 5x increase in runtime. Although this may be ameliorated with newer technology and greater cluster resources when available. On the other hand, the focal PDB and AlphaFold AA+3Di IQtree runs, complete with partitioning, automated model selection, and 1000 ultrafast bootstraps, took 13.6 and 25.25 minutes on a standard desktop computer. The latest approach makes structural phylogenetics more accessible since it can be performed with small computational resources.

## Discussion

Although protein structural phylogenetics has been explored for decades, its use has been limited by the availability of structural data, the lack of ready-to-use inference methods other than distance-based neighbor-joining, and the challenges of integrating distance-based methods into the model-based framework of modern statistical phylogenetics. Highly desirable tools like parametric model fitting, statistical model comparison, Maximum Likelihood and Bayesian inference, statistical measurement of uncertainty through bootstrapping, posterior probabilities or other techniques, integration of multiple data sources in a unified statistical framework, and simulation-based *in silico* tests of inference methods, all remained challenging in a distance-based framework.

The combination of two developments, namely AlphaFold and the Foldseek tertiary interaction encoder, raises the prospect that this barrier has suddenly fallen, and the way is open for the emergence of model-based structural phylogenetics. Our study represents a proof-of-concept: a complete pipeline stepping from amino acid sequences, to AlphaFold structures, to structure-informed alignments, to 3Di character datasets that appear to be well-behaved, and helpful, in statistical model selection (Table 2) and phylogenetic inference (Figs. 4-8).

In the case of the ferritin-like superfamily, the application of this pipeline produced estimated phylogenies, with formal ultrafast bootstrapping estimates of uncertainty, in minutes instead of the weeks required in the previous state-of-the-art analysis of Malik et al. (2020). Across the different IQtree run conditions, the phylogenies that were the best topological matches to the Malik et al. (2020) were the ones that one might predict *a priori* to be phylogenetic best-practice (Table 1): combining both AA and 3Di alignments, using structure to inform the alignment, using a custom model list that included a 3DI-specific substitution model, and using statistical model comparison on the partitioned dataset to find the best-fit model separately for AA and 3Di characters. Combining AA+3Di characters seemed to ameliorate long-branch attraction artifacts found in AA-only analyses (Figs. 6-8), and also produced closer matching to the Malik et al. tree on the branch-score distance (Fig. 4b). Probably these are related phenomena, suggesting that AA+3Di analyses are less affected by saturation of sequence divergence on deep branches. All of these results are just what would be expected when a set of characters even more conserved than amino acids are added to an amino acid dataset.

In summary, our results suggest that the inferred structure-informed phylogenies are comparable, and perhaps superior, to a previously-published phylogeny of the ferritin-like superfamily using structural distances. The addition of 3Di characters to ML phylogenetic inference is appealing because of the approach’s ease-of-use, computational speed, and potential to add highly-conserved, slowly-evolving characters just where they are typically needed: deep in phylogenies, where AA sequence similarity is highly decayed. Therefore, it is tempting to suggest that this approach can, and should, be tried on almost every tough problem in protein phylogenetics. While we do encourage researchers to explore and build on this approach, our study highlights a number of considerations that should be kept in mind.

In the case of the ferritin-like superfamily, AlphaFold-predicted structures have high confidence scores (see Supporting Information) and performed equally well to PDB structures in our analyses. However, while AlphaFold has been a huge step forward in structural prediction, some limitations have become evident. AlphaFold predictions will tend to be more accurate on structures more similar to its (admittedly vast) training set. Protein types that have fewer solved high-resolution structures (membrane-embedded proteins or those with no stable fold) will have more uncertain AlphaFold predictions and therefore will presumably have noisier 3Di characters. Intrinsically disordered proteins/regions/loops are also poorly predicted by AlphaFold (Ruff & Pappu 2021; Bertoline et al. 2023), although such regions are likely to be less conserved anyway. The AlphaFold database also has the limitation of holding only structural predictions of monomers. Interactions with cofactors, chaperones, and/or within multimers will all potentially change a predicted structure (Perrakis & Sixma 2021) and thus any extracted characters. Interactions can be modeled with AlphaFold, but for the moment this is a bespoke process rather than a precalculated database.

### Unresolved questions about 3Di characters

The prospect of estimating phylogenies with the assistance of a 3Di character dataset that might change based on how protein structures are modeled and predicted may give pause to many molecular phylogeneticists who are used to sequence data being fixed and highly confident (leaving aside cases of sequencing error and contamination). A broader focus on the kinds of characters that have been used in phylogenetics generally might help bridge this gap. In organismal biology and paleontology, sources of characters have included morphology of bones as well as soft tissue, ultrastructure, characters observable only in a particular sex (antlers) or embryonic developmental stage (post-anal tail), characters that are revealed only after substantial processing or imaging of a specimen (biochemical fossils, internal bone structure), or even ephemeral events like bird songs or stereotyped mating behaviors. Just like 3Di characters, many of these other data sources produce characters that are dependent on a particular ascertainment method. As long as an ascertainment method can be described and replicated by other researchers, it is a potentially valid data source, and its utility for phylogenetics becomes an empirical question. For a novel data source, tree structure in character data, phylogenetic congruence with other evidence, and reliability of character coding under perturbation of the input material are all standard research questions that can be answered statistically (Matzke 2015).

A case can be made that 3Di states are actually easier to accept as phylogenetic characters than routinely accepted data sources like discrete morphological characters. Morphologists often encounter difficult questions such as: How should continuous data be discretized? Is a character state not observed because it is not present in the organism, or is it missing due to specimen incompleteness, decay or damage (taphonomy; Sansom et al. 2010), the specimen being a juvenile, or population-level, environmental, or random variation in development? Sometimes there is an element of subjectivity in the judgment calls made in morphological character coding that can be difficult to completely remove. In comparison, 3Di characters have no such difficulties. For a given experimental or AlphaFold-predicted structure file, the output 3Di characters are a deterministic result that is easily repeatable by any researcher. It is, of course, likely that future improvements in structural prediction and in tertiary-interaction encoding will produce variant datasets, but these will also be replicable and subject to statistical tests for phylogenetic signal. Replicability will require that any future iterations on structural character encoding schemes are given clear version numbers that are reliably reported by researchers, and also that old versions of encoders are kept available in software packages for backwards comparability.

Despite the good arguments for the validity of 3Di states as characters, many questions do need further research. We caution that it remains unclear how sensitive the 3Di encoding is to small perturbations in the protein structure, such as indels, or minor errors made during protein structure solving or prediction. As 3Di characters are based on minimum distances between residues, small changes in one region of the protein might sometimes induce rippling effects that perturb the 3Di characters elsewhere, potentially disrupting the phylogenetic signal. However, the success of Foldseek for remote homology search, and our results showing high conservation over vast evolutionary distances (Fig. 2), as well as congruence between PDB- and AlphaFold-derived phylogenies (Figs. 6-7), suggest that 3Di characters are likely to be fairly robust to perturbations. We would, however, caution that it may be hazardous to “mix and match” structures from different sources: for example, a dataset consisting of some PDB experimental structures, some AlphaFold-database monomer predictions, and some bespoke AlphaFold predictions of chains in a multimeric conformation might result in systematic biases that end up grouping structures by method rather than by history. (On the other hand, it is also possible that in some structures, the evolutionary-conserved tertiary structure is best observed/predicted in the multimeric state.)

### Other caveats

Model-based phylogenetic estimation using discretized structural characters is novel, and several classic questions in statistical phylogenetics need to be revisited when the methods are applied to 3Di characters. First, programs like IQtree employ the standard phylogenetic assumption of site independence in their likelihood calculations. This assumption allows the log-likelihood of each site in the alignment to be calculated independently; summing across sites allows rapid calculation of the total log-likelihood. The very definition of 3Di characters as descriptors of tertiary interactions between pairs of residues identified as closest neighbors arguably violates this assumption, although the question is complex as neighbor relationships do not have to be symmetric. Violations of the site independence assumption in other data are already well-known; for example, in coevolution of AA sites (Korber et al. 1993), or in co-variation of morphological characters due to a shared developmental pathway or convergence on an ecological role. These are usually not considered fatal for phylogenetic inference, but co-dependence may be even greater for 3Di characters. Failure to account for site correlation may lead to spurious results by placing overconfidence in the wrong tree (i.e., inflated bootstrap or posterior supports), as information is being double counted (Nasrallah et al. 2011).

IQtree also assumes independence between AA and 3Di sequences. While inspection of AA and 3Di alignments proves that they are far from identical, and clearly have some nonoverlapping grouping information, ideally any nonindependence between the datasets could be measured and corrected for.

Another limitation concerns phylogenetic information that 3Di characters may be missing. 3Dis describe the tertiary relationships between segments of a structure’s backbone. This approach does not directly capture structural changes that only affect side chains. In addition, the dihedral angles that directly describe the changes in backbone orientation from one residue to the next are not directly captured by 3Di. Whether these additional kinds of structure-derived datasets could be combined with AAs and 3Di in a total-evidence analysis, or this would only exacerbate the non-independence problem, is an open question.

### Future outlook

Alongside the methodological questions above, 3Di characters present phylogeneticists with a number of additional research opportunities. Just as DNA models advanced from Jukes-Cantor to GTR and codon models, and just as AA models advanced from PAM and BLOSUM matrices to LG and mixture models, character-based structural phylogenetics is likely to advance from the Poisson and 3DI matrices to matrices optimized for different kinds of proteins and protein regions. It is likely that the frequencies of different 3Di character states change between alpha-helix and beta-sheet regions of proteins, so an obvious first step would be to explore the potential of partitioning a 3Di dataset on a protein family containing both (the ferritin fold is largely alpha-helical).

The speed of ML analyses including 3Di structural information also means that Bayesian structural phylogenetics is clearly feasible. This would allow the incorporation of additional modeling flexibility and sources of information, including relaxed clocks, absolute and relative (Shih and Matzke, 2013) time constraints, joint inference of substitution model and tree, and potentially even uncertainty in structural prediction and the derived 3Di characters.

Finally, while *a priori* considerations and our proof-of-concept study strongly suggest that the addition of 3Di characters to AA datasets will improve phylogenetic inference, especially deep in the twilight zone, the ideal study would test this hypothesis in a situation where the true phylogenetic tree is known. While an *in silico* test would be relatively easy to perform under the model assumed by IQtree, where 3Dis are just another 20-state character where every site evolves independently under the 3DI rate matrix, this form of simulation-inference experiment is only of limited interest as it does not model the evolution of the 3-dimensional protein structure. While models of structural evolution are beginning to be developed, a realistically complex forward model of protein structural evolution remains a major challenge. An alternative source of known phylogenies could be experiments, but this would require a system where AA sequence identity could decay from 100% to <30% in the course of the experiment, all while retaining substantial structural similarity. It may be that this could be achieved in a rapidly-evolving virus system, although whether this would imitate protein structural evolution at billion-year timescales could be debated.

Apart from methodological investigations, the major research opportunity presented by character-based structural phylogenetics is a potential revolution in the study of deep phylogenetics questions stretching back before the Last Universal Common Ancestor. The origin and evolution of protein families and fundamental cellular machinery can potentially be addressed or re-addressed with more data and more confidence. The deepest, hardest to resolve relationships – the relationships of the bacterial phyla, of archaea and eukaryotes, and of different classes of viruses, may all become more accessible. However, it may also be that structural characters will be useful even for improving the resolution of shallower relationships, as reported by Moi et al. (2023). We think it likely that structural phylogenetics will soon move from revolutionary to routine.

## Acknowledgements and Funding

This collaboration emerged from the inaugural Australasian Structural Phylogenetics Meeting (ASPM2023) that took place at the University of Auckland on October 24th and 25th, 2023. The meeting was supported by the Centre for Computational Evolution (CCE) and the Centre for Cellular and Molecular Physiology (CMP), School of Biological Sciences at the University of Auckland. CPL, MABB and NJM are supported by Human Frontier Science Program Grant No. RGY0072/2021. NJM was additionally supported by NZ RSTA grants 21-UOA-040 & 18-UOA-034. JD is supported by the Alfred P. Sloan Foundation Matter-to-Life program Grant number G2021-16944. DBA is supported by the investigator grant from the National Health and Medical Research Council (NHMRC) of Australia [GNT1174405] and the Victorian government Operational Infrastructure Support program. DL is supported by grants from Vetenskapsrådet 2021-03992 and Knut och Alice Wallenbergs stiftelse 2019.0436 to Martin Högbom. MSF is supported by Marsden Fast Start #3726031. MS acknowledges the support by the National Research Foundation of Korea, grants [2020M3-A9G7-103933, 2021-R1C1-C102065, 2021-M3A9-I4021220], Samsung DS research fund and the Creative-Pioneering Researchers Program through Seoul National University.

## Supporting Information

Data files, analysis scripts, results tables, and alignment and IQtree run inputs and outputs are available at https://github.com/nmatzke/3diphy or as the 191 MB zipfile *3diphy.zip*. The README file in the main directory points to key files/analyses mentioned in the main text: *_00_README_SuppInf_files_v2.txt*.

## Notes

https://github.com/nmatzke/3diphy

### Competing Interest Statement

The authors have declared no competing interest.

### Summary of Updates

Updated Figure 3b to correct coloring of 3Di characters on ribbon structure. Added as coauthor: Hyunbin Kim from the Steinegger Lab for his contribution on color schemes for 3Di characters.

