## Supplementary material for "Tertiary-interaction characters enable fast, model-based structural phylogenetics beyond the twilight zone": 02_add_ferritins_split_into_chains_v1.R

```
# Original 36 ferritins
cp *A.pdb /GitHub/str2phy/ex/ferritins_M20

# Manually delete 2 that Malik et al. 2020 excluded
# 2cwl_A.pdb	MalikExcluded	
# 2ib0_A.pdb	MalikExcluded


# Get the rest:

Batch download from:
https://www.rcsb.org/downloads

1bg7_A
1n1q_A
1qgh_A
1r03_A
1tjo_A
1tk6_A
1z6o_M
2fjc_A
2vzb_A
3e6s_A
1mxr_A
1oqu_A
3ee4_A
1mty_B
1mty_D
2inc_B
2inp_C
2uw1_B
1otk_A


cd /GitHub/str2phy/ex/ferritins_M20/rawdata/z_full_structures
gunzip *.gz


#######################################################
# Then, extract specific chains with R package bio3d
#######################################################

library(bio3d)

wd = "/GitHub/str2phy/ex/ferritins_M20/rawdata/z_full_structures"
setwd(wd)

output_path = "/GitHub/str2phy/ex/ferritins_M20/rawdata"

pdb.files = c("1bg7.pdb", 
"1n1q.pdb", 
"1qgh.pdb", 
"1r03.pdb", 
"1tjo.pdb", 
"1tk6.pdb", 
"1z6o.pdb", 
"2fjc.pdb", 
"2vzb.pdb", 
"3e6s.pdb", 
"1mxr.pdb", 
"1oqu.pdb", 
"3ee4.pdb", 
"1mty.pdb", 
"1mty.pdb", 
"2inc.pdb", 
"2inp.pdb", 
"2uw1.pdb", 
"1otk.pdb")

ids = c("1bg7_A", 
"1n1q_A", 
"1qgh_A", 
"1r03_A", 
"1tjo_A", 
"1tk6_A", 
"1z6o_M", 
"2fjc_A", 
"2vzb_A", 
"3e6s_A", 
"1mxr_A", 
"1oqu_A", 
"3ee4_A", 
"1mty_B", 
"1mty_D", 
"2inc_B", 
"2inp_C", 
"2uw1_B", 
"1otk_A")


chain_split_fns = bio3d::pdbsplit(pdb.files=pdb.files, ids=ids, path=output_path)
chain_split_fns


# Count the number of PDB chain files in directory, in Terminal
cd /GitHub/str2phy/ex/ferritins_M20/rawdata/
ls -1 *.pdb | wc -l

# 53 files
```
