## Supplementary material for "Tertiary-interaction characters enable fast, model-based structural phylogenetics beyond the twilight zone": put_3di_into_FAMSA.R

```
#######################################################
# Convert a 3di substitution matrix into a format
# suitable for inserting into FAMSA source code
# (FAMSA source code is at: https://github.com/refresh-bio/FAMSA/releases )
#######################################################

wd = "/GitHub/str2phy/substmat/"
setwd(wd)

# Load the Foldseek 3di cost matrix
di3_fn = "mat3di.out"
di3 = read.table(di3_fn, header=TRUE, sep="")
di3

# Find the minimum value
minval = min(di3)
minval
# -17

# Load the FAMSA MIQS cost matrix
# (source: Yamada, K. & Tomii, K. Revisiting amino acid substitution matrices for identifying distantly related proteins. Bioinformatics 30, 317–325 (2014). ) 
miqs_fn = "FAMSA_MIQS.txt"
miqs = read.table(miqs_fn, header=TRUE, sep="")
colnames(miqs)[length(colnames(miqs))] = "star" # The "*" converts to "V.". Rename "star"
rownames(miqs) = colnames(miqs)
miqs

# Note: the -6.1, used for "characters" "B	Z	X	*", 
# is just the MINIMUM value observed in the main
# 20x20 part of MIQS.

# Convert the 3di matrix to FAMSA ordering and format

# Reorder the 3di
di3_to_famsa_order = match(x=names(miqs)[1:20], table=names(di3)[1:20])
di3_to_famsa_order

tmp = di3[di3_to_famsa_order, di3_to_famsa_order]

# Yes they match
names(miqs)
names(miqs)[1:20]
names(tmp)
rownames(tmp)

# Copy reordered 3di into miqs
di3_famsa = miqs
di3_famsa[1:20, 1:20] = tmp
di3_famsa

di3_famsa[,21:24] = minval
di3_famsa[21:24,] = minval
di3_famsa

#######################################################
# Write out to text for insertion into famsa3di
#######################################################
for (i in 1:nrow(di3_famsa))
	{
	txt1 = paste0(di3_famsa[i,], collapse=", ")
	txt2 = paste0(" { ", txt1, "}, // ", rownames(di3_famsa)[i])
	txt2
	cat(txt2)
	cat("\n")
	}


# Compile as follows:
cd /Users/nickm/Downloads/FAMSA_3di-2.2.2/
make
mv famsa famsa3di
cp famsa3di /usr/local/bin
cp famsa3di /Applications/FAMSA-2.2.2/
cp famsa3di /Applications/FAMSA-2.2.2/
cp famsa3di /GitHub/str2phy/substmat/
```
