## Supplementary material for "Tertiary-interaction characters enable fast, model-based structural phylogenetics beyond the twilight zone": _00_README_SuppInf_files_v2.txt

=========================================
Supporting Information for Puente-Leilivre et al. 2023

These files are provided to promote:

* data accessibility
* future analyses of the same or expanded datasets
* to document the commands and scripts used to perform the analyses in this paper

They are, however, are not designed to be a "run from scratch" tutorial. Readers wishing to run the scripts may therefore require some prior experience with command-line, R, and phylogenetics programs, and the scripts will require some modifications (e.g., installing R packages, changing the "wd = " (working directory) lines).
=========================================


=========================================
Data files and script/command files, as used, in order, in the Main Text:
=========================================

..."53 PDB structures"

List of structure files:
01_ferritin_taxa_v1.xlsx

PDB files & derivatives:
ferritins_M20/rawdata

AlphaFold files & derivatives:
ferritins_M20af/rawdata


[Structures] "...clipped to the single chains used by Malik et al. using bio3d::pdbsplit"

02_add_ferritins_split_into_chains_v1.R


"The list of PDB chain files was processed to AA and 3Di FASTA files"

ferritins_M20/00_pdbs_to_3di_v1.R
ferritins_M20af/01_pdbs_to_3di_v1.R
Rsrc/str2phy_v1.R


"...when no 100% match was found, the closest-hits AlphaFold structure was chosen."

This occurred in 9/53 cases: 1bg7_A (152/153 AAs identical), 1dps_A (158/159), 1jts_A (153/155), 1mty_D (511/512), 1uzr_A (281/282), 2uw1_A (320/327), 2uw1_B (332/338), 2uw2_A (274/275), 2za7_A (169/170).

See:
ferritins_M20af/53_ferritin_alphafold_hits_v2.xlsx


"See SI for a breakdown of pLDDT summaries."

ferritins_M20af/confidence.tsv
ferritins_M20af/plotConfidence.R


"...a 3Di alignment can be transferred position-by-position to unaligned AA characters and vice versa, a procedure we implemented in the custom R function align_3dis_to_AAs."


Example:
ferritins_M20/__cmds_ferritins_M20_famsa3di_USaln_then_trim_iqtree_v1.R
Rsrc/str2phy_v1.R


"The custom list included a custom 3Di rate matrix, which we calculated from Foldseek's 3Di substitution cost matrix..."

Starting with: 
3DI_substmat/mat3di.out from Foldseek

Conversion:
3DI_substmat/converting_AA_matrices_v4.xlsx

Result (the same 3DI transition matrix in 3 different files for different IQtree commands):
3DI_substmat/3di_substmat.txt
3DI_substmat/3DI
3DI_substmat/3DI.nexus


"All IQtree runs were conducted with 1000 ultrafast bootstraps (UFBoot)..."

These subdirectories contain the different alignment & IQtree runs:

# PDB-structure based
ferritins_M20/01_FAMSA_AAs_aln_v1 (only AAs used, structure ignored)
ferritins_M20/02_FAMSA_3dis_aln_v1 (famsa3di structural alignment of 3Dis, AAs mapped to this)
ferritins_M20/03_USaln_v1 (USalign joint alignment of 3D structure+AAs, 3Dis mapped to this)

# The same for AlphaFold structures
ferritins_M20/01_FAMSA_AAs_aln_v1 (only AAs used, structure ignored)
ferritins_M20/02_FAMSA_3dis_aln_v1 (famsa3di structural alignment of 3Dis, AAs mapped to this)
ferritins_M20/03_USaln_v1 (USalign joint alignment of 3D structure+AAs, 3Dis mapped to this)

# An example alignment / trimming / IQtree script 
# (only part is R, the rest is Terminal command-line commands)
ferritins_M20/01_FAMSA_AAs_aln_v1/00_alignIQtree_AAs_wFAMSA_v1.R


"Using custom R scripts (SI) we extracted from each IQtree run the best fit models and corresponding log-likelihood, AIC, AICc, BIC..."

Rsrc/parsing_iqtree_file_v1.R
ferritins_M20/00_plot_contrees_v3.R


"...FASTA files are available in SI."

For alignment commands and resulting FASTA alignment files:

# PDB-structure based
ferritins_M20/01_FAMSA_AAs_aln_v1/00_align
ferritins_M20/02_FAMSA_3dis_aln_v1/00_align
ferritins_M20/align/03_USalign_v1

# The same for AlphaFold structures
ferritins_M20af/01_FAMSA_AAs_aln_v1/00_align
ferritins_M20af/02_FAMSA_3dis_aln_v1/00_align
ferritins_M20af/03_USaln_v1/00_align


"Table 1 shows the best-fitting models for each IQtree run that used a custom model list (for all 60 runs, see SI)."

tree_comparison/60iqtree_runs_compared_v4_sumMatches_order.xlsx


"When bootstrap percent scores are summed across the tree for all 60 models (see SI)..."

tree_comparison/60iqtrees_sorted_by_sumBootstraps.xlsx


#######################################################
# Key files for phylogeny figures
#######################################################

# Malik et al. 2020 Figure 10b tree is here

# With M20 figure 10b labels:
ferritins_M20/tree_comparison/Malik_2020_Fig10b_FerritinData_distance_matrix_BIONJtree_M20root_wLabels.newick

# with M20's bootstraps
ferritins_M20/tree_comparison/Malik_2020_Fig10b_FerritinData_distance_matrix_BIONJtree_M20root_wBS.newick

# A table of the M20 tree, one line per node
ferritins_M20/tree_comparison/Malik_2020_Fig10b_FerritinData_distance_matrix_BIONJtree_M20root_trtable.txt


#  IQtree phylogeny for Figure 6 (best-match tree using PDB structures)
/ferritins_M20/02_FAMSA_3dis_aln_v1/iqtree_BothTrim35/partitions_v1.raxml.contree

#  IQtree phylogeny for Figure 7 (best-match tree using AlphaFold-predicted structures)
/ferritins_M20af/03_USaln_v1/iqtree_BothTrim35/partitions_v1.raxml.contree

#  IQtree phylogeny for Figure 8 (best-match tree using AA-only, structure-free analysis)
/ferritins_M20/01_FAMSA_AAs_aln_v1/iqtree_AAsTrim35_allmodels/53_ferritin_FAMSA_AAs_alnTrim.fasta.contree


=========================================
Note: the core command-line commands for converting a PDB or AlphaFold structure file to AA and 3Di FASTA entries are given at:

examples/pdb_to_3di/_README_str_to_3di_v2_WORKS.txt
=========================================
