## Supplementary material for "Tertiary-interaction characters enable fast, model-based structural phylogenetics beyond the twilight zone": 00_plot_contrees_v3.R

```
#######################################################
# Plot IQtree trees
#######################################################

library(ape)
library(BioGeoBEARS)
library(Quartet)   # for Quartet distances
library(gdata)		# for trim
library(stringr)		# for ?
library(stringi)		# for fancy string search/replace
library(phangorn)		# for treedist
library(psych)			# for psych::pairs.panels
wd = "/GitHub/str2phy/ex/ferritins_M20/iqtree/"
setwd(wd)

cmds='
cd /GitHub/str2phy/ex/ferritins_M20/iqtree/
find . -type f -name '*.tree'
' # END cmds

distance_trfn = "Malik_2020_Fig10b_FerritinData_distance_matrix_BIONJtree.newick"
distance_tr = read.tree(distance_trfn)


pdffn = "Malik_etal_2020_Fig10b_structural_dists_BIONJ_v1.pdf"
pdf(file=pdffn, width=12, height=12)

# Distance tree
tr = phytools::midpoint.root(distance_tr)
tr = ladderize(tr)
plot.phylo(tr, type="unrooted", lab4ut="axial", cex=0.65)
add.scale.bar()
title("Malik et al. 2020 structural distances tree with Splitstree:BIONJ")
dev.off()
cmdstr = paste0("open ", pdffn)
system(cmdstr)

alphafold_xls_order = c("1afr_A",
"1bcf_A",
"1bg7_A",
"1dps_A",
"1eum_A",
"1jgc_A",
"1ji4_A",
"1ji5_A",
"1jig_A",
"1jk0_A",
"1jts_A",
"1krq_A",
"1lb3_A",
"1lko_A",
"1mty_B",
"1mty_D",
"1mxr_A",
"1n1q_A",
"1nfv_A",
"1o9r_A",
"1oqu_A",
"1otk_A",
"1qgh_A",
"1r03_A",
"1r2f_A",
"1tjo_A",
"1tk6_A",
"1uvh_A",
"1uzr_A",
"1vlg_A",
"1w68_A",
"1yuz_A",
"1z6o_A",
"1z6o_M",
"1za0_A",
"2chp_A",
"2fjc_A",
"2fkz_A",
"2fzf_A",
"2inc_B",
"2inp_C",
"2jd7_A",
"2uw1_A",
"2uw1_B",
"2uw2_A",
"2ux1_A",
"2vzb_A",
"2za7_A",
"3dhg_A",
"3e1q_A",
"3e6s_A",
"3ee4_A",
"3qhb_A")

reorder_alphafold_xls = match(alphafold_xls_order, table=tr$tip.label)
cat(reorder_alphafold_xls, sep="\n")


#######################################################
# Re-"root" the BIONJ according to Malik 2020, Fig. 10b
# (they have an unrooted tree, but we are using their classification)
# Output clades 
#######################################################
rerooted_M20_Fig10b_trfn = "/GitHub/str2phy/ex/ferritins_M20/tree_comparison/Malik_2020_Fig10b_FerritinData_distance_matrix_BIONJtree_M20root.newick"
rerooted_M20_Fig10b_tr = read.tree(rerooted_M20_Fig10b_trfn)
rerooted_M20_Fig10b_trtable = prt(rerooted_M20_Fig10b_tr)
tail(rerooted_M20_Fig10b_trtable)
outfn = gsub(".newick", "_trtable.txt", rerooted_M20_Fig10b_trfn)
write.table(unlist_df3(rerooted_M20_Fig10b_trtable), file=outfn, sep="\t")


# Nodes having bootstraps in the distances tree:
rerooted_M20_Fig10b_trBS = rerooted_M20_Fig10b_tr
numtips = length(rerooted_M20_Fig10b_trBS$tip.label)

rerooted_M20_Fig10b_trBS$node.label = rep("", times=rerooted_M20_Fig10b_tr$Nnode)

nodenum = 58
rerooted_M20_Fig10b_trBS$node.label[nodenum-numtips] = "0"

nodenum = 59
rerooted_M20_Fig10b_trBS$node.label[nodenum-numtips] = "27"

nodenum = 60
rerooted_M20_Fig10b_trBS$node.label[nodenum-numtips] = "100"

nodenum = 73
rerooted_M20_Fig10b_trBS$node.label[nodenum-numtips] = "6"

nodenum = 77
rerooted_M20_Fig10b_trBS$node.label[nodenum-numtips] = "89"

nodenum = 93
rerooted_M20_Fig10b_trBS$node.label[nodenum-numtips] = "100"

nodenum = 99
rerooted_M20_Fig10b_trBS$node.label[nodenum-numtips] = "42"

nodenum = 100
rerooted_M20_Fig10b_trBS$node.label[nodenum-numtips] = "100"

nodenum = 102
rerooted_M20_Fig10b_trBS$node.label[nodenum-numtips] = "100"

nodenum = 103
rerooted_M20_Fig10b_trBS$node.label[nodenum-numtips] = "100"

rerooted_M20_Fig10b_trBS$node.label

rerooted_M20_Fig10b_wBS_trfn = gsub(pattern="\\.newick", replacement="_wBS.newick", x=rerooted_M20_Fig10b_trfn)
write.tree(rerooted_M20_Fig10b_trBS, file=rerooted_M20_Fig10b_wBS_trfn)


#######################################################
# Add the clade labels to another copy of the rerooted tree
#######################################################

node_in_rerooted_M20_Fig10b_tr = as.numeric(c("54",
"55",
"56",
"57",
"58",
"59",
"60",
"61",
"68",
"73",
"77",
"87",
"89",
"90",
"91",
"92",
"93",
"95",
"97",
"99",
"100",
"102",
"103"))

tipnames = c("1afr_A,1bcf_A,1bg7_A,1dps_A,1eum_A,1jgc_A,1ji4_A,1ji5_A,1jig_A,1jk0_A,1jts_A,1krq_A,1lb3_A,1lko_A,1mty_B,1mty_D,1mxr_A,1n1q_A,1nfv_A,1o9r_A,1oqu_A,1otk_A,1qgh_A,1r03_A,1r2f_A,1tjo_A,1tk6_A,1uvh_A,1uzr_A,1vlg_A,1w68_A,1yuz_A,1z6o_A,1z6o_M,1za0_A,2chp_A,2fjc_A,2fkz_A,2fzf_A,2inc_B,2inp_C,2jd7_A,2uw1_A,2uw1_B,2uw2_A,2ux1_A,2vzb_A,2za7_A,3dhg_A,3e1q_A,3e6s_A,3ee4_A,3qhb_A",
"1bcf_A,1bg7_A,1dps_A,1eum_A,1jgc_A,1ji4_A,1ji5_A,1jig_A,1jts_A,1krq_A,1lb3_A,1lko_A,1n1q_A,1nfv_A,1o9r_A,1qgh_A,1r03_A,1tjo_A,1tk6_A,1uvh_A,1vlg_A,1yuz_A,1z6o_A,1z6o_M,2chp_A,2fjc_A,2fkz_A,2fzf_A,2jd7_A,2ux1_A,2vzb_A,2za7_A,3e1q_A,3e6s_A,3qhb_A",
"1bcf_A,1bg7_A,1dps_A,1eum_A,1jgc_A,1ji4_A,1ji5_A,1jig_A,1jts_A,1krq_A,1lb3_A,1n1q_A,1nfv_A,1o9r_A,1qgh_A,1r03_A,1tjo_A,1tk6_A,1uvh_A,1vlg_A,1z6o_A,1z6o_M,2chp_A,2fjc_A,2fkz_A,2fzf_A,2jd7_A,2ux1_A,2vzb_A,2za7_A,3e1q_A,3e6s_A",
"1bcf_A,1bg7_A,1dps_A,1eum_A,1jgc_A,1ji4_A,1ji5_A,1jig_A,1jts_A,1krq_A,1lb3_A,1n1q_A,1nfv_A,1o9r_A,1qgh_A,1r03_A,1tjo_A,1tk6_A,1uvh_A,1vlg_A,1z6o_A,1z6o_M,2chp_A,2fjc_A,2fkz_A,2jd7_A,2ux1_A,2vzb_A,2za7_A,3e1q_A,3e6s_A",
"1bcf_A,1dps_A,1jgc_A,1ji4_A,1ji5_A,1jig_A,1jts_A,1n1q_A,1nfv_A,1o9r_A,1qgh_A,1tjo_A,1tk6_A,1uvh_A,2chp_A,2fjc_A,2fkz_A,2ux1_A,2vzb_A,3e1q_A",
"1dps_A,1ji4_A,1ji5_A,1jig_A,1jts_A,1n1q_A,1o9r_A,1qgh_A,1tjo_A,1tk6_A,1uvh_A,2chp_A,2fjc_A,2ux1_A,2vzb_A",
"1dps_A,1ji4_A,1ji5_A,1jig_A,1jts_A,1n1q_A,1o9r_A,1qgh_A,1tjo_A,1tk6_A,1uvh_A,2chp_A,2fjc_A,2ux1_A",
"1ji4_A,1ji5_A,1jig_A,1n1q_A,1qgh_A,2chp_A,2fjc_A,2ux1_A",
"1dps_A,1jts_A,1o9r_A,1tjo_A,1tk6_A,1uvh_A",
"1bcf_A,1jgc_A,1nfv_A,2fkz_A,3e1q_A",
"1bg7_A,1eum_A,1krq_A,1lb3_A,1r03_A,1vlg_A,1z6o_A,1z6o_M,2jd7_A,2za7_A,3e6s_A",
"1lko_A,1yuz_A,3qhb_A",
"1afr_A,1jk0_A,1mty_B,1mty_D,1mxr_A,1oqu_A,1otk_A,1r2f_A,1uzr_A,1w68_A,1za0_A,2inc_B,2inp_C,2uw1_A,2uw1_B,2uw2_A,3dhg_A,3ee4_A",
"1afr_A,1jk0_A,1mty_B,1mty_D,1mxr_A,1oqu_A,1r2f_A,1uzr_A,1w68_A,1za0_A,2inc_B,2inp_C,2uw1_A,2uw1_B,2uw2_A,3dhg_A,3ee4_A",
"1jk0_A,1mty_B,1mty_D,1mxr_A,1oqu_A,1r2f_A,1uzr_A,1w68_A,2inc_B,2inp_C,2uw2_A,3dhg_A,3ee4_A",
"1jk0_A,1mxr_A,1oqu_A,1r2f_A,1uzr_A,1w68_A,2uw2_A,3ee4_A",
"1jk0_A,1mxr_A,1oqu_A,1r2f_A,1uzr_A,1w68_A,2uw2_A",
"1jk0_A,1w68_A,2uw2_A",
"1oqu_A,1r2f_A,1uzr_A",
"1mty_B,1mty_D,2inc_B,2inp_C,3dhg_A",
"1mty_B,2inc_B,2inp_C",
"1mty_D,3dhg_A",
"1afr_A,1za0_A,2uw1_A,2uw1_B")

Malik_2020_Fig10b_clade_labels = c("all",
"type3_plus_rubreythrin",
"Dps_and_related_plus_Ferritins_and_Bacterioferritins_plus_2fzf_A",
"type3_Dps_and_related_plus_Ferritins_and_Bacterioferritins",
"Dps_and_related_plus_Bacterioferritins",
"Dps_and_related_plus_2vzb_A",
"Dps_and_related",
"Dps_and_related_clade2",
"Dps_and_related_clade1",
"Bacterioferritins",
"Ferritins",
"Rubrerythrin_Pfam02915_3taxa",
"type1_type2",
"type1_type2_minus_1otkA",
"type_1",
"RNR_R2",
"RNR_R2_minus_3ee4_A",
"RNR_R2_3taxaA",
"RNR_R2_3taxaB",
"Pfam02332_Phenol_hydrox",
"BMM_beta",
"BMM_alpha",
"type_2")


rerooted_M20_Fig10b_trLabels = rerooted_M20_Fig10b_tr
numtips = length(rerooted_M20_Fig10b_trLabels$tip.label)

rerooted_M20_Fig10b_trLabels$node.label = rep("", times=rerooted_M20_Fig10b_trLabels$Nnode)
rerooted_M20_Fig10b_trLabels$node.label[node_in_rerooted_M20_Fig10b_tr-numtips] = Malik_2020_Fig10b_clade_labels

# WORKS
# rerooted_M20_Fig10b_trLabels$node.label[node_in_rerooted_M20_Fig10b_tr-numtips] = tipnames


rerooted_M20_Fig10b_wLabels_trfn = gsub(pattern="\\.newick", replacement="_wLabels.newick", x=rerooted_M20_Fig10b_trfn)
write.tree(rerooted_M20_Fig10b_trLabels, file=rerooted_M20_Fig10b_wLabels_trfn)


pdffn = "Malik_etal_2020_Fig10b_structural_dists_BIONJ_wLabels_v1.pdf"
pdf(file=pdffn, width=12, height=12)

# Distance tree
rerooted_M20_Fig10b_trLabels_ladderized = phytools::midpoint.root(rerooted_M20_Fig10b_trLabels)
rerooted_M20_Fig10b_trLabels_ladderized = ladderize(rerooted_M20_Fig10b_trLabels_ladderized)

rerooted_M20_Fig10b_trLabels_ladderized_trtable = prt(rerooted_M20_Fig10b_trLabels_ladderized)
nodenums = (length(rerooted_M20_Fig10b_trLabels_ladderized$tip.label)+1):(length(rerooted_M20_Fig10b_trLabels_ladderized$tip.label)+rerooted_M20_Fig10b_trLabels_ladderized$Nnode)
nodetxt = rerooted_M20_Fig10b_trLabels_ladderized$node.label
nodenums = nodenums[nodetxt != ""]
edgenums = rerooted_M20_Fig10b_trLabels_ladderized_trtable$parent_br[nodenums]
nodetxt = nodetxt[nodetxt != ""]

TF = !is.na(edgenums)
edgenums = edgenums[TF]
nodetxt = nodetxt[TF]

plot.phylo(rerooted_M20_Fig10b_trLabels_ladderized, type="unrooted", lab4ut="axial", cex=0.65)
edgelabels(text=nodetxt, edge=edgenums, cex=0.5, bg="white")

add.scale.bar()
title("Malik et al. 2020 structural distances tree with Splitstree:BIONJ,\n with major clades named", cex.main=0.8)

dev.off()
cmdstr = paste0("open ", pdffn)
system(cmdstr)


#######################################################
# Calculate bipartition matches to distances tree
#######################################################

fn = "/GitHub/str2phy/ex/ferritins_M20/tree_comparison/output_filenames_v1.txt"
lines = readLines(fn, skipNul=TRUE)

TF = startsWith(x=lines, prefix="#")
lines = lines[TF==FALSE]
TF = lines == ""
lines = lines[TF==FALSE]
length(lines)

trfns = c(rerooted_M20_Fig10b_wLabels_trfn, 
rerooted_M20_Fig10b_wBS_trfn,
lines)


analysis_names = c("M20_distances_tr_wLabels",
"M20_distances_tr_wBootstraps",
"iqtree_FAMSA_AAonly_trim35",
"iqtree_FAMSA_AAonly_trim35_allmodels",
"iqtree_FAMSA_AAonly_full",
"iqtree_FAMSA_AAonly_full_allmodels",
"iqtree_FAMSA3di_3diFull",
"iqtree_FAMSA3di_3diFull_allmodels",
"iqtree_FAMSA3di_3diTrim35",
"iqtree_FAMSA3di_3diTrim35_allmodels",
"iqtree_FAMSA3di_AAfull",
"iqtree_FAMSA3di_AAfull_allmodels",
"iqtree_FAMSA3di_AA_trim35",
"iqtree_FAMSA3di_AA_trim35_allmodels",
"iqtree_FAMSA3di_BothFull",
"iqtree_FAMSA3di_BothFull_allmodels",
"iqtree_FAMSA3di_BothFullNonPart",
"iqtree_FAMSA3di_BothFullNonPart_allmodels",
"iqtree_FAMSA3di_BothTrim35",
"iqtree_FAMSA3di_BothTrim35_allmodels",
"iqtree_FAMSA3di_BothTrim35NonPart",
"iqtree_FAMSA3di_BothTrim35NonPart_allmodels",
"iqtree_USaln_3di_Full",
"iqtree_USaln_3di_Trim",
"iqtree_USaln_AAs_Full",
"iqtree_USaln_AAs_Trim",
"iqtree_USaln_Both_Full",
"iqtree_USaln_Both_Full_allmodels",
"iqtree_USaln_Both_Trim",
"iqtree_USaln_Both_Trim_allmodels",
"iqtrAF_FAMSA3di_3diFull",
"iqtrAF_FAMSA3di_3diFull_allmodels",
"iqtrAF_FAMSA3di_3diTrim35",
"iqtrAF_FAMSA3di_3diTrim35_allmodels",
"iqtrAF_FAMSA3di_AAfull",
"iqtrAF_FAMSA3di_AAfull_allmodels",
"iqtrAF_FAMSA3di_AA_trim35",
"iqtrAF_FAMSA3di_AA_trim35_allmodels",
"iqtrAF_FAMSA3di_BothFull",
"iqtrAF_FAMSA3di_BothFull_allmodels",
"iqtrAF_FAMSA3di_BothFullNonPart",
"iqtrAF_FAMSA3di_BothFullNonPart_allmodels",
"iqtrAF_FAMSA3di_BothTrim35",
"iqtrAF_FAMSA3di_BothTrim35_allmodels",
"iqtrAF_FAMSA3di_BothTrim35NonPart",
"iqtrAF_FAMSA3di_BothTrim35NonPart_allmodels",
"iqtrAF_USaln_3diFull",
"iqtrAF_USaln_3diFull_allmodels",
"iqtrAF_USaln_3diTrim35",
"iqtrAF_USaln_3diTrim35_allmodels",
"iqtrAF_USaln_AAfull",
"iqtrAF_USaln_AAfull_allmodels",
"iqtrAF_USaln_AA_trim35",
"iqtrAF_USaln_AA_trim35_allmodels",
"iqtrAF_USaln_BothFull",
"iqtrAF_USaln_BothFull_allmodels",
"iqtrAF_USaln_BothFullNonPart",
"iqtrAF_USaln_BothFullNonPart_allmodels",
"iqtrAF_USaln_BothTrim35",
"iqtrAF_USaln_BothTrim35_allmodels",
"iqtrAF_USaln_BothTrim35NonPart",
"iqtrAF_USaln_BothTrim35NonPart_allmodels")

partitioned_modelnames = c("iqtree_FAMSA3di_BothFull",
"iqtree_FAMSA3di_BothFull_allmodels",
"iqtree_FAMSA3di_BothTrim35",
"iqtree_FAMSA3di_BothTrim35_allmodels",
"iqtree_USaln_Both_Full",
"iqtree_USaln_Both_Full_allmodels",
"iqtree_USaln_Both_Trim",
"iqtree_USaln_Both_Trim_allmodels",
"iqtrAF_FAMSA3di_BothFull",
"iqtrAF_FAMSA3di_BothFull_allmodels",
"iqtrAF_FAMSA3di_BothTrim35",
"iqtrAF_FAMSA3di_BothTrim35_allmodels",
"iqtrAF_USaln_BothFull",
"iqtrAF_USaln_BothFull_allmodels",
"iqtrAF_USaln_BothTrim35",
"iqtrAF_USaln_BothTrim35_allmodels")


source("/GitHub/str2phy/Rsrc/str2phy_v1.R")
source("/GitHub/str2phy/Rsrc/parsing_iqtree_file_v1.R")

trs_list = list()
dtf_list = list()
sum_matches = rep(0, times=length(trfns))
sum_deepnodes = rep(0, times=length(trfns))
sum_M20boots = rep(0, times=length(trfns))
trlen = rep(0, times=length(trfns))
sumBoots = rep(0, times=length(trfns))
tdists_mat = NULL
tdists_mat_s1 = NULL

spr = NULL
RF = NULL
normRF = NULL
normSim_RF = NULL
wRF = NULL
norm_wRF = NULL
normSim_wRF = NULL
KF = NULL
pathDist = NULL

# Normalized, with total treelength summing to 1 (s1)
spr_s1 = NULL
RF_s1 = NULL
normRF_s1 = NULL
normSim_RF_s1 = NULL
wRF_s1 = NULL
norm_wRF_s1 = NULL
normSim_wRF_s1 = NULL
KF_s1 = NULL
pathDist_s1 = NULL


# Get the deep nodes reflecting major categories in the Malik et al. 2020 tree
trtable_rerooted_M20_Fig10b_trLabels = prt(rerooted_M20_Fig10b_trLabels)
deepnode_TF = trtable_rerooted_M20_Fig10b_trLabels$label != ""
deepnode_TF[1:length(rerooted_M20_Fig10b_trLabels$tip.label)] = FALSE
deepnode_tipnames = trtable_rerooted_M20_Fig10b_trLabels$tipnames[deepnode_TF]
deepnode_tipnames

# Get the nodes Malik et al. 2020 bootstrapped
trtable_rerooted_M20_Fig10b_trBS = prt(rerooted_M20_Fig10b_trBS)
MalikBS_TF = trtable_rerooted_M20_Fig10b_trBS$label != ""
MalikBS_TF[1:length(rerooted_M20_Fig10b_trBS$tip.label)] = FALSE
MalikBS_tipnames = trtable_rerooted_M20_Fig10b_trBS$tipnames[MalikBS_TF]
MalikBS_tipnames

iqtree_fns = gsub(pattern="\\.contree", replacement=".iqtree", x=trfns)
iqtree_fns

iqtree_stats = NULL

iqtree_stats_names = c("n",
"best_lnL",
"best_lnL_k",
"best_lnL_k_wo_brlens",
"best_lnL_Model",
"best_AIC",
"best_AIC_lnL",
"best_AIC_k",
"best_AIC_k_wo_brlens",
"best_AIC_Model",
"best_AICc",
"best_AICc_lnL",
"best_AICc_k",
"best_AICc_k_wo_brlens",
"best_AICc_Model",
"best_BIC",
"best_BIC_lnL",
"best_BIC_k",
"best_BIC_k_wo_brlens",
"best_BIC_Model")

txt = paste0("\nCalculating statistics for ", length(trfns), " tree files. Calc #: ")
cat(txt)
for (i in 1:length(trfns))
	{
	cat(i, ",", sep="")
	
	trs_list[[i]] = read.tree(trfns[i])
	#print(length(trs_list[[i]]$tip.label))
	trs_list[[i]]$tip.label = gsub(pattern="\\.pdb", replacement="", x=trs_list[[i]]$tip.label)

	tr1 = distance_tr
	tr1 = phytools::midpoint.root(tr1)
	tr1 = ladderize(tr1)
	tr1 = read.tree(file="", text=write.tree(tr1, file=""))
	tr1$tip.label = gsub(pattern="\\.pdb", replacement="", x=tr1$tip.label)

	tr2 = trs_list[[i]]
	tr2 = phytools::midpoint.root(tr2)
	tr2 = ladderize(tr2)
	tr2 = read.tree(file="", text=write.tree(tr2, file=""))
	tr2$tip.label = gsub(pattern="\\.pdb", replacement="", x=tr2$tip.label)
	
	trlen[i] = sum(tr2$edge.length)
	sumBoots[i] = sum(as.numeric(tr2$node.label), na.rm=TRUE)
	dtf = check_bipartitions_in_tree2(tr1=tr1, tr2=tr2, remove_txt="\\.pdb")
	dtf
	
	sum_matches[i] = sum(as.numeric(dtf$bipart))
	
	# Sum matches for subsets of the nodes
	tr2_table = prt(tr2)
	# Deep nodes representing established groups
	TF = tr2_table$tipnames %in% deepnode_tipnames
	sum_deepnodes[i] = sum(TF)
	# Get the nodes Malik et al. 2020 bootstrapped
	TF = tr2_table$tipnames %in% MalikBS_tipnames
	sum_M20boots[i] = sum(TF)

	
	# Tree-to-tree distances
	tdists = phangorn::treedist(tree1=tr1, tree2=tr2, check.labels=TRUE)
	tdists_mat = rbind(tdists_mat, tdists)

	spr = rbind(spr, phangorn::sprdist(tree1=tr1, tree2=tr2))
	RF = c(RF, phangorn::RF.dist(tree1=tr1, tree2=tr2))
	normRF = c(normRF, phangorn::RF.dist(tree1=tr1, tree2=tr2, normalize=TRUE))
	normSim_RF = c(normSim_RF, 1-phangorn::RF.dist(tree1=tr1, tree2=tr2, normalize=TRUE))
	wRF = c(wRF, phangorn::wRF.dist(tree1=tr1, tree2=tr2))
	norm_wRF = c(norm_wRF, phangorn::wRF.dist(tree1=tr1, tree2=tr2, normalize=TRUE))
	normSim_wRF = c(normSim_wRF, 1-phangorn::wRF.dist(tree1=tr1, tree2=tr2, normalize=TRUE))
	KF = c(KF, phangorn::KF.dist(tree1=tr1, tree2=tr2))
	pathDist = c(pathDist, phangorn::path.dist(tree1=tr1, tree2=tr2))
	
	# Tree-to-tree distances, when the trees are normalized to sum to 1 (=s1)
	tr1_s1 = tr1
	tr1_s1$edge.length = tr1$edge.length / sum(tr1$edge.length)

	tr2_s1 = tr2
	tr2_s1$edge.length = tr2$edge.length / sum(tr2$edge.length)

	tdists_s1 = phangorn::treedist(tree1=tr1_s1, tree2=tr2_s1, check.labels=TRUE)
	tdists_mat_s1 = rbind(tdists_mat_s1, tdists_s1)

	spr_s1 = rbind(spr_s1, phangorn::sprdist(tree1=tr1_s1, tree2=tr2_s1))
	RF_s1 = c(RF_s1, phangorn::RF.dist(tree1=tr1_s1, tree2=tr2_s1))
	normRF_s1 = c(normRF_s1, phangorn::RF.dist(tree1=tr1_s1, tree2=tr2_s1, normalize=TRUE))
	normSim_RF_s1 = c(normSim_RF_s1, 1-phangorn::RF.dist(tree1=tr1_s1, tree2=tr2_s1, normalize=TRUE))
	wRF_s1 = c(wRF_s1, phangorn::wRF.dist(tree1=tr1_s1, tree2=tr2_s1))
	norm_wRF_s1 = c(norm_wRF_s1, phangorn::wRF.dist(tree1=tr1_s1, tree2=tr2_s1, normalize=TRUE))
	normSim_wRF_s1 = c(normSim_wRF_s1, 1-phangorn::wRF.dist(tree1=tr1_s1, tree2=tr2_s1, normalize=TRUE))
	KF_s1 = c(KF_s1, phangorn::KF.dist(tree1=tr1_s1, tree2=tr2_s1))
	pathDist_s1 = c(pathDist_s1, phangorn::path.dist(tree1=tr1_s1, tree2=tr2_s1))
	
	# Add _s1 to names
	colnames(spr_s1) = paste0(colnames(spr), "_s1")
	
	# Add in statistics from the .iqtree files
	iqtree_fn = iqtree_fns[i]
	
	if (endsWith(x=iqtree_fn, suffix=".iqtree") == TRUE)
		{
		# If *not* partitioned:
		partitioned_TF = analysis_names[i] %in% partitioned_modelnames
		partitioned_TF
	
		if (partitioned_TF == FALSE)
			{
			iqtree_df = read_iqtree_model_scores(iqtree_fn, start_txt="List of models sorted by BIC scores: ", calc_extra=TRUE)
	
			n = iqtree_df$ndata_AIC[1]
	
			# Best lnL model
			best_lnL = max(iqtree_df$LogL)
			best_lnL_k = iqtree_df$k[iqtree_df$LogL == best_lnL][1]
			best_lnL_k_wo_brlens = iqtree_df$k_wo_brlens[iqtree_df$LogL == best_lnL][1]
			best_lnL_Model = iqtree_df$Model[iqtree_df$LogL == best_lnL][1]
	
			# Best AIC model
			best_AIC = min(iqtree_df$AIC)
			best_AIC_lnL = iqtree_df$LogL[iqtree_df$AIC == best_AIC][1]
			best_AIC_k = iqtree_df$k[iqtree_df$AIC == best_AIC][1]
			best_AIC_k_wo_brlens = iqtree_df$k_wo_brlens[iqtree_df$AIC == best_AIC][1]
			best_AIC_Model = iqtree_df$Model[iqtree_df$AIC == best_AIC][1]

			# Best AICc model
			best_AICc = min(iqtree_df$AICc)
			best_AICc_lnL = iqtree_df$LogL[iqtree_df$AICc == best_AICc][1]
			best_AICc_k = iqtree_df$k[iqtree_df$AICc == best_AICc][1]
			best_AICc_k_wo_brlens = iqtree_df$k_wo_brlens[iqtree_df$AICc == best_AICc][1]
			best_AICc_Model = iqtree_df$Model[iqtree_df$AICc == best_AICc][1]

			# Best BIC model
			best_BIC = min(iqtree_df$BIC)
			best_BIC_lnL = iqtree_df$LogL[iqtree_df$BIC == best_BIC][1]
			best_BIC_k = iqtree_df$k[iqtree_df$BIC == best_BIC][1]
			best_BIC_k_wo_brlens = iqtree_df$k_wo_brlens[iqtree_df$BIC == best_BIC][1]
			best_BIC_Model = iqtree_df$Model[iqtree_df$BIC == best_BIC][1]
	
			iqtree_nums = c(n,
			best_lnL,
			best_lnL_k,
			best_lnL_k_wo_brlens,
			best_lnL_Model,
			best_AIC,
			best_AIC_lnL,
			best_AIC_k,
			best_AIC_k_wo_brlens,
			best_AIC_Model,
			best_AICc,
			best_AICc_lnL,
			best_AICc_k,
			best_AICc_k_wo_brlens,
			best_AICc_Model,
			best_BIC,
			best_BIC_lnL,
			best_BIC_k,
			best_BIC_k_wo_brlens,
			best_BIC_Model)
			
			names(iqtree_nums) = iqtree_stats_names
			} # END if (partitioned_TF == FALSE)

		if (partitioned_TF == TRUE)
			{
			iqtree_df = read_iqtree_model_scores(iqtree_fn, start_txt="List of best-fit models per partition:", calc_extra=TRUE, partitioned_TF=TRUE)
	
			n = iqtree_df$ndata_AIC[1]
	
			# Best lnL model
			best_lnL = max(iqtree_df$LogL)
			best_lnL_k = iqtree_df$k[iqtree_df$LogL == best_lnL][1]
			best_lnL_k_wo_brlens = iqtree_df$k_wo_brlens[iqtree_df$LogL == best_lnL][1]
			best_lnL_Model = iqtree_df$Model[iqtree_df$LogL == best_lnL][1]
	
			# Best AIC model
			best_AIC = min(iqtree_df$AIC)
			best_AIC_lnL = iqtree_df$LogL[iqtree_df$AIC == best_AIC][1]
			best_AIC_k = iqtree_df$k[iqtree_df$AIC == best_AIC][1]
			best_AIC_k_wo_brlens = iqtree_df$k_wo_brlens[iqtree_df$AIC == best_AIC][1]
			best_AIC_Model = iqtree_df$Model[iqtree_df$AIC == best_AIC][1]

			# Best AICc model
			best_AICc = min(iqtree_df$AICc)
			best_AICc_lnL = iqtree_df$LogL[iqtree_df$AICc == best_AICc][1]
			best_AICc_k = iqtree_df$k[iqtree_df$AICc == best_AICc][1]
			best_AICc_k_wo_brlens = iqtree_df$k_wo_brlens[iqtree_df$AICc == best_AICc][1]
			best_AICc_Model = iqtree_df$Model[iqtree_df$AICc == best_AICc][1]

			# Best BIC model
			best_BIC = min(iqtree_df$BIC)
			best_BIC_lnL = iqtree_df$LogL[iqtree_df$BIC == best_BIC][1]
			best_BIC_k = iqtree_df$k[iqtree_df$BIC == best_BIC][1]
			best_BIC_k_wo_brlens = iqtree_df$k_wo_brlens[iqtree_df$BIC == best_BIC][1]
			best_BIC_Model = iqtree_df$Model[iqtree_df$BIC == best_BIC][1]
	
			iqtree_nums = c(n,
			best_lnL,
			best_lnL_k,
			best_lnL_k_wo_brlens,
			best_lnL_Model,
			best_AIC,
			best_AIC_lnL,
			best_AIC_k,
			best_AIC_k_wo_brlens,
			best_AIC_Model,
			best_AICc,
			best_AICc_lnL,
			best_AICc_k,
			best_AICc_k_wo_brlens,
			best_AICc_Model,
			best_BIC,
			best_BIC_lnL,
			best_BIC_k,
			best_BIC_k_wo_brlens,
			best_BIC_Model)
			
			names(iqtree_nums) = iqtree_stats_names
			} # END if (partitioned_TF == TRUE)
		} else {
		# if (endsWith(x=iqtree_fn, suffix=".iqtree") == FALSE)
		iqtree_nums = rep(NA, times=length(iqtree_stats_names))
		names(iqtree_nums) = iqtree_stats_names
		} # END if (endsWith(x=iqtree_fn, suffix="\\.iqtree") == TRUE)

	iqtree_stats = rbind(iqtree_stats, iqtree_nums)
	dtf_list[[i]] = dtf
	} # END for (i in 1:length(trfns))
cat("...done!\n")
class(trs_list) = c("list", "multiPhylo")

tdists_df = as.data.frame(tdists_mat, stringsAsFactors=FALSE)
names(tdists_df) = c("symDiff", "branchscoreDiff","pathDiff","quadPathDiff")
tdists_df2 = tdists_df
tdists_df2$symDiff = tdists_df2$symDiff / max(tdists_df2$symDiff)
tdists_df2$branchscoreDiff = tdists_df2$branchscoreDiff / max(tdists_df2$branchscoreDiff)
tdists_df2$pathDiff = tdists_df2$pathDiff / max(tdists_df2$pathDiff)
tdists_df2$quadPathDiff = tdists_df2$quadPathDiff / max(tdists_df2$quadPathDiff)
names(tdists_df2) = c("relSymDiff", "relBranchscore","relPathDiff","relQuadPathDiff")

tdists_df_s1 = as.data.frame(tdists_mat_s1, stringsAsFactors=FALSE)
names(tdists_df_s1) = c("symDiff_s1", "branchscoreDiff_s1","pathDiff_s1","quadPathDiff_s1")
tdists_df2_s1 = tdists_df_s1
tdists_df2_s1$symDiff_s1 = tdists_df2_s1$symDiff / max(tdists_df2_s1$symDiff)
tdists_df2_s1$branchscoreDiff_s1 = tdists_df2_s1$branchscoreDiff / max(tdists_df2_s1$branchscoreDiff)
tdists_df2_s1$pathDiff_s1 = tdists_df2_s1$pathDiff / max(tdists_df2_s1$pathDiff)
tdists_df2_s1$quadPathDiff_s1 = tdists_df2_s1$quadPathDiff / max(tdists_df2_s1$quadPathDiff)
names(tdists_df2_s1) = c("relSymDiff_s1", "relBranchscore_s1","relPathDiff_s1","relQuadPathDiff_s1")


iqtree_stats_df = as.data.frame(iqtree_stats, stringsAsFactors=FALSE)
iqtree_stats_df

alldf = cbind(tdists_df, tdists_df_s1, tdists_df2, tdists_df2_s1, spr, RF, normRF, normSim_RF, wRF, norm_wRF, normSim_wRF, KF, pathDist, spr_s1, RF_s1, normRF_s1, normSim_RF_s1, wRF_s1, norm_wRF_s1, normSim_wRF_s1, KF_s1, pathDist_s1, iqtree_stats_df)
head(alldf)

sum_matches


# Distances tree against itself:
dtf_list[[1]]


# Compare trees
sq_status = Quartet::QuartetStatus(trs_list)
SimilarityMetrics(sq_status)

res = cbind(analysis_names, sum_matches, sum_deepnodes, sum_M20boots, trlen, sumBoots, SimilarityMetrics(sq_status))
resdf = as.data.frame(res, stringsAsFactors=FALSE)

#RFdist = Quartet::RobinsonFoulds(sq_status, similarity=TRUE) # deprecated
RFdist = Quartet::RawSymmetricDifference(sq_status, similarity=TRUE)
relRFdist = RFdist / max(RFdist)

resdf = cbind(resdf, RFdist, relRFdist)

resdf = dfnums_to_numeric(resdf)

resalldf = cbind(resdf, alldf)
resalldf

bigtable_fn = "60iqtrees_compared_df_v3.txt"
write.table(resalldf, file=bigtable_fn, quote=FALSE, sep="\t")
head(resalldf)
tail(resalldf)

cmdstr = paste0("open ", bigtable_fn)
system(cmdstr)


resdf[rev(order(resalldf$MarczewskiSteinhaus)),]
resdf[order(resalldf$MarczewskiSteinhaus),]

resdf[rev(order(resalldf$relRFdist)),]
resdf[order(resalldf$relRFdist),]


#######################################################
# Plot distances against each other
#######################################################

# Old; many are redundant

# Original order
cols = c("sum_matches", "sum_deepnodes", "sum_M20boots", "trlen", "sumBoots", "DoNotConflict", "MarczewskiSteinhaus", "QuartetDivergence", "SimilarityToReference", "RFdist", "relRFdist", "symDiff", "branchscoreDiff", "pathDiff", "relQuadPathDiff", "relSymDiff_s1", "relBranchscore_s1", "relPathDiff_s1", "relQuadPathDiff_s1", "spr", "spr_extra", "rf", "hdist", "RF", "normRF", "normSim_RF", "wRF", "norm_wRF", "normSim_wRF", "KF", "pathDist", "spr_s1", "spr_extra_s1", "rf_s1", "hdist_s1", "RF_s1", "normRF_s1", "normSim_RF_s1", "wRF_s1", "norm_wRF_s1", "normSim_wRF_s1", "KF_s1")

# Reordered
cols = c("sum_matches", "sum_deepnodes", "sum_M20boots", "trlen", "sumBoots", "spr", "spr_extra", "spr_s1", "spr_extra_s1", "DoNotConflict", "MarczewskiSteinhaus", "QuartetDivergence", "SimilarityToReference", "RFdist", "relRFdist", "symDiff", "RF", "normRF", "normSim_RF", "rf", "hdist", "rf_s1", "hdist_s1", "RF_s1", "normRF_s1", "normSim_RF_s1", "relSymDiff_s1", "branchscoreDiff", "relQuadPathDiff", "wRF", "norm_wRF", "normSim_wRF", "KF", "relBranchscore_s1", "wRF_s1", "norm_wRF_s1", "normSim_wRF_s1", "KF_s1", "pathDist", "pathDiff", "relPathDiff_s1", "relQuadPathDiff_s1")

# symmetricDiff = rf = many many similar metrics when you have a bifurcating tree
# branchScoreDiff = weighted RFdist = wRFdist =  KF
# branchScoreDiff_s1 = wRFdist_s1 = KF_s1

#cols = c("sum_matches", "sum_deepnodes", "sum_M20boots", "trlen", "sumBoots", "RFdist", "symDiff", "branchscoreDiff", "pathDiff", "relQuadPathDiff", "relBranchscore_s1", "relQuadPathDiff_s1", "spr", "spr_extra", "hdist", "norm_wRF",  "wRF_s1")

TF = cols %in% names(resalldf)
all(TF)

tmpdf = resalldf[, cols]

head(tmpdf)
dim(tmpdf)

wd = "/GitHub/str2phy/ex/ferritins_M20/tree_comparison"
setwd(wd)

pdffn = "pairs_42x42.pdf"
pdf(file=pdffn, width=20, height=20)

pairs(tmpdf, pch=".")

dev.off()
cmdstr = paste0("open ", pdffn)
system(cmdstr)


# Original order
# DoNotConflict = many others including "RFdist" & many many similar metrics when you have a bifurcating tree
# symmetricDiff = rf = RF = number of matching clades
# branchScoreDiff = weighted RFdist = wRFdist =  KF
# branchScoreDiff_s1 = wRFdist_s1 = KF_s1
# pathDist = pathDiff
cols = c("sum_matches", "sum_deepnodes", "sum_M20boots", "trlen", "sumBoots", "spr", "DoNotConflict", "symDiff", "branchscoreDiff", "relBranchscore_s1", "pathDiff", "relQuadPathDiff", "relQuadPathDiff_s1")

TF = cols %in% names(resalldf)
all(TF)

tmpdf = resalldf[, cols]

head(tmpdf)
dim(tmpdf)

# Make SPR distance negative for better plot
tmpdf$spr = -1 * tmpdf$spr

wd = "/GitHub/str2phy/ex/ferritins_M20/tree_comparison"
setwd(wd)

colors=c("black","black","red","red","red","red","yellow4","yellow4","yellow4","yellow4","orange3","orange3","orange3","orange3","blue","blue","blue","blue","lightblue","lightblue","lightblue","lightblue","yellow4","yellow4","orange3","orange3","blue","blue","lightblue","lightblue","yellow4","yellow4","yellow4","yellow4","orange3","orange3","orange3","orange3","blue","blue","blue","blue","lightblue","lightblue","lightblue","lightblue","yellow4","yellow4","yellow4","yellow4","orange3","orange3","orange3","orange3","blue","blue","blue","blue","lightblue","lightblue","lightblue","lightblue")

symbols=c("+","+","A","A","A","A","d","d","d","d","A","A","A","A","B","B","B","B","B","B","B","B","d","d","A","A","B","B","B","B","d","d","d","d","A","A","A","A","B","B","B","B","B","B","B","B","d","d","d","d","A","A","A","A","B","B","B","B","B","B","B","B")

pdffn = "pairs_13x13.pdf"
pdf(file=pdffn, width=20, height=20)

pairs(tmpdf, pch=".")

library(psych)
psych::pairs.panels(tmpdf, 
             method = "pearson", # correlation method
						 hist.col = "#00AFBB",
             density = TRUE,  # show density plotspch=".")
						 lm = TRUE,
						 pch=symbols,
						 col=colors)
						 
dev.off()
cmdstr = paste0("open ", pdffn)
system(cmdstr)


#######################################################
# Plot of distance tree branchlengths, vs. bootstraps
#######################################################
brlens = as.numeric(dtf_list[[1]]$brlen2)
notNA = !is.na(brlens)
xlims = as.numeric(c(0.0, max(brlens[notNA])))


pdffn = "distance_brlens_v_60iqtrees_bootstraps.pdf"
pdf(file=pdffn, width=6, height=6)

plot(x=c(0.0, max(brlens)), y=c(0,100), col="white", pch=".", xlim=xlims, ylim=c(0,100), xlab="branch length (on distance tree)", ylab="bootstrap (on sequence tree)")
title("Ferritin superfamily: Branch lengths (structural distance tree)\nvs. bootstraps (sequence phylogenies #1-16)", cex=0.6)
for (i in 3:length(dtf_list))
	{
	dtf = dtf_list[[i]]
	TF = dtf$bipart[notNA] == 1
	points(x=jitter(brlens[notNA][TF]), jitter(as.numeric(dtf$bootstrap2[notNA][TF])), pch=as.character(i-2), cex=0.5)
	}

dev.off()
cmdstr = paste0("open ", pdffn)
system(cmdstr)


pdffn = "distance_brlens_v_60iqtrees_brlens.pdf"
pdf(file=pdffn, width=6, height=6)

plot(x=c(0.0, max(brlens)), y=c(0,100), col="white", pch=".", xlim=xlims, ylim=c(0,1.6), xlab="branch length (on distance tree)", ylab="bootstrap (on sequence tree)")
title("Ferritin superfamily: Branch lengths (structural distance tree)\nvs. branchlengths (sequence phylogenies #1-16)", cex=0.6)
for (i in 3:length(dtf_list))
	{
	dtf = dtf_list[[i]]
	TF = dtf$bipart[notNA] == 1
	points(x=jitter(brlens[notNA][TF]), jitter(as.numeric(dtf$brlen2[notNA][TF])), pch=as.character(i-2), cex=0.5)
	}

dev.off()
cmdstr = paste0("open ", pdffn)
system(cmdstr)


pdffn = "distance_brlens_v_brlens.pdf"
pdf(file=pdffn, width=6, height=6)

for (i in 2:length(dtf_list))
	{
	dtf = dtf_list[[i]]
	
	brlens = as.numeric(dtf_list[[2]]$brlen2)
	notNA = !is.na(brlens)
	TF = dtf$bipart[notNA] == 1
	num_matches = sum(TF)
	
	xlims = as.numeric(c(0.0, max(brlens[notNA])))
	ylims = c(0, 1.6)
	x = brlens[notNA][TF]
	y = dtf$brlen2[notNA][TF]

	linear_regression_plot(x, y, xlabel="branch length (on distance tree)", ylabel="branch length (on sequence tree)", tmppch=1, xlim=xlims, ylim=ylims)
	
	txt = paste0("Ferritin superfamily: Branch lengths (structural distance tree)\nvs. branchlengths (", num_matches, "/52 matches for #", i, ": ", analysis_names[i], ")")
	title(txt, cex.main=0.8)
	}

dev.off()
cmdstr = paste0("open ", pdffn)
system(cmdstr)


pdffn = "distance_brlens_v_bootstraps.pdf"
pdf(file=pdffn, width=6, height=6)

for (i in 2:length(dtf_list))
	{
	dtf = dtf_list[[i]]

	brlens = as.numeric(dtf_list[[2]]$brlen2)
	notNA = !is.na(brlens)
	TF = dtf$bipart[notNA] == 1
	num_matches = sum(TF)
	
	xlims = as.numeric(c(0.0, max(brlens[notNA])))
	ylims = c(0, 100)
	x = brlens[notNA][TF]
	y = dtf$bootstrap2[notNA][TF]
	
	x = as.numeric(x)
	y = as.numeric(y)
	y
	# Filter out missing bootstraps
	x = x[!is.na(y)]
	y = y[!is.na(y)]
	
	
	linear_regression_plot(x, y, xlabel="branch length (on distance tree)", ylabel="bootstrap (on sequence tree)", tmppch=1, xlim=xlims, ylim=ylims, legend_x=0.15, legend_y=25)
	
	txt = paste0("Ferritin superfamily: Branch lengths (structural distance tree)\nvs. bootstraps (", num_matches, "/52 matches for #", i, ": ", analysis_names[i], ")")
	title(txt, cex.main=0.8)
	}

dev.off()
cmdstr = paste0("open ", pdffn)
system(cmdstr)


# Make an array of results
dta = array(data=NA, dim=c(nrow(dtf_list[[1]]), ncol(dtf_list[[1]]), length(dtf_list)))
for (i in 1:length(dtf_list))
	{
	dtm = as.matrix(dtf_list[[i]])
	dta[,,i] = dtm
	}

dtf_list[[3]]
dtf_list[[4]]

# bipartitions
bipartitions_df = as.data.frame(dta[,2,], stringsAsFactors=FALSE)
bipartitions_df = df_to_numeric(bipartitions_df)
names(bipartitions_df) = analysis_names
bipartitions_df

# branch lengths
brlens_df = as.data.frame(dta[,3,], stringsAsFactors=FALSE)
brlens_df = df_to_numeric(brlens_df)
names(brlens_df) = analysis_names
brlens_df

outfn = "brlens_df.txt"
write.table(brlens_df, file=outfn, quote=FALSE, sep="\t")
system(paste0("open ", outfn))

# bootstraps
bootstraps_df = as.data.frame(dta[,4,], stringsAsFactors=FALSE)
bootstraps_df = df_to_numeric(bootstraps_df)
names(bootstraps_df) = analysis_names
bootstraps_df

outfn = "bootstraps_df.txt"
write.table(bootstraps_df, file=outfn, quote=FALSE, sep="\t")
system(paste0("open ", outfn))


bootstraps_mat = matrix(data=as.numeric(unlist(bootstraps_df[,2:ncol(bootstraps_df)])), nrow=nrow(bootstraps_df), byrow=FALSE)
bootstraps_mat

# Number of matches
colSums(!is.na(bootstraps_mat), na.rm=TRUE)

# Node heights above root: absolute
node_hts_df = as.data.frame(dta[,5,], stringsAsFactors=FALSE)
node_hts_df = df_to_numeric(node_hts_df)
names(node_hts_df) = analysis_names
node_hts_df
apply(X=node_hts_df, MARGIN=2, FUN=max, na.rm=TRUE)

# Node heights above root: relative
rel_hts_df = as.data.frame(dta[,6,], stringsAsFactors=FALSE)
rel_hts_df = df_to_numeric(rel_hts_df)
names(rel_hts_df) = analysis_names
rel_hts_df
apply(X=rel_hts_df, MARGIN=2, FUN=max, na.rm=TRUE)

outfn = "rel_hts_df.txt"
write.table(rel_hts_df, file=outfn, quote=FALSE, sep="\t")
system(paste0("open ", outfn))


pdffn = "rel_hts_above_root.pdf"
pdf(file=pdffn, width=6, height=6)


for (i in 2:length(dtf_list))
	{
	dtf = dtf_list[[i]]

	brlens = as.numeric(dtf_list[[2]]$brlen2)
	notNA = !is.na(brlens)
	TF = dtf$bipart[notNA] == 1
	num_matches = sum(TF)

	x = as.numeric(dtf_list[[2]]$rel_ht[notNA][TF])
	y = as.numeric(dtf$rel_ht[notNA][TF])
	
	xlims = c(0, 1)
	ylims = c(0, 1)
	linear_regression_plot(x, y, xlabel="relative node height above root (on distance tree)", ylabel="relative node height above root (on sequence tree)", tmppch=1, xlim=xlims, ylim=ylims, cex=0.6)

	txt = paste0("Ferritin superfamily: Relative node heights above root (structural distance tree)\nvs. relative node heights above root (", num_matches, "/52 matches for #", i, ": ", analysis_names[i], ")")

	title(txt, cex.main=0.6)
	}

dev.off()
cmdstr = paste0("open ", pdffn)
system(cmdstr)


#######################################################
# Display the PDB trees
#######################################################

txt_to_cut1 = "/GitHub/str2phy/ex/ferritins_M20/iqtree/full/"
txt_to_cut2 = "/GitHub/str2phy/ex/ferritins_M20/iqtree/"

pdffn = "53_ferritins_PDB_contrees_v1.pdf"
pdf(file=pdffn, width=12, height=12)

# Distance tree
tr = phytools::midpoint.root(distance_tr)
tr = ladderize(tr)
plot.phylo(tr, type="unrooted", lab4ut="axial", cex=0.65)
add.scale.bar()
title("Malik et al. 2020 structural distances tree with Splitstree:BIONJ")


i=1
for (i in 1:length(trfns))
	{
	titletxt = gsub(pattern=txt_to_cut1, replacement="", x=trfns[i])
	titletxt = gsub(pattern=txt_to_cut2, replacement="", x=titletxt)
	
	tr = read.tree(trfns[i])
	tr = phytools::midpoint.root(tr)
	tr = ladderize(tr)
	trtable = prt(tr)
	
	nodenums = (length(tr$tip.label)+1):(length(tr$tip.label)+tr$Nnode)
	nodetxt = tr$node.label
	nodenums = nodenums[nodetxt != ""]
	edgenums = trtable$parent_br[nodenums]
	nodetxt = nodetxt[nodetxt != ""]
	
	# Plot the tree
	plot.phylo(tr, type="unrooted", lab4ut="axial", cex=0.65)
	edgelabels(text=nodetxt, edge=edgenums, cex=0.5, bg="white")
	add.scale.bar()
	title(titletxt)
	}


dev.off()
cmdstr = paste0("open ", pdffn)
system(cmdstr)


#######################################################
# Extract the model selection results
#######################################################
```
