## Supplementary material for "Tertiary-interaction characters enable fast, model-based structural phylogenetics beyond the twilight zone": 01_pdbs_to_3di_v1.R

```
#######################################################
# PDB structures to AA & 3di sequences
#######################################################
#library(bio3d)
source("/GitHub/str2phy/Rsrc/str2phy_v1.R")


wd = "/GitHub/str2phy/ex/ferritins_M20/rawdata"
setwd(wd)

pdb_fns = list.files(path=".", pattern="*.pdb")
length(pdb_fns)
# 53

fns_df = str_to_3di(str_fn=pdb_fns, suffix="\\.pdb")
fns_df


# Make a fasta file of the AAs
fasta_fns = fns_df$di3_fn
# ls -1 *_aa.fasta | while read fn ; do cat "$fn" >> 53_ferritin_AAs.fasta; done
cat *_aa.fasta > 53_ferritin_AAs.fasta
head 53_ferritin_AAs.fasta
tail 53_ferritin_AAs.fasta


# Make a fasta file of the 3dis
#ls -1 *_3di.fasta | while read fn ; do cat "$fn" >> 53_ferritin_3dis.fasta; done
cd /GitHub/str2phy/ex/ferritins_M20/rawdata
cat *_3di.fasta > 53_ferritin_3dis.fasta
cat *_aa.fasta > 53_ferritin_AAs.fasta

head 53_ferritin_3dis.fasta
tail 53_ferritin_3dis.fasta


#######################################################
# USalign run
#######################################################
# Put the chains in a "chains" directory
mkdir chains
cp *.pdb chains
rm chainslist.txt

# DO NOT call the new file "list_of_chains.txt", 
# this cat command loops & fills up hard drive!!
cat *_chains.txt > chainslist.txt  
head chainslist.txt

USalign -dir chains/ chainslist.txt -mol prot -mm 4 -o ferritins_pymol | tee USalign_ferritins_so1.txt &

open USalign_ferritins_so1.txt

# Edit in BBedit, re-save to:
usalign_ferritins_aa.fasta
# #Total CPU time is 17.15 seconds

iqtree -t BIONJ -s usalign_ferritins_aa.fasta -m LG+F+G --ufboot 1000 -bnni | tee iqtree_ferritins_aa.so1.txt &
```
