## Supplementary material for "Tertiary-interaction characters enable fast, model-based structural phylogenetics beyond the twilight zone": __cmds_ferritins_M20_famsa3di_USaln_then_trim_iqtree_v1.R

```
#######################################################
# Let's try FAMSA
# https://github.com/refresh-bio/FAMSA
#
#
# FAMSA: Fast and accurate multiple sequence alignment of ...
# Nature Journal
# S Deorowicz · 2016 
#######################################################

cd /GitHub/FAMSA
make
cp famsa /usr/local/bin

# See also /GitHub/str2phy/substmat/ for
# creation & compilation of famsa3di,
# a version of famsa with the 3di cost matrix


#######################################################
# Align the 3dis with FAMSA
#######################################################
cd /GitHub/str2phy/ex/ferritins_M20/rawdata/
cp 53_ferritin_3dis.fasta /GitHub/str2phy/ex/ferritins_M20/02_FAMSA_3dis_aln_v1
cp 53_ferritin_AAs.fasta /GitHub/str2phy/ex/ferritins_M20/02_FAMSA_3dis_aln_v1

cd /GitHub/str2phy/ex/ferritins_M20/02_FAMSA_3dis_aln_v1/
famsa3di 53_ferritin_3dis.fasta 53_ferritin_3dis.fasta.aln

# View in AliView
# Save to Fasta


# Convert the AAs to match 3di alignment

# In R:
#######################################################
# Convert an aa fasta file to a 3di fasta file
#######################################################
library(ape)
library(seqinr)
library(BioGeoBEARS)
sourceall("/GitHub/bioinfRhints/Rsrc")
source("/GitHub/str2phy/Rsrc/str2phy_v1.R")

wd = "/GitHub/str2phy/ex/ferritins_M20/02_FAMSA_3dis_aln_v1/"
setwd(wd)

aligned_fn = "53_ferritin_3dis.fasta.aln"
unaligned_fn = "53_ferritin_AAs.fasta"
outfn = paste0(unaligned_fn, ".aln")
aligned_AAs = align_3dis_to_AAs(aligned_fn, unaligned_fn, outfn)
moref(outfn)
#######################################################
# END: Convert an aa fasta file to a 3di fasta file
#######################################################


#######################################################
# Re-align the AAs with USalign
#######################################################
cd /GitHub/str2phy/ex/ferritins_M20/align/03_USalign_v1
mkdir chains
cp /GitHub/str2phy/ex/ferritins_M20/rawdata/*.pdb chains
cd chains
ls *.pdb > chainslist.txt
mv chainslist.txt ..
cd /GitHub/str2phy/ex/ferritins_M20/align/03_USalign_v1
head chainslist.txt


# -i  Use alignment specified by 'align.txt' - NOTE: doesn't work with -mm (multiple structure alignment)
USalign -dir chains/ chainslist.txt -mol prot -mm 4 -o ferritins_pymol | tee USalign_ferritins_so1.txt &


# Remove the header junk after ":"
cut -d : -f 1 USalign_53_ferritins_aln.fasta > USalign_53_ferritins_aln2.fasta


# In R:
#######################################################
# Convert an aa fasta file to a 3di fasta file
#######################################################
library(ape)
library(seqinr)
library(BioGeoBEARS)
sourceall("/GitHub/bioinfRhints/Rsrc")
source("/GitHub/str2phy/Rsrc/str2phy_v1.R")

wd = "/GitHub/str2phy/ex/ferritins_M20/align/03_USalign_v1/"
setwd(wd)

aligned_fn = "USalign_53_ferritins_aln2.fasta"
unaligned_fn = "53_ferritin_3dis.fasta"
outfn = paste0(unaligned_fn, ".aln")
aligned_AAs = align_3dis_to_AAs(aligned_fn, unaligned_fn, outfn)
moref(outfn)
#######################################################
# END: Convert an aa fasta file to a 3di fasta file
#######################################################


#######################################################
# Reorder, according to an early phylogeny
#######################################################
library(ape)
library(seqinr)
library(BioGeoBEARS)
sourceall("/GitHub/bioinfRhints/Rsrc")
source("/GitHub/str2phy/Rsrc/str2phy_v1.R")

#wd = "/GitHub/str2phy/ex/ferritins_M20/02_FAMSA_3dis_aln_v1/z_FAMSA2.2.2_plain_MIQS_matrix/"
#setwd(wd)

trfn = "/GitHub/str2phy/ex/ferritins_M20/02_FAMSA_3dis_aln_v1/z_FAMSA2.2.2_plain_MIQS_matrix/53_ferritin_both.fasta.aln.contree.newick"
tr = read.tree(trfn)
tr$tip.label = gsub(pattern="'", replacement="", x=tr$tip.label)
cat(tr$tip.label, sep="\n")


fasta_fn = "53_ferritin_AAs.fasta.aln"
outfn=NULL
type="AA"
tip_txt = tr$tip.label
outfn = gsub(pattern="\\.fasta", replacement="", x=fasta_fn)
outfn = paste0(outfn, "_reord.fasta")
alignment = reorder_fasta(fasta_fn, tip_txt=tip_txt, outfn=outfn, type="AA")
alignment
moref(outfn)


#######################################################
# Trim the 3di alignment
#######################################################
cd /GitHub/str2phy/ex/ferritins_M20/02_FAMSA_3dis_aln_v1/

trimal -in 53_ferritin_3dis.aln_reord.fasta -out 53_ferritin_3dis.aln_reord_trim35.fasta -gt 0.35 -colnumbering | tee 53_ferritin_3dis.aln_reord_trim35_cols.txt

trimal -in 53_ferritin_AAs.aln_reord.fasta -out 53_ferritin_AAs.aln_reord_trim35.fasta -gt 0.35 -colnumbering | tee 53_ferritin_AAs.aln_reord_trim35_cols.txt


#######################################################
# PRACTICE, DISCARD
#######################################################
cd /GitHub/str2phy/ex/ferritins_M20/02_FAMSA_3dis_aln_v1/

trimal -in 53_ferritin_3dis.aln_reord.fasta -out 53_ferritin_3dis.aln_reord_trimAuto1.fasta -automated1 -matrix matrix.3DI -colnumbering

trimal -in 53_ferritin_3dis.aln_reord.fasta -out 53_ferritin_3dis.aln_reord_trimAuto1.fasta -automated1 -colnumbering


trimal -in 53_ferritin_3dis.aln_reord.fasta -out 53_ferritin_3dis.aln_reord_trimStrict.fasta -strict -matrix matrix.3DI -colnumbering

trimal -in 53_ferritin_3dis.aln_reord.fasta -out 53_ferritin_3dis.aln_reord_trimStrict.fasta -strict -colnumbering


# DOESNT WORK
#trimal -in 53_ferritin_3dis.aln_reord.fasta -out 53_ferritin_3dis.aln_reord_trimGapOut.fasta -gappyout -matrix matrix.3DI -colnumbering

trimal -in 53_ferritin_3dis.aln_reord.fasta -out 53_ferritin_3dis.aln_reord_trimStrictP.fasta -strictplus -matrix matrix.3DI -colnumbering

trimal -in 53_ferritin_3dis.aln_reord.fasta -out 53_ferritin_3dis.aln_reord_trimStrictP.fasta -strictplus -colnumbering


trimal -in 53_ferritin_3dis.aln_reord.fasta -out 53_ferritin_3dis.aln_reord_trimGt50.fasta -gt 0.5 -matrix matrix.3DI -colnumbering

trimal -in 53_ferritin_3dis.aln_reord.fasta -out 53_ferritin_3dis.aln_reord_trimGt50.fasta -gt 0.5 -colnumbering


trimal -in 53_ferritin_3dis.aln_reord.fasta -out 53_ferritin_3dis.aln_reord_trimGt50.fasta -st 0.1 -colnumbering

trimal -in 53_ferritin_3dis.aln_reord.fasta -out 53_ferritin_3dis.aln_reord_trimGt50.fasta -st 0.1 -matrix matrix.BLOSUM62 -colnumbering

trimal -in 53_ferritin_3dis.aln_reord.fasta -out 53_ferritin_3dis.aln_reord_trimGt50.fasta -st 0.1 -matrix matrix.3DI -colnumbering


#######################################################
# Manually cut famsa alignment to conserved cores?
# Not for now...
#######################################################
53_ferritin_3dis.aln_reord_trim35.fasta

# In R:
#######################################################
# Horizontally concatenate (hcat) an AA fasta file to a 3di fasta file
#######################################################
library(ape)
library(seqinr)
library(BioGeoBEARS)
sourceall("/GitHub/bioinfRhints/Rsrc")
source("/GitHub/str2phy/Rsrc/str2phy_v1.R")

wd = "/GitHub/str2phy/ex/ferritins_M20/02_FAMSA_3dis_aln_v1/"
setwd(wd)

fasta_fn1 = "53_ferritin_AAs.aln_reord.fasta"
fasta_fn2 = "53_ferritin_3dis.aln_reord.fasta"
outfn = "53_ferritin_both.fasta.aln"
aa3di_alignment = hcat_fastas(fasta_fn1, fasta_fn2, outfn=outfn)
moref(outfn)


fasta_fn1 = "53_ferritin_AAs.aln_reord_trim35.fasta"
fasta_fn2 = "53_ferritin_3dis.aln_reord_trim35.fasta"
outfn = "53_ferritin_both_trim35.fasta.aln"
aa3di_alignment = hcat_fastas(fasta_fn1, fasta_fn2, outfn=outfn)
moref(outfn)
```
