## Supplementary material for "Tertiary-interaction characters enable fast, model-based structural phylogenetics beyond the twilight zone": 00_pdbs_to_3di_v1.R

```
#######################################################
# PDB structures to AA & 3di sequences
#######################################################
#library(bio3d)
library(BioGeoBEARS)
source("/GitHub/str2phy/Rsrc/str2phy_v1.R")


wd = "/GitHub/str2phy/ex/ferritins_M20af/rawdata"
setwd(wd)

pdb_fns = list.files(path=".", pattern="\\.pdb")
length(pdb_fns)
# 53

fns_df = str_to_3di(str_fn=pdb_fns, suffix="\\.pdb")
fns_df

fns_fn = "53_ferritin_pdbs_converted_to_AA_3di.txt"
write.table(fns_df, fns_fn, quote=FALSE, sep="\t")
moref(fns_fn)


# Make a fasta file of the AAs
fasta_fns = fns_df$di3_fn

cd /GitHub/str2phy/ex/ferritins_M20af/rawdata/
# ls -1 *_aa.fasta | while read fn ; do cat "$fn" >> 53_ferritin_AAs.fasta; done
cat *_aa.fasta > 53_ferritin_alphafolds_AAs.fasta
cat *_3di.fasta > 53_ferritin_alphafolds_3dis.fasta

head 53_ferritin_alphafolds_AAs.fasta
head 53_ferritin_alphafolds_3dis.fasta

cp 53_ferritin_alphafolds_AAs.fasta 53_ferritin_alphafolds_AAs_ORIG.fasta
cp 53_ferritin_alphafolds_3dis.fasta 53_ferritin_alphafolds_3dis_ORIG.fasta

cut -d"_" -f1 -f2 53_ferritin_alphafolds_AAs_ORIG.fasta > 53_ferritinAFs_AAs.fasta
open 53_ferritinAFs_AAs.fasta
cut -d"_" -f1 -f2 53_ferritin_alphafolds_3dis_ORIG.fasta > 53_ferritinAFs_3dis.fasta
open 53_ferritinAFs_3dis.fasta

# Reorder according to tree order

#######################################################
# Reorder the USaln of alphafold-derived sequences
#######################################################
library(ape)
library(seqinr)
library(BioGeoBEARS)
sourceall("/GitHub/bioinfRhints/Rsrc")
source("/GitHub/str2phy/Rsrc/str2phy_v1.R")

wd = "/GitHub/str2phy/ex/ferritins_M20af/rawdata/"
setwd(wd)

trfn = "/GitHub/str2phy/ex/ferritins_M20/02_FAMSA_3dis_aln_v1/z_old/z_FAMSA2.2.2_plain_MIQS_matrix/53_ferritin_both.fasta.aln.contree.newick"
tr = read.tree(trfn)
tr$tip.label = gsub(pattern="'", replacement="", x=tr$tip.label)
tr$tip.label = gsub(pattern="\\.pdb", replacement="", x=tr$tip.label)
cat(tr$tip.label, sep="\n")


fasta_fn = "53_ferritinAFs_AAs.fasta"

outfn=NULL
type="AA"

tip_txt = tr$tip.label

outfn = gsub(pattern="\\.fasta", replacement="", x=fasta_fn)
outfn = paste0(outfn, "_reord.fasta")

alignment = reorder_fasta(fasta_fn, tip_txt=tip_txt, outfn=outfn, type="AA")
alignment
moref(outfn)


fasta_fn = "53_ferritinAFs_3dis.fasta"

outfn=NULL
type="AA"

tip_txt = tr$tip.label

outfn = gsub(pattern="\\.fasta", replacement="", x=fasta_fn)
outfn = paste0(outfn, "_reord.fasta")

alignment = reorder_fasta(fasta_fn, tip_txt=tip_txt, outfn=outfn, type="AA")


#######################################################
# USalign run
#######################################################
# Put the chains in a "chains" directory
mkdir chains
cp *.pdb chains
rm chainslist.txt

# DO NOT call the new file "list_of_chains.txt", 
# this cat command loops & fills up hard drive!!
cat *_chains.txt > chainslist.txt  
head chainslist.txt

USalign -dir chains/ chainslist.txt -mol prot -mm 4 -o ferritins_pymol | tee USalign_ferritins_so1.txt &

open USalign_ferritins_so1.txt

# Edit in BBedit, re-save to:
usalign_ferritins_aa.fasta
# #Total CPU time is 17.15 seconds

iqtree -t BIONJ -s usalign_ferritins_aa.fasta -m LG+F+G --ufboot 1000 -bnni | tee iqtree_ferritins_aa.so1.txt &
```
