## Supplementary material for "Tertiary-interaction characters enable fast, model-based structural phylogenetics beyond the twilight zone": __cmds_ferritins_M20af_famsa3di_or_USaln_then_trim_iqtree_v1.R

```
#######################################################
# Let's try FAMSA
# https://github.com/refresh-bio/FAMSA
#
#
# FAMSA: Fast and accurate multiple sequence alignment of ...
# Nature Journal
# S Deorowicz · 2016 
#######################################################

cd /GitHub/FAMSA
make
cp famsa /usr/local/bin


#######################################################
# Align the 3dis with FAMSA
#######################################################
cd /GitHub/str2phy/ex/ferritins_M20af/rawdata/
cp 53_ferritinAFs_AAs_reord.fasta /GitHub/str2phy/ex/ferritins_M20af/align/02_FAMSA_aln_v1
cp 53_ferritinAFs_3dis_reord.fasta /GitHub/str2phy/ex/ferritins_M20af/align/02_FAMSA_aln_v1


cd /GitHub/str2phy/ex/ferritins_M20af/align/02_FAMSA_aln_v1/
#famsa 53_ferritinAFs_3dis.fasta 53_ferritinAFs_3dis.fasta.aln
famsa3di 53_ferritinAFs_3dis_reord.fasta 53_ferritinAFs_3dis_famsa3di.fasta

# View in AliView
# Save to Fasta


# Convert the AAs to match 3di alignment

# In R:
#######################################################
# Convert an aa fasta file to a 3di fasta file
#######################################################
library(ape)
library(seqinr)
library(BioGeoBEARS)
sourceall("/GitHub/bioinfRhints/Rsrc")
source("/GitHub/str2phy/Rsrc/str2phy_v1.R")

wd = "/GitHub/str2phy/ex/ferritins_M20af/align/02_FAMSA_aln_v1/"
setwd(wd)

aligned_fn = "53_ferritinAFs_3dis_famsa3di.fasta"
unaligned_fn = "53_ferritinAFs_AAs_reord.fasta"
outfn = "53_ferritinAFs_AAs_famsa3di.fasta"
aligned_AAs = align_3dis_to_AAs(aligned_fn, unaligned_fn, outfn)
moref(outfn)
#######################################################
# END: Convert an aa fasta file to a 3di fasta file
#######################################################


# In R:
#######################################################
# Horizontally concatenate (hcat) an AA fasta file to a 3di fasta file
#######################################################
library(ape)
library(seqinr)
library(BioGeoBEARS)
sourceall("/GitHub/bioinfRhints/Rsrc")
source("/GitHub/str2phy/Rsrc/str2phy_v1.R")

wd = "/GitHub/str2phy/ex/ferritins_M20af/align/02_FAMSA_aln_v1/"
setwd(wd)

fasta_fn1 = "53_ferritinAFs_AAs_famsa3di.fasta"
fasta_fn2 = "53_ferritinAFs_3dis_famsa3di.fasta"
outfn = "53_ferritinAFs_both_famsa3di.fasta"
aa3di_alignment = hcat_fastas(fasta_fn1, fasta_fn2, outfn=outfn)
moref(outfn)

# PDB: 1496 columns, 748 per dataset
# Alphafold 3di aligned with famsa3di: 1548 columns, 774 each
#######################################################
# END: Horizontally concatenate (hcat) an AA fasta file to a 3di fasta file
#######################################################


#######################################################
# Trim the FAMSA3di alignments of alphafold structures
#######################################################

cd /GitHub/str2phy/ex/ferritins_M20af/align/02_FAMSA_aln_v1/
trimal -in 53_ferritinAFs_3dis_famsa3di.fasta -out 53_ferritinAFs_3dis_famsa3di_trim35.fasta -gt 0.35 -colnumbering | tee 53_ferritinAFs_3dis_famsa3di_trim35_cols.txt

cd /GitHub/str2phy/ex/ferritins_M20af/align/02_FAMSA_aln_v1/
trimal -in 53_ferritinAFs_AAs_famsa3di.fasta -out 53_ferritinAFs_AAs_famsa3di_trim35.fasta -gt 0.35 -colnumbering | tee 53_ferritinAFs_AAs_famsa3di_trim35_cols.txt

cd /GitHub/str2phy/ex/ferritins_M20af/align/02_FAMSA_aln_v1/
trimal -in 53_ferritinAFs_both_famsa3di.fasta -out 53_ferritinAFs_both_famsa3di_trim35.fasta -gt 0.35 -colnumbering | tee 53_ferritinAFs_both_famsa3di_trim35_cols.txt
# 462 positions, 231 per


 
 
 
#######################################################
# alphafold structures -> 3di alignment with famsa3di --> iqtree
#######################################################


#######################################################
# Single-dataset (AA or 3di) runs
#######################################################

# Trim35 = only keep columns with 35% or higher sites 
cd /GitHub/str2phy/ex/ferritins_M20af/02_FAMSA_3dis_aln_v1/iqtree_AAsTrim35/
iqtree -s 53fer.fasta -mset Blosum62,Dayhoff,DCMut,JTT,JTTDCMut,LG,Poisson,Poisson+FQ,Poisson,PMB,WAG,EX2,EX3,EHO,EX_EHO,3DI -mfreq FU,F -mrate E,G,R --ufboot 1000 -alrt 1000 -bnni --redo | tee 53fer_so1.txt &

cd /GitHub/str2phy/ex/ferritins_M20af/02_FAMSA_3dis_aln_v1/iqtree_AAsTrim35_allmodels/
iqtree -s 53fer.fasta -madd 3DI -mdef 3DI.nexus  --ufboot 1000 -alrt 1000 -bnni --redo | tee 53fer_so1.txt &


cd /GitHub/str2phy/ex/ferritins_M20af/02_FAMSA_3dis_aln_v1/iqtree_3disTrim35/
iqtree -s 53fer.fasta -mset Blosum62,Dayhoff,DCMut,JTT,JTTDCMut,LG,Poisson,Poisson+FQ,Poisson,PMB,WAG,EX2,EX3,EHO,EX_EHO,3DI -mfreq FU,F -mrate E,G,R --ufboot 1000 -alrt 1000 -bnni --redo | tee 53fer_so1.txt &


cd /GitHub/str2phy/ex/ferritins_M20af/02_FAMSA_3dis_aln_v1/iqtree_3disTrim35_allmodels/
iqtree -s 53fer.fasta -madd 3DI -mdef 3DI.nexus  --ufboot 1000 -alrt 1000 -bnni --redo | tee 53fer_so1.txt &


# Full single AAs or 3dis -- no trimming
cd /GitHub/str2phy/ex/ferritins_M20af/02_FAMSA_3dis_aln_v1/iqtree_AAsFull/
iqtree -s 53fer.fasta -mset Blosum62,Dayhoff,DCMut,JTT,JTTDCMut,LG,Poisson,Poisson+FQ,Poisson,PMB,WAG,EX2,EX3,EHO,EX_EHO,3DI -mfreq FU,F -mrate E,G,R --ufboot 1000 -alrt 1000 -bnni --redo | tee 53fer_so1.txt &

cd /GitHub/str2phy/ex/ferritins_M20af/02_FAMSA_3dis_aln_v1/iqtree_AAsFull_allmodels/
iqtree -s 53fer.fasta -madd 3DI -mdef 3DI.nexus  --ufboot 1000 -alrt 1000 -bnni --redo | tee 53fer_so1.txt &


cd /GitHub/str2phy/ex/ferritins_M20af/02_FAMSA_3dis_aln_v1/iqtree_3disFull/
iqtree -s 53fer.fasta -mset Blosum62,Dayhoff,DCMut,JTT,JTTDCMut,LG,Poisson,Poisson+FQ,Poisson,PMB,WAG,EX2,EX3,EHO,EX_EHO,3DI -mfreq FU,F -mrate E,G,R --ufboot 1000 -alrt 1000 -bnni --redo | tee 53fer_so1.txt &

cd /GitHub/str2phy/ex/ferritins_M20af/02_FAMSA_3dis_aln_v1/iqtree_3disFull_allmodels/
iqtree -s 53fer.fasta -madd 3DI -mdef 3DI.nexus  --ufboot 1000 -alrt 1000 -bnni --redo | tee 53fer_so1.txt &


# Trim35 FAMSA 3di alignments on BOTH AA+3di, no partitioning

cd /GitHub/str2phy/ex/ferritins_M20af/02_FAMSA_3dis_aln_v1/iqtree_BothTrim35_NonPartitioned/
iqtree -s 53fer.fasta -mset Blosum62,Dayhoff,DCMut,JTT,JTTDCMut,LG,Poisson,Poisson+FQ,Poisson,PMB,WAG,EX2,EX3,EHO,EX_EHO,3DI -mfreq FU,F -mrate E,G,R --ufboot 1000 -alrt 1000 -bnni --redo | tee 53fer_so1.txt &

cd /GitHub/str2phy/ex/ferritins_M20af/02_FAMSA_3dis_aln_v1/iqtree_BothTrim35_NonPartitioned_allmodels/
iqtree -s 53fer.fasta -madd 3DI -mdef 3DI.nexus --ufboot 1000 -alrt 1000 -bnni --redo | tee 53fer_so1.txt &

cd /GitHub/str2phy/ex/ferritins_M20af/02_FAMSA_3dis_aln_v1/iqtree_BothTrim35/
iqtree -s 53fer.fasta -spp partitions_v1.raxml -m MFP+MERGE -madd 3DI -mdef 3DI.nexus -mset Blosum62,Dayhoff,DCMut,JTT,JTTDCMut,LG,Poisson,Poisson+FQ,Poisson,PMB,WAG,EX2,EX3,EHO,EX_EHO,3DI -mfreq FU,F -mrate E,G,R --ufboot 1000 -alrt 1000 -bnni --redo | tee 53fer_so1.txt &

cd /GitHub/str2phy/ex/ferritins_M20af/02_FAMSA_3dis_aln_v1/iqtree_BothTrim35_allmodels/
iqtree -s 53fer.fasta -spp partitions_v1.raxml -m MFP+MERGE -madd 3DI -mdef 3DI.nexus --ufboot 1000 -alrt 1000 -bnni --redo | tee 53fer_so1.txt &


# Full FAMSA 3di alignments on BOTH AA+3di, with partitioning
cd /GitHub/str2phy/ex/ferritins_M20af/02_FAMSA_3dis_aln_v1/iqtree_BothFull_NonPartitioned/
iqtree -s 53fer.fasta -mset Blosum62,Dayhoff,DCMut,JTT,JTTDCMut,LG,Poisson,Poisson+FQ,Poisson,PMB,WAG,EX2,EX3,EHO,EX_EHO,3DI -mfreq FU,F -mrate E,G,R --ufboot 1000 -alrt 1000 -bnni --redo | tee 53fer_so1.txt &

cd /GitHub/str2phy/ex/ferritins_M20af/02_FAMSA_3dis_aln_v1/iqtree_BothFull_NonPartitioned_allmodels/
iqtree -s 53fer.fasta -madd 3DI -mdef 3DI.nexus --ufboot 1000 -alrt 1000 -bnni --redo | tee 53fer_so1.txt &

cd /GitHub/str2phy/ex/ferritins_M20af/02_FAMSA_3dis_aln_v1/iqtree_BothFull/
iqtree -s 53fer.fasta -spp partitions_v1.raxml -m MFP+MERGE -madd 3DI -mdef 3DI.nexus -mset Blosum62,Dayhoff,DCMut,JTT,JTTDCMut,LG,Poisson,Poisson+FQ,Poisson,PMB,WAG,EX2,EX3,EHO,EX_EHO,3DI -mfreq FU,F -mrate E,G,R --ufboot 1000 -alrt 1000 -bnni --redo | tee 53fer_so1.txt &

cd /GitHub/str2phy/ex/ferritins_M20af/02_FAMSA_3dis_aln_v1/iqtree_BothFull_allmodels/
iqtree -s 53fer.fasta -spp partitions_v1.raxml -m MFP+MERGE -madd 3DI -mdef 3DI.nexus --ufboot 1000 -alrt 1000 -bnni --redo | tee 53fer_so1.txt &
```
