## Supplementary material for "Tertiary-interaction characters enable fast, model-based structural phylogenetics beyond the twilight zone": _02_download_alphafolds_loop_v2.R

```
#######################################################
# Downloading alphafold .cif files in a loop
# (closest match to a series of FASTA entries)
#######################################################

library(ape)
library(msa)
library(seqinr)
library(BioGeoBEARS)
library(rvest) # for html_table()

sourceall("/GitHub/bioinfRhints/Rsrc/")
source("/GitHub/str2phy/Rsrc/str2phy_v1.R")


wd = "/GitHub/str2phy/ex/ferritins_M20/alphafolds"
setwd(wd)

fasta_fn = "53_ferritin_AAs.fasta"
tmpseqs = ape::read.FASTA(fasta_fn, type="AA")
numseqs = length(tmpseqs)
numseqs

outdfs = NULL; i=1

# outdfs = outdfs[1:124,]
#exclude_is = c(125,152, 159, 160)
exclude_is = c()
#for (i in 804:804)
for (i in 1:numseqs)
#exclude_is = c()
#for (i in 1:numseqs)
	{
	txt = paste0("Getting alphafold structure for sequence #", i, "/", numseqs)
	cat("\n")
	cat(txt)
	cat("\n")
	
	if ((i %in% exclude_is) == TRUE)
		{
		seqs = ape::read.FASTA(fasta_fn, type="AA")
		header = names(seqs)[[i]]
		gid = firstword(header)
		
		txt = paste0("Sequence #", i, "/", numseqs, ", aka '", gid, "', skipped as it is known to cause a problem.")
		cat("\n")
		cat(txt)
		cat("\n")
		outdf = rep(NA, times=13)
		names(outdf) = c("seqnum", "AlphaFold Model", "Query Sequence", "Identity %", "Coverage %", "cif_fn", "html_fn", "table_fn", "dump_fn", "chain_names_fn", "di3_fn", "aa_fn", "both_fn")
		outdfs = rbind(outdfs, outdf)
		next()
		} # END if (i %in% exclude_is)
	
	outdf = fasta_to_alphafold(fasta_fn=fasta_fn, seqnum=i)
	print(outdf[1:5])
	outdfs = rbind(outdfs, outdf)
	outdfs_fn = "3281_BRDs_alphafolds_pt03partial.txt"
	write.table(x=outdfs, file=outdfs_fn, quote=FALSE, sep="\t", row.names=FALSE, col.names=TRUE)
	}

dim(outdfs)

outdfs_fn = "53_ferritin_alphafolds.txt"
write.table(x=outdfs, file=outdfs_fn, quote=FALSE, sep="\t", row.names=FALSE, col.names=TRUE)

#outdfs_fn = "3281_BRDs_alphafolds_WORKED_BKUP.txt"
#write.table(x=outdfs, file=outdfs_fn, quote=FALSE, sep="\t", row.names=FALSE, col.names=TRUE)


# -mm 4         -- 4: alignment of multiple monomeric chains into a consensus alignment
# -ter 2        -- 2: (default) only align the first chain
# -TMscore 0 	  -- 0: (default) sequence independent structure alignment

# Put all the matching filenames into a text file
cp *.cif chains
ls chains/*.cif > list_of_chains.txt

# Add :A to each line, as these are all A chains
#sed -i '' 's/$/:A/'  list_of_chains.txt
# Remove chains/chains/ to chains/
sed -i '' 's/chains\///'  list_of_chains.txt
cp list_of_chains.txt list_of_chains_orig.txt
wc -l list_of_chains_orig.txt
head -100 list_of_chains_orig.txt > list_of_chains.txt
wc -l list_of_chains.txt
head list_of_chains.txt

tail list_of_chains.txt

USalign -dir chains/ list_of_chains.txt -mol prot -mm 4 | tee usalign_test1.txt &

head -14 usalign_test1.txt
wc -l usalign_test1.txt
tail -202 usalign_test1.txt > usalign_test2.fasta

head usalign_test2.fasta
tail usalign_test2.fasta
wc -l usalign_test2.fasta

head -200 usalign_test2.fasta > usalign_test3.fasta
tail usalign_test3.fasta
wc -l usalign_test3.fasta
```
