## Supplementary material for "Tertiary-interaction characters enable fast, model-based structural phylogenetics beyond the twilight zone": plotConfidence.R

```
library(seqinr)


# Confidence distribution per structure
confidence.df = data.frame(structure = character(0), lower = numeric(0), mean = numeric(0), upper = numeric(0))


# Confidence per site
alignment = read.fasta("align/USalign_53_ferritinAFs_AAs_FULL.fasta.aln")
names(alignment) = gsub("[:].+", "", names(alignment))
nsites = length(alignment[[1]])
alignment.scores = list()
for (i in 1:nsites){
	alignment.scores[[i]] = numeric(0)
}


# All pdb structures
pdbs = list.files("rawdata", pattern=".+[.]pdb")

for (pdb in pdbs){


	acc = pdb# strsplit(pdb, "_")[[1]][1]
	contents = readLines(paste0("rawdata/", pdb))
	contents = contents[grep("^ATOM", contents)]
	contents = contents[substr(contents, 14, 15) == "CA"] # Alpha carbon only
	confidence = as.numeric(substr(contents, 62, 66))

	summ = summary(confidence)
	lower = as.numeric(quantile(confidence, 0.1))
	m = mean(confidence)
	upper = as.numeric(quantile(confidence, 0.9))

	confidence.df2 = data.frame(structure = acc, lower = lower, mean = m, upper = upper)
	confidence.df = rbind(confidence.df, confidence.df2)


	# Step through sequence
	aln.seq = as.character(alignment[[pdb]])
	npos = sum(aln.seq != "-")

	if (npos != length(confidence)){
		cat(paste0("Warning: aligned length is not same as pdb length", pdb, "\n"))
		next
	}


	# For each non-gap site
	ungapped.pos = 1
	for (site in 1:nsites){
		char = aln.seq[site]
		if (char == "-"){
			next
		}

		alignment.scores[[site]] = c(alignment.scores[[site]], confidence[ungapped.pos])

		ungapped.pos = ungapped.pos + 1


	}


}


write.table(confidence.df, "confidence.tsv", sep="\t", quote=F, row.names=F)


cat(paste0("The average structure has a mean pLDDT of ", signif(mean(confidence.df$mean), 3), "%, with 90% of all sites scoring over ", signif(mean(confidence.df$lower), 3), "%.\n"))


# nrows = sapply(alignment.scores, length)
# confidence.per.site.mean = sapply(alignment.scores, mean)
# confidence.per.site.lower = sapply(alignment.scores, function(ele) as.numeric(quantile(ele, 0.25)) )
# confidence.per.site.upper = sapply(alignment.scores, function(ele) as.numeric(quantile(ele, 0.75)) )
# keep = which(nrows>length(pdbs)/2)


# confidence.per.site.mean[-keep] = 0
# confidence.per.site.lower[-keep] = 0
# confidence.per.site.upper[-keep] = 0


# plot(0, 0, type="n", xlim=c(0, nsites+1), ylim=c(0, 100), xlab="Alignment position", ylab="pLDDT score")
# polygon(c(1:nsites, nsites:1), c(confidence.per.site.lower, confidence.per.site.upper), col="#008cba")
```
