## Supplementary material for "Tertiary-interaction characters enable fast, model-based structural phylogenetics beyond the twilight zone": 01_change_AAs_to_3Dis_in_PDB_v1.R

```
library(ape)
library(seqinr)
library(stringr)
library(openxlsx)

wd = "/GitHub/3diphy/jalview/change_colors/"
setwd(wd)


# Chain used
chain_name = "2jd7_A"

# Translate positions of PDB chain to AA alignment
xlsfn = "2jd7_A_match_AAs_to_PDB.xlsx"

#######################################################
# Read the chain to a table
#######################################################
fn = "2jd7_A_AAs_to_3Di.pdb"
lines = readLines(fn)

fn2 = "tmp_pdb.pdb"
writeLines(text=lines[1:(length(lines)-1)], con=fn2)

dtf = read.table(fn2, skip=1)
pdb_AAAs = dtf[,4]
pdb_AAs = seqinr::a(stringr::str_to_title(pdb_AAAs))
pdb_AAs

# Unique AA positions
uniq_positions = unique(dtf$V6)

first_line_for_new_AAs = match(uniq_positions, table=dtf$V6)
first_line_for_new_AAs

pdb_AAs = pdb_AAs[first_line_for_new_AAs]

#######################################################
# Read the alignment
#######################################################
AAfn = "53_ferritinAFs_AAs_famsa3di_trim35.fasta"
di3fn = "53_ferritinAFs_3dis_famsa3di_trim35.fasta"

aln_aaBin = ape::read.FASTA(AAfn, type="AA")
aa_chars = as.character(aln_aaBin)

aln_di3_Bin = ape::read.FASTA(di3fn, type="AA")
di3_chars = as.character(aln_di3_Bin)


seq_names1 = names(aa_chars)
seq_names2 = names(di3_chars)

seq_names1 == seq_names2

TF = seq_names1 == chain_name
seqnum = (1:length(TF))[TF]
seqnum


aa_chars[[seqnum]]
di3_chars[[seqnum]]

aa_nogaps = aa_chars[[seqnum]][aa_chars[[seqnum]] != "-"]
aa_nogaps
di3_nogaps = di3_chars[[seqnum]][di3_chars[[seqnum]] != "-"]
di3_nogaps

cat(aa_nogaps, sep="\n")

cat(pdb_AAs, sep="\n")

pdb_AAs

# Load the position translation table
translate_table1 = openxlsx::read.xlsx(xlsfn)

# Double-check nothing is missing
test1 = translate_table1$AA[translate_table1$AA != "-"]
test2 = aa_nogaps
cbind(test1, test2)
# Works

# Substitute in the 3di characters
translate_table2 = translate_table1
translate_table2$AA[translate_table2$AA != "-"] = di3_nogaps

# Check
cbind(translate_table1, translate_table2)
tail(cbind(translate_table1, translate_table2), 15)

# Convert the 3di codes to 3-letter codes
di3_as_3letter_AA_codes = seqinr::aaa(translate_table2$AA)
di3_as_3letter_AA_codes
di3_as_3letter_AA_codes = toupper(di3_as_3letter_AA_codes)
di3_as_3letter_AA_codes
di3_as_3letter_AA_codes = di3_as_3letter_AA_codes[is.na(di3_as_3letter_AA_codes) == FALSE]

dtf2 = dtf

# Check that 3dis to substitute in are the same or longer than the AAs in the PDB
length(unique(dtf2$V6))
length(di3_as_3letter_AA_codes)

# Substitute back into PDB structure
dtf2 = dtf
for (i in 1:length(unique(dtf2$V6)))
	{
	TF = dtf2$V6 == i
	print(sum(TF))
	print(di3_as_3letter_AA_codes[i])
	dtf2$V4[TF] = di3_as_3letter_AA_codes[i]
	}

head(dtf)
head(dtf2)

tail(dtf)
tail(dtf2)


# Write back out to file
outfn = "2jd7_A_AAs_to_3Di_DONE.pdb"
write(lines[1], file=outfn, append=FALSE)

# Writing MUST happen with correct padding
spaces1 = 7 - nchar(dtf2$V2)
spaces2 = rep(2, times=length(dtf2$V2))
spaces3 = 4 - nchar(dtf2$V3)
spaces4 = rep(1, times=length(dtf2$V2))
spaces5 = 4 - nchar(dtf2$V6)
spaces6 = 12 - nchar(dtf2$V7)
spaces7 = 8 - nchar(dtf2$V8)
spaces8 = 8 - nchar(dtf2$V9)
spaces9 = rep(2, times=length(dtf2$V2))
spaces10 = rep(1, times=length(dtf2$V2))
spaces11 = rep(11, times=length(dtf2$V2))

for (i in 1:nrow(dtf2))
	{
	spaces1 = paste0(rep(" ", times=(7 - nchar(dtf2$V2[i]))), collapse="")
	spaces2 = "  "
	spaces3 = paste0(rep(" ", times=(4 - nchar(dtf2$V3[i]))), collapse="")
	spaces4 = " "
	spaces5 = paste0(rep(" ", times=(4 - nchar(dtf2$V6[i]))), collapse="")
	spaces6 = paste0(rep(" ", times=(12 - nchar(sprintf("%.3f", dtf2$V7[i])))), collapse="")
	spaces7 = paste0(rep(" ", times=(8 - nchar(sprintf("%.3f", dtf2$V8[i])))), collapse="")
	spaces8 = paste0(rep(" ", times=(8 - nchar(sprintf("%.3f", dtf2$V9[i])))), collapse="")
	spaces9 = "  "
	spaces10 = " "
	spaces11 = "           "
	
	tmpline = paste0(dtf2$V1[i], spaces1, 
	dtf2$V2[i], spaces2, 
	dtf2$V3[i], spaces3, 
	dtf2$V4[i], spaces4, 
	dtf2$V5[i], spaces5, 
	dtf2$V6[i], spaces6, 
	sprintf("%.3f", dtf2$V7[i]), spaces7, 
	sprintf("%.3f", dtf2$V8[i]), spaces8, 
	sprintf("%.3f", dtf2$V9[i]), spaces9, 
	sprintf("%.2f", dtf2$V10[i]), spaces10, 
	sprintf("%.2f", dtf2$V11[i]), spaces11, 
	dtf2$V12[i], collapse="")
	
	write(tmpline, file=outfn, append=TRUE)
	}

tmp = cbind(dtf$V1, spaces1, dtf$V2, spaces2, collapse=NULL)
apply(X=tmp, MAR=1, FUN=paste0, collapse="")

# NO
#write.table(x=dtf2, file=outfn, col.names=FALSE, row.names=FALSE, append=TRUE, quote=FALSE)
write(lines[length(lines)], file=outfn, append=TRUE)
```
