## Supplementary material for "Tertiary-interaction characters enable fast, model-based structural phylogenetics beyond the twilight zone": Puente-Lelievre_etal_2023_BIORXIV-2023-571181v1-Matzke.pdf

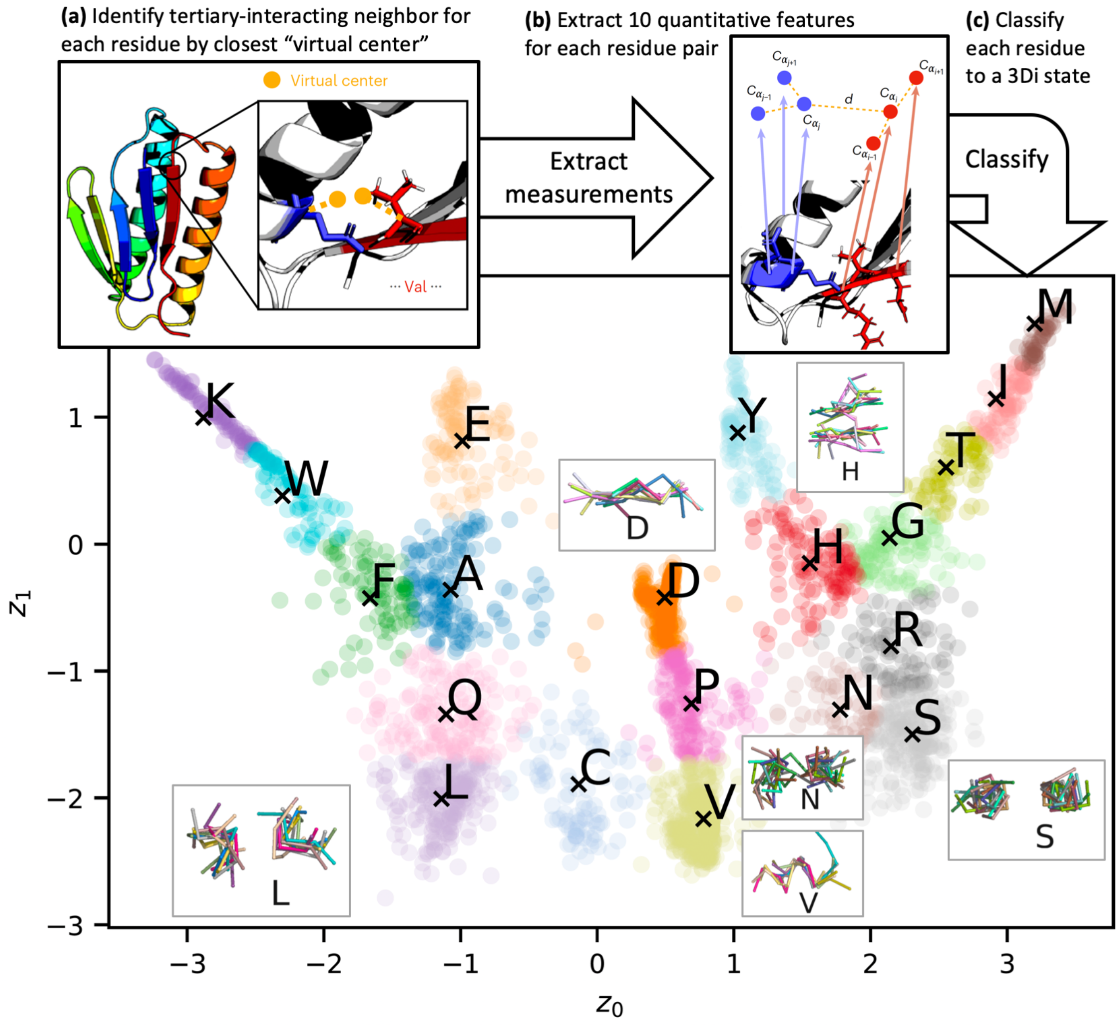


**Figure 1.** Summary of Foldseek's system for classifying tertiary interactions into the 20-state 3Di alphabet. For each residue in a protein structure **(a)** a virtual center is defined, and each residue is assigned a tertiary neighbor based on the closest virtual center. **(b)** 10 quantitative traits are collected for each residue+neighbor interaction, and then **(c)** assigned to the best-matching 3Di state using the classifier pre-trained by Van Kempen et al. 2023. The main panel shows a two-dimensional representation of the 20 states. Examples of 3Di states are inset; each of these represents 10 pairs of backbone fragments showing the typical arrangement for a 3Di state. Graphics modified from Van Kempen et al. 2023.

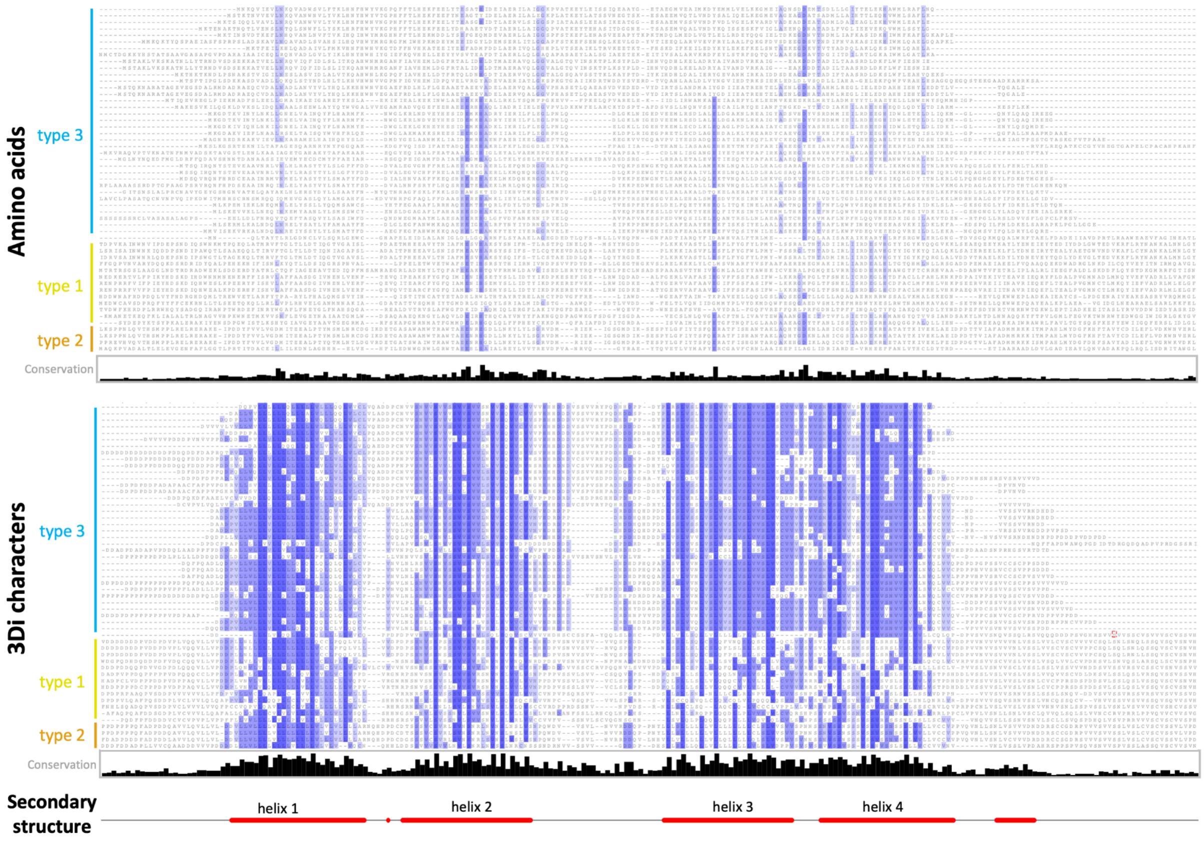


**Figure 2.** AA and 3Di alignments for 53 AlphaFold structures of the ferritin-like superfamily, displayed with Jalview's pairwise identity color scheme, where characters matching the majority character (if any) are colored, and darker colors are closer to 100% agreement. The 3Di alignment was generated with famsa3di; the AA alignment was generated by one-to-one replacement of 3Di states with AAs. Sites with <35% data were trimmed. FASTA files are available in SI.

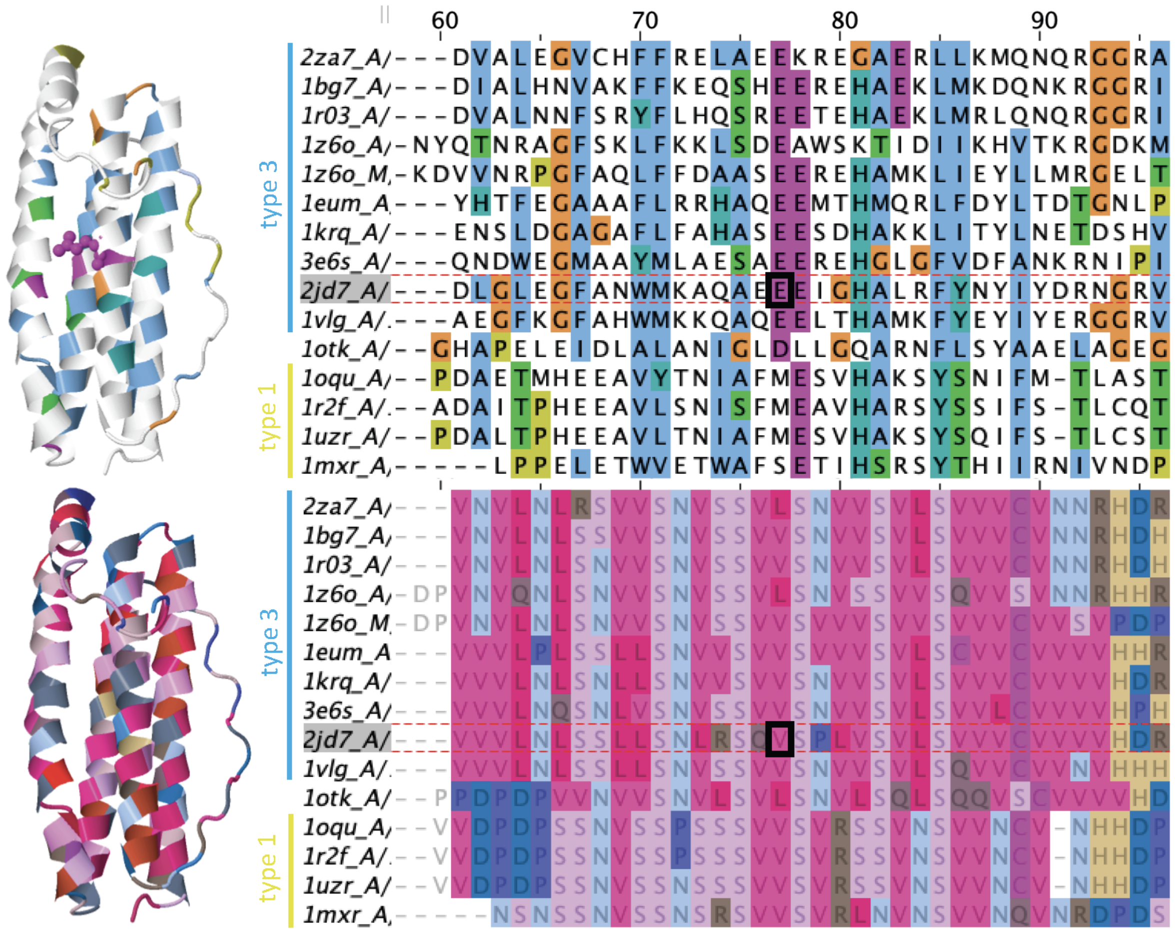


**Figure 3.** Illustrating the differences between AA and 3Di characters by zooming in on a portion of the alignment in Fig. 2, corresponding to the end of helix 2 and beginning of the subsequent loop. Glutamic acid E77 is displayed on the structure for reference. **Top:** AA alignment next to a ribbon diagram of PDB entry 2JD7:A (Tatur et al. 2007), both colored with the ClustalX color scheme (Jeanmougin et al. 1998). **Bottom:** 3Di alignment using Jalview's gecos-Sunset color scheme.

While both alignments have some repetitive structure in the depicted alpha-helical region, the 3Di characters are more conserved and contain information that is distinct from the AAs (Figure 3). For example, in the section of the complete alignment shown in Figure 3, sites 70-71 are both hydrophobic AAs, but do not show obvious grouping information between type 3 and type 1 groups. The corresponding 3Di states show a clear fixed difference at position 70 (V/L vs. S), and invariance at 71 (S). Sites 79-80 show little AA conservation, but the corresponding 3Di sites show fixed differences between the type 1 and type 3 groups. These "fixed" differences do not strictly hold for the full alignment, but type 3 vs. type 1/2 grouping is evident in the 3Di alignment (Figure 2).

*Model selection.* AA data are typically best fit by Blosum62 or LG, standard matrices for highly diverged AA sequences, or by EX3+FU, a mixture model containing three matrices modeling different transition rates and AA frequencies for exposed, intermediate, and buried sites (Le et al. 2008). Interestingly, LG is selected only for AA data aligned by a structure-free method. All 3Di alignments are best-fit by the 3Di substitution matrix with empirical base frequencies, with minor variations in the treatment of among-site rate variation (G4, R3-R5) (Table 1, Supporting Information).

**Table 1.** Best-fit substitution models for IQtree runs, according to log-likelihood (lnL) and BIC. IQtree inferences were conducted using the BIC-best model. Runs are sorted by deep & shallow similarity to the M20 tree (first line). Results are shown only for IQtree runs using the custom model list (with partitioning for AA+3Di datasets) which included the 3Di substitution matrix. *k* represents the number of free parameters in the substitution model; all runs also included 103 free branch-length parameters in the BIC calculation. Note: lnL & BIC values can only be compared within lines, not between them, as they are only comparable for identical alignments.

| Structure source | Alignment data | Alignment method | Phylogeny data | # bipartitions matched | | | Best log-likelihood models: | | | Best BIC models: | | | |
|---|---|---|---|---|---|---|---|---|---|---|---|---|---|
| | | | | all | deep | shal-low | lnL | *k* | Model | BIC | lnL | *k* | Model |
| PDB | structures | Superpose | *Q* score | 51 | 22 | 29 | | | | | | | |
| alphafold | struct.+AAs | USalign | AA+3di, trim | 40 | 20 | 20 | -24477.7 | 27 | EX3+FU+G; 3DI+F+R3 | 49728.3 | -24477.7 | 27 | EX3+FU+G; 3DI+F+R3 |
| PDB | 3di | famsa3di | AA+3di | 39 | 20 | 19 | -39984.6 | 48 | Blosum62+F+G4; 3DI+F+R5 | 81081.6 | -39984.6 | 48 | Blosum62+F+G4; 3DI+F+R5 |
| alphafold | struct.+AAs | USalign | AA+3di | 39 | 20 | 19 | -43208.6 | 47 | Blosum62+F+R3; 3DI+F+R3 | 87609.2 | -43208.6 | 47 | Blosum62+F+R3; 3DI+F+R3 |
| PDB | 3di | famsa3di | AA+3di, trim | 37 | 20 | 17 | -29156.4 | 41 | Blosum62+F+G4; 3DI+F+G4 | 59179.9 | -29156.4 | 41 | Blosum62+F+G4; 3DI+F+G4 |
| PDB | struct.+AAs | USalign | AA | 37 | 19 | 18 | -15160.7 | 25 | Blosum62+F+R4 | 30976.0 | -15208.7 | 4 | Blosum62+R3 |
| alphafold | 3di | famsa3di | AA+3di | 36 | 19 | 17 | -42057.8 | 44 | Blosum62+F+G4; 3DI+F+R3 | 85195.4 | -42057.8 | 44 | Blosum62+F+G4; 3DI+F+R3 |
| PDB | struct.+AAs | USalign | AA+3di | 34 | 19 | 15 | -40816.9 | 47 | Blosum62+F+R3; 3DI+F+R3 | 82772.5 | -40816.9 | 47 | Blosum62+F+R3; 3DI+F+R3 |
| alphafold | struct.+AAs | USalign | AA, trim | 38 | 18 | 20 | -16270.9 | 25 | Blosum62+F+R4 | 33187.1 | -16315.2 | 3 | EX3+FU+G |
| PDB | 3di | famsa3di | AA | 36 | 18 | 18 | -24783.6 | 25 | Blosum62+F+R4 | 50404.9 | -24792.0 | 20 | Blosum62+F+G4 |
| alphafold | struct.+AAs | USalign | AA | 36 | 18 | 18 | -27095.9 | 25 | Blosum62+F+R4 | 55106.9 | -27107.4 | 20 | Blosum62+F+G4 |
| PDB | struct.+AAs | USalign | AA | 36 | 18 | 18 | -24877.6 | 25 | Blosum62+F+R4 | 50631.3 | -24881.1 | 23 | Blosum62+F+R3 |
| PDB | struct.+AAs | USalign | AA+3di, trim | 33 | 18 | 15 | -23679.1 | 25 | Blosum62+R3; 3DI+F+G4 | 48115.0 | -23679.1 | 25 | Blosum62+R3; 3DI+F+G4 |
| alphafold | 3di | famsa3di | AA+3di, trim | 36 | 17 | 19 | -30459.0 | 27 | EX3+FU+G; 3DI+F+R3 | 61715.6 | -30459.0 | 27 | EX3+FU+G; 3DI+F+R3 |
| alphafold | 3di | famsa3di | 3di | 32 | 17 | 15 | -14950.2 | 25 | 3DI+F+R4 | 30745.2 | -14953.6 | 23 | 3DI+F+R3 |
| alphafold | struct.+AAs | USalign | 3di | 31 | 16 | 15 | -15985.7 | 25 | 3DI+F+R4 | 32886.3 | -15986.2 | 23 | 3DI+F+R3 |
| alphafold | 3di | famsa3di | AA | 32 | 14 | 18 | -26932.0 | 27 | Blosum62+F+R5 | 54691.6 | -26993.3 | 3 | EX3+FU+G |
| alphafold | 3di | famsa3di | 3di, trim | 31 | 14 | 17 | -10259.3 | 27 | 3DI+F+R5 | 21218.1 | -10260.7 | 25 | 3DI+F+R4 |
| PDB | 3di | famsa3di | AA, trim | 30 | 14 | 16 | -18633.2 | 25 | Blosum62+F+R4 | 37938.1 | -18641.4 | 20 | Blosum62+F+G4 |
| PDB | struct.+AAs | USalign | 3di, trim | 23 | 14 | 9 | -8173.4 | 25 | 3DI+F+R4 | 17006.9 | -8174.6 | 23 | 3DI+F+R3 |
| PDB | struct.+AAs | USalign | 3di | 22 | 14 | 8 | -15612.5 | 23 | 3DI+F+R3 | 32094.1 | -15612.5 | 23 | 3DI+F+R3 |
| alphafold | struct.+AAs | USalign | 3di, trim | 28 | 13 | 15 | -8037.0 | 25 | 3DI+F+R4 | 16740.9 | -8039.5 | 23 | 3DI+F+R3 |
| PDB | 3di | famsa3di | 3di, trim | 25 | 13 | 12 | -10215.4 | 25 | 3DI+F+R4 | 21107.5 | -10218.1 | 23 | 3DI+F+R3 |
| none | AAs | famsa | AA, trim | 30 | 12 | 18 | -24136.9 | 23 | LG+F+R3 | 49084.3 | -24139.0 | 20 | LG+F+G4 |
| alphafold | 3di | famsa3di | AA, trim | 29 | 12 | 17 | -20019.9 | 25 | LG+F+R4 | 40665.8 | -20044.4 | 3 | EX3+FU+G |
| PDB | 3di | famsa3di | 3di | 24 | 12 | 12 | -14851.9 | 25 | 3DI+F+R4 | 30553.0 | -14856.1 | 23 | 3DI+F+R3 |
| none | AAs | famsa | AA | 29 | 10 | 19 | -18766.9 | 25 | LG+F+R4 | 38190.2 | -18813.5 | 1 | LG+G4 |

Standard AA models fit best to AA data, with comparatively small differences between Blosum62, LG, and EX3. A much worse fit is the Poisson model, a Jukes-Cantor-style model which assumes that transition rates are equal between all residues. Finally, the least fitting model for AA data is the 3Di rate matrix. On the other hand, for 3Di data, the pattern is reversed, where the 3Di model dominates, followed by Poisson and then the AA models (Table 2).

**Table 2.** Illustrative models compared on AA and 3Di datasets. Alignment source: USalign on AlphaFold structures+AAs, trimmed sites with <35% data; phylogeny estimated with IQtree on custom model list including 3DI substitution matrix.

| Data | Model | lnL | ΔlnL | *k* | BIC |
|---|---|---|---|---|---|
| amino acids (*n* =191 sites) | EX3+FU+G | -16315.2 | 35.9 | 106 | 33187.1 |
| | Blosum62+F+G4 | -16279.2 | 0.0 | 123 | 33204.5 |
| | LG+F+G4 | -16289.1 | 9.8 | 123 | 33224.2 |
| | EX3+F+G | -16299.1 | 19.8 | 125 | 33254.6 |
| | Poisson+F+G4 | -16888.9 | 609.7 | 123 | 34423.9 |
| | 3DI+F+G4 | -17950.8 | 1671.5 | 123 | 36547.6 |
| 3Di characters (*n* =191 sites) | 3DI+F+G4 | -8054.9 | 0.0 | 123 | 16755.9 |
| | Poisson+F+G4 | -8329.4 | 274.5 | 123 | 17304.9 |
| | Blosum62+F+G4 | -8485.0 | 430.0 | 123 | 17616.0 |
| | EX3+F+G | -8620.9 | 566.0 | 125 | 17898.3 |
| | LG+F+G4 | -8734.1 | 679.1 | 123 | 18114.1 |
| | EX3+FU+G | -10415.9 | 2361.0 | 106 | 21388.6 |

*Phylogenetic analyses*. IQtree custom-model runs by the number of deep and shallow bipartitions that match the reference M20 tree are shown in Table 1. Results for all 60 ML runs are shown in Figure 4. Our results indicate that runs using both AA+3Di do best at matching the M20 tree topology, although with substantial overlap with structure-aligned AA runs. 3Di-only runs are consistently lower ranked.

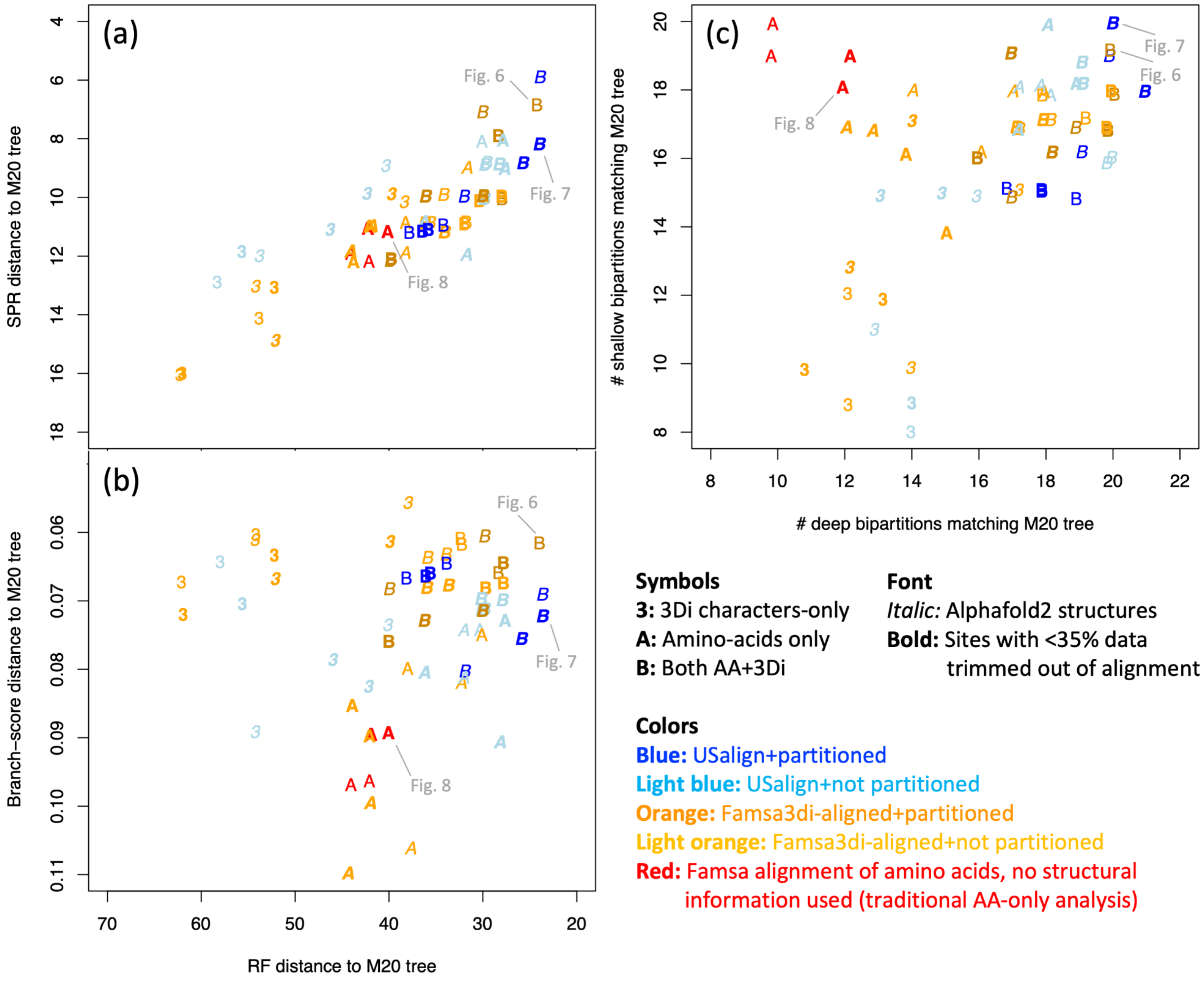


**Figure 4.** Similarity of all 60 IQtree runs to the Malik et al. (2020) structural-distances NJ tree (M20) for the ferritin-like superfamily. **(a)** Robinson-Foulds (RF) distance vs. Subtree Pruning-Regrafting (SPR) distance. SPR approximates the number of edits required to match the M20 topology. The closest matches to M20 on both metrics tend to be AA+3Di analyses with partitioning. AA-only and 3Di-only analyses tend to fare worse. **(b)** RF distance vs. branch-score distance, which takes branch lengths into account, unlike RF and SPR. 3Di-only match M20 well on the branch-score measure, but poorly on RF; however, AA+3Di performs well on both. **(c)** Number of M20 deep bipartitions matched vs. shallow bipartitions matched. "Deep" bipartitions are those annotated by Malik et al. to indicate major well-recognised categories, and groupings of those. Shallow bipartitions are all others, e.g. groups of 2-5 structures near the tips of the unrooted tree. AA-only runs retrieved shallow bipartitions but not deep ones; 3Di-only was weak on both, but AA+3Di retrieved both shallow and deep. *Note:* Points have been jittered for visibility.

Phylogenetic analyses using 3Di-only tend to be competitive in branch score difference. Structure-aligned AA-only runs are somewhat competitive in RF distance; only both AA+3Di runs are competitive in both (Figure 4b).

Amino-acid-only (structure-free) analyses are competitive for resolving shallow bipartitions, but are systematically poorer for deep bipartitions. 3Di-only analyses are poor for both. However, in combined analyses, the AA and 3Di data together seem to mutually compensate for the weaknesses of each individually and produce the best matching of both shallow and deep bipartitions (Figure 4c).

Also of interest are cases where differences in method do not seem to produce major differences in performance. It appears that PDB and AlphaFold structural data are equivalent in terms of phylogenetic results matching the M20 tree. Trimming, or not, also does not appear to make a major difference. Model partitioning did not seem to produce a consistent difference in terms of tree similarity to the M20 tree. However, lack of partitioning produces higher bootstrap supports. When bootstrap percent scores are summed across the tree for all 60 models (see SI), all 12 of the AA+3Di runs without partitioning were ranked in the top 15 (sum of bootstraps: 4219-4374). The next 16 runs contain 13 AA+3Di partitioned runs and 3 structure-aligned AA-only runs (sum of bootstraps: 4091-4216). The following 15 runs are all AA-only (3815-4087), with structure-free AA runs representing the last 4 (3815-3924), and the final 14 runs are all 3Di-only (3471-3799). While declining bootstraps for the noisier AA-only and 3Di-only datasets accords with expectations, elevated bootstraps for non-partitioned AA+3Di datasets probably represents artificial elevation of confidence due to a single model poorly fitting two very different partitions.

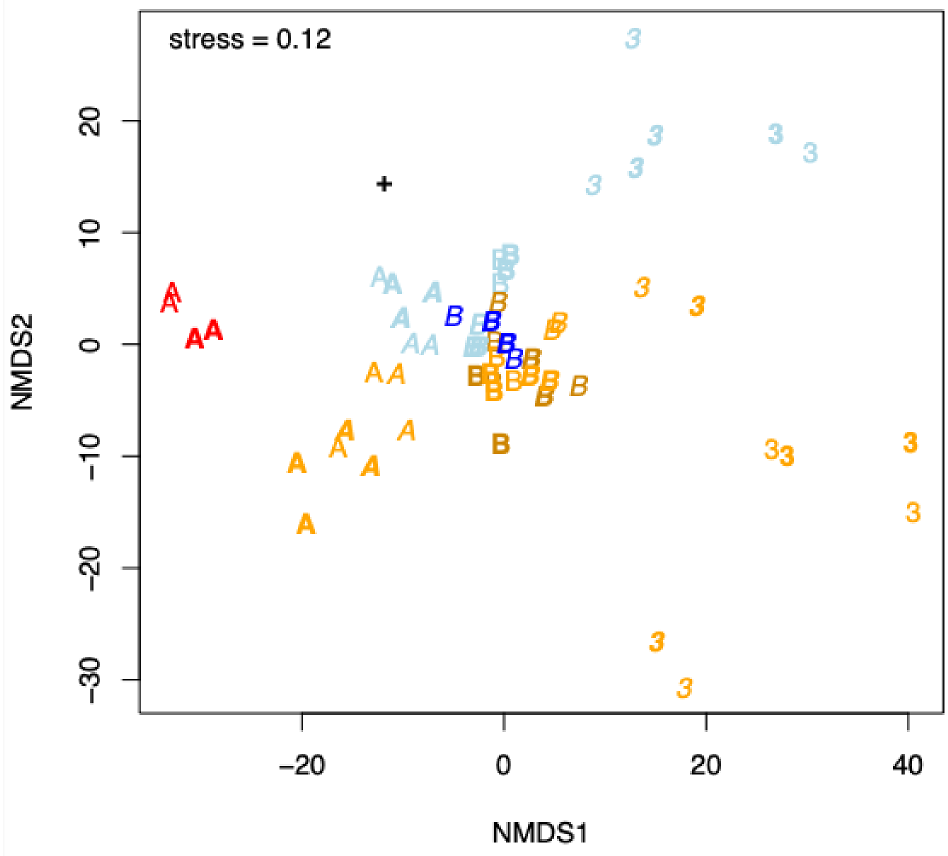


**Fig. 5.** Nonmetric Multidimensional Scaling (NMDS) representation of the 61x61 matrix of RF distances between all ML trees (60 IQtree runs + the M20 tree). Values <0.1 are typically taken to indicate "excellent," and <0.2 "good," 2-dimensional approximations of the original distances (Clarke 1993). Legend as in Figure 4. M20 tree = **+**. The trees using both AA+3Di data ("B") show a merger of both signals and cluster tightly, compared to the variability between the AA-only or (particularly) 3Di-only trees.

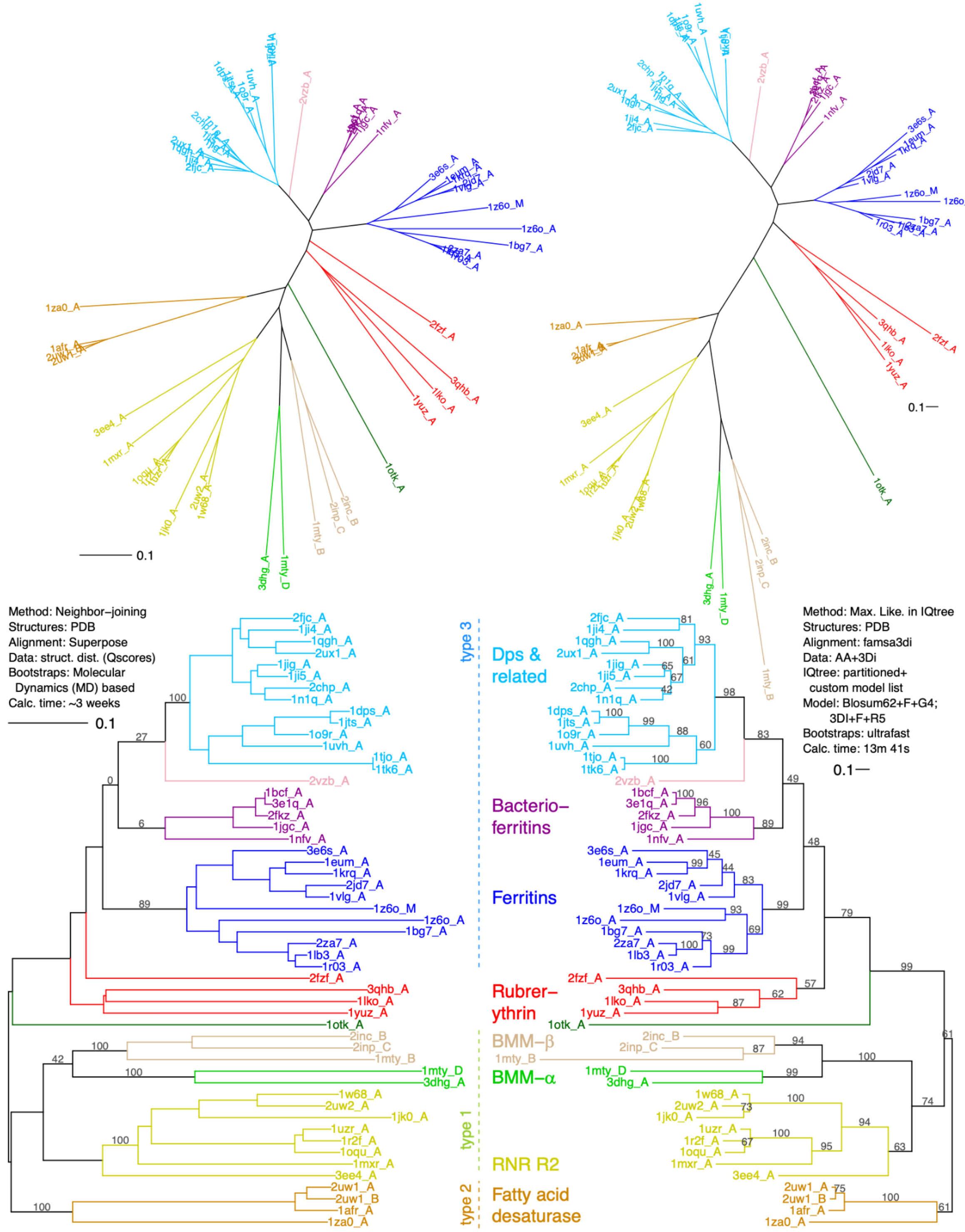


**Figure 6.** Comparison of trees using PDB structures. **Left:** M20 structural distances tree, with MD-based bootstraps where available in Malik et al. 2020. **Right:** the closest-match PDB-based IQtree analysis using AAs+3Dis from PDB structural data. This tree shares 20 deep and 19 shallow bipartitions with M20. **Top row:** unrooted trees. **Bottom:** arbitrary rooting for visual comparison. Branch length units for the M20 tree are $Q_{score}$-based distances; for the ML tree they are expected substitutions per site. The types refer to dimer geometries, likely slowly evolving structural characters (Lundin et al. 2012).

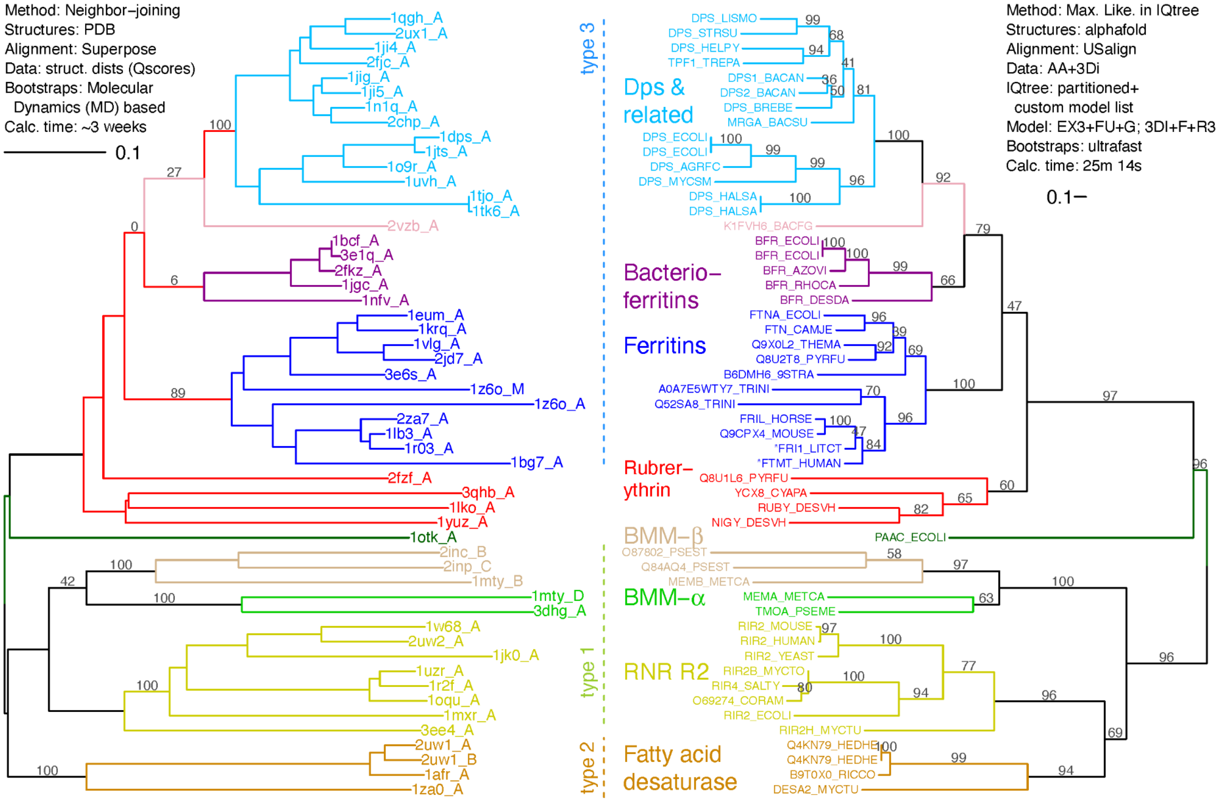


**Figure 7.** AlphaFold-based ML AA+3Di tree (right) compared to the M20 structural distances tree (left). This ML run was the closest overall topology match to the M20 tree, and shares 20 deep and 20 shallow bipartitions with M20. Tip labels are Uniprot IDs for which AlphaFold predictions were downloaded. Each Uniprot label on the right represents the closest AA match to the corresponding PDB structure on the left, with the exception of two tips marked with asterisks; here, FRI1_LITCT corresponds to 1bh7_A, and FTMT_HUMAN corresponds to 1r03_A. The types refer to dimer geometries, likely slowly evolving structural characters (Lundin et al. 2012).

The resulting topology from a "standard", sequence-only, structure-free alignment (Figure 8) shares 19 shallow bipartitions with M20, but only 12 deep ones. Severe long-branch attraction effects are evident; instead of 2fzf_A grouping with the rubrerythrins and 1z60_A with 1z60_M and other ferritins, they instead form a group of oddballs along with the long-branched 1otk_A. The BMM-β group breaks apart and renders fatty acid desaturases and ferritins paraphyletic.

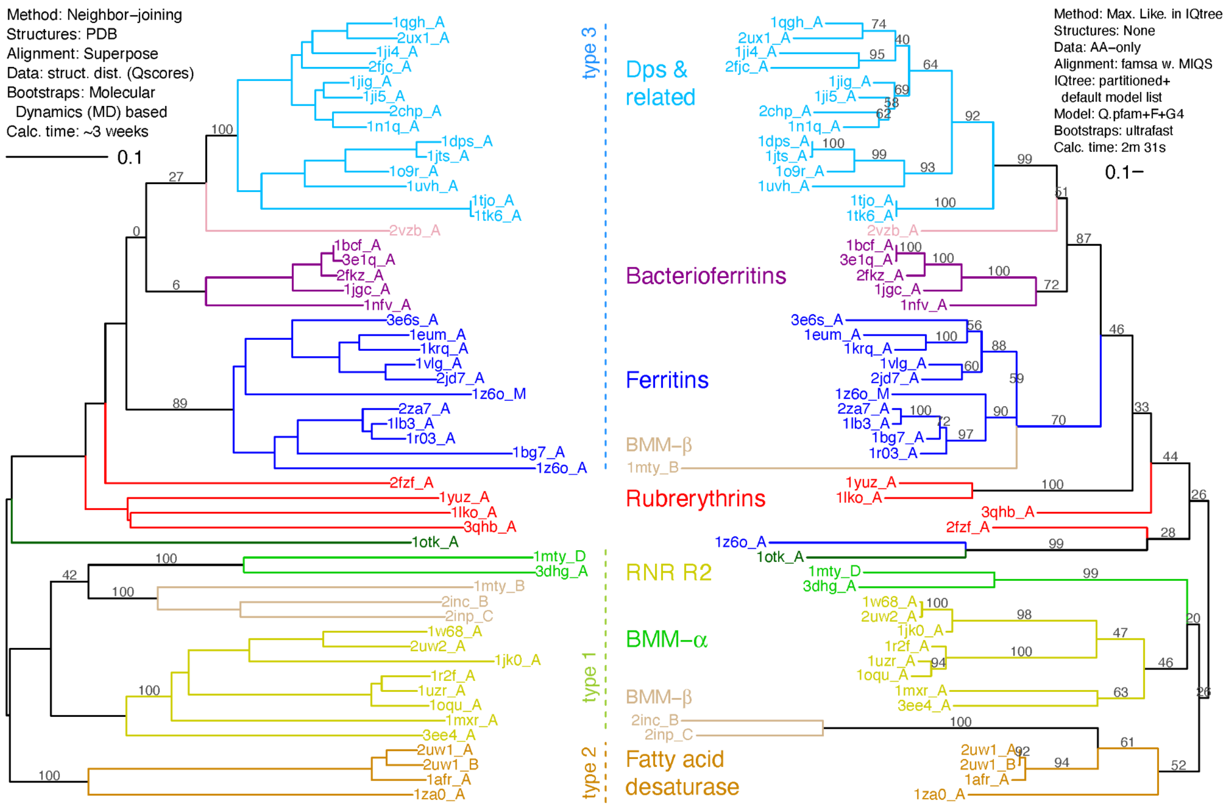


**Figure 8.** Structure, AA-only ML tree (right) compared to the M20 structural distances tree (left). ML tree is from the closest match "standard" AA-only analysis. The tree shares 12 deep and 19 shallow bipartitions with M20. The types refer to dimer geometries, likely slowly evolving structural characters (Lundin et al. 2012).

Goldman N, Thorne JL, Jones DT. Assessing the impact of secondary structure and solvent accessibility on protein evolution. *Genetics*. 149(1), 445-458. doi: 10.1093/genetics/149.1.445

Grant BJ, Rodrigues AP, ElSawy KM, McCammon JA, Caves LS. Bio3d: an R package for the comparative analysis of protein structures. *Bioinformatics*. 22(21), 2695-2696. doi: 10.1093/bioinformatics/btl461.

Grant BJ, Skjaerven L, Yao XQ. The Bio3D packages for structural bioinformatics. *Protein Sci*. 30(1), 20-30. doi: 10.1002/pro.3923

Guindon, Stéphane; Dufayard, Jean-François; Lefort, Vincent; Anisimova, Maria; Hordijk, Wim; Gascuel, Olivier (2010). New Algorithms and Methods to Estimate Maximum-Likelihood Phylogenies: Assessing the Performance of PhyML 3.0. *Systematic Biology*, 59(3), 307-321. doi: 10.1093/sysbio/syq010

Hasegawa H, Holm L. Advances and pitfalls of protein structural alignment. *Curr Opin Struct* Biol. 19(3), 341-348. doi: 10.1016/j.sbi.2009.04.003

Holm L, Sander C. Dali/FSSP classification of three-dimensional protein folds. *Nucleic Acids Research* 25(1), 231-234. doi: 10.1093/nar/25.1.231

Huelsenbeck, J. P., Bull, J. J., & Cunningham, C. W. (1996). Combining data in phylogenetic analysis. *Trends in Ecology & Evolution*, 11(4), 152-158. doi: 10.1016/0169-5347(96)10006-9

Huson, D. H., & Bryant, D. (2006). Application of Phylogenetic Networks in Evolutionary Studies, *Mol. Biol. Evol*., 23(2), 254-267.

Jeanmougin, F., Thompson, J. D., Gouy, M., Higgins, D. G., & Gibson, T. J. (1998). Multiple sequence alignment with Clustal X. *Trends in Biochemical Sciences*, *23*(10), 403-405.

Jumper J, Evans R, Pritzel A, Green T, Figurnov M, Ronneberger O, Tunyasuvunakool K, Bates R, Žídek A, Potapenko A, Bridgland A, Meyer C, Kohl SAA, Ballard AJ, Cowie A, Romera-Paredes B, Nikolov S, Jain R, Adler J, Back T, Petersen S, Reiman D, Clancy E, Zielinski M, Steinegger M, Pacholska M, Berghammer T, Bodenstein S, Silver D, Vinyals O, Senior AW, Kavukcuoglu K, Kohli P, Hassabis D. Highly accurate protein structure prediction with AlphaFold. *Nature*. 596(7873), 583-589. doi: 10.1038/s41586-021-03819-2

Kabsch W, Sander C. Dictionary of protein secondary structure: pattern recognition of hydrogen-bonded and geometrical features. *Biopolymers*. 22(12), 2577-2637. doi: 10.1002/bip.360221211

Katoh, Kazutaka, et al. "MAFFT: a novel method for rapid multiple sequence alignment based on fast Fourier transform." *Nucleic Acids Research* 30(14) (2002), 3059-3066.

Korber, Bette T., et al. "Covariation of mutations in the V3 loop of human immunodeficiency virus type 1 envelope protein: an information theoretic analysis." *Proceedings of the National Academy of Sciences* 90(15), 7176-7180.

Krissinel E, Henrick K. Secondary-structure matching (SSM), a new tool for fast protein structure alignment in three dimensions. *Acta Crystallogr D Biol Crystallogr.* 60(Pt 12 Pt 1), 2256-2268. doi: 10.1107/S0907444904026460

Lake JA, Henderson E, Oakes M, Clark MW. Eocytes: a new ribosome structure indicates a kingdom with a close relationship to eukaryotes. *Proceedings of the National Academy of Sciences* 81(12), 3786-3790. doi: 10.1073/pnas.81.12.3786.

Lai, Jhih-Siang 2020. *Protein structural phylogeny, a missing chapter in molecular evolutionary biology*. Ph.D. thesis, University of Queensland. doi: 10.14264/uql.2020.984

Lai, J-S, Rost, B, Kobe, B, Bodén, M. (2020). Evolutionary model of protein secondary structure capable of revealing new biological relationships. *Proteins*. 88, 1251-1259. https://doi-org/10.1002/prot.25898

Le, Si Quang; Lartillot, Nicolas; Gascuel, Olivier (2008). Phylogenetic mixture models for proteins. *Phil. Trans. R. Soc. B*. 363, 3965-3976. http://doi.org/10.1098/rstb.2008.0180

Le SQ, Gascuel O. Accounting for solvent accessibility and secondary structure in protein phylogenetics is clearly beneficial. *Syst Biol*. 59(3), 277-287. doi: 10.1093/sysbio/syq002

Lefort V, Desper R, Gascuel O. FastME 2.0: A Comprehensive, Accurate, and Fast Distance-Based Phylogeny Inference Program. *Mol Biol Evol*. 32(10), 2798-2800. doi: 10.1093/molbev/msv150

Lundin D, Poole AM, Sjöberg BM, Högbom M. Use of structural phylogenetic networks for classification of the ferritin-like superfamily. *J Biol Chem*. 287(24), 20565-20575. doi: 10.1074/jbc.M112.367458.

Malik, A. J., Poole, A. M., & Allison, J. R. (2020). Structural Phylogenetics with Confidence. *Molecular Biology and Evolution*, 37(9), 2711-2726. doi: 10.1093/molbev/msaa100

Matzke, Nicholas J.; Shih, Patrick M.; Kerfeld, Cheryl A. (2014). "Bayesian Analysis of Congruence of Core Genes in *Prochlorococcus* and *Synechococcus* and Implications on Horizontal Gene Transfer." *PLoS One*, 9(1), e85103. doi: 10.1371/journal.pone.0085103

Matzke, Nicholas (2015). The Evolution of Antievolution Policies After *Kitzmiller v. Dover*. *Science*, 351(6268), 10-12. doi: 10.1126/science.aad4057

Mariani V, Biasini M, Barbato A, Schwede T. lDDT: a local superposition-free score for comparing protein structures and models using distance difference tests. *Bioinformatics*. 29(21), 2722-2728. doi: 10.1093/bioinformatics/btt473

Meng EC, Goddard TD, Pettersen EF, Couch GS, Pearson ZJ, Morris JH, Ferrin TE. UCSF ChimeraX: Tools for structure building and analysis. *Protein Sci*. 32(11), e4792.

Minh, B. Q., Lanfear, R., Ly-Trong, N., Trifinopoulos, J., Schrempf, D., & Schmidt, H. A. (2022). IQ-TREE version 2.2.0: Tutorials and Manual. Phylogenomic software by maximum likelihood. March 25, 2022. http://www.iqtree.org/doc/iqtree-doc.pdf

Moi, David; Bernard, Charles; Steinegger, Martin; Nevers, Yannis; Langleib, Mauricio; Dessimoz, Christophe (2023). Structural phylogenetics unravels the evolutionary diversification of communication systems in gram-positive bacteria and their viruses. *bioRxiv* 2023.09.19.558401. doi: 10.1101/2023.09.19.558401

Nasrallah, C. A., Mathews, D. H., & Huelsenbeck, J. P. (2011). Quantifying the impact of dependent evolution among sites in phylogenetic inference. *Systematic Biology,* 60(1), 60-73.

Perrakis A, Sixma TK. AI revolutions in biology: The joys and perils of AlphaFold. *EMBO Rep.* 22(11), e54046. doi: 10.15252/embr.202154046

RCSB Protein Data Bank (RCSB.org): delivery of experimentally-determined PDB structures alongside one million computed structure models of proteins from artificial intelligence/machine learning (2023) *Nucleic Acids Research* 51: D488–D508 doi: 10.1093/nar/gkac1077

Ruff KM, Pappu RV. AlphaFold and Implications for Intrinsically Disordered Proteins. *J Mol Biol.* 433(20), 167208. doi: 10.1016/j.jmb.2021.167208

Salvador Capella-Gutierrez; Jose M. Silla-Martinez; Toni Gabaldon. (2009). trimAl: a tool for automated alignment trimming in large-scale phylogenetic analyses. *Bioinformatics*, 25: 1972-1973.

Rajapaksa, Sandun; Konagurthu, Arun S.; Lesk, Arthur M. (2023). Sequence and structure alignments in post-AlphaFold era, *Current Opinion in Structural Biology*, 79, 102539. doi: 10.1016/j.sbi.2023.102539.

Sansom RS, Gabbott SE, Purnell MA. Non-random decay of chordate characters causes bias in fossil interpretation. *Nature*. 463(7282), 797-800. doi: 10.1038/nature08745

Schliep KP. phangorn: phylogenetic analysis in R. *Bioinformatics*. 27(4) 592-593. doi: 10.1093/bioinformatics/btq706

Shih, Patrick M.; Matzke, Nicholas J. (2013). "Primary endosymbiosis events date to the later Proterozoic with cross-calibrated phylogenetic dating of duplicated ATPase proteins." *Proceedings of the National Academy of Sciences*, 110(30), 12355-12360.

Sillitoe I, Dawson N, Lewis TE, Das S, Lees JG, Ashford P, Tolulope A, Scholes HM, Senatorov I, Bujan A, Ceballos Rodriguez-Conde F, Dowling B, Thornton J, Orengo CA. CATH: expanding the horizons of structure-based functional annotations for genome sequences. *Nucleic Acids Res*. 47(D1), D280-D284. doi: 10.1093/nar/gky1097

Skolnick J, Gao M, Zhou H, Singh S. AlphaFold 2: Why It Works and Its Implications for Understanding the Relationships of Protein Sequence, Structure, and Function. *J Chem Inf Model*. 61(10), 4827-4831. doi: 10.1021/acs.jcim.1c01114

Tammi, Martti T. (2021). Construction of substitution matrices. Bioinformaticshome.com. Accessed 2021-06-01. https://bioinformaticshome.com/bioinformatics_tutorials/sequence_alignment/substitution_matrices.html

Tatur, J.; Hagen, W. R.; Matias, P. M. (2007). Crystal structure of the ferritin from the hyperthermophilic archaeal anaerobe *Pyrococcus furiosus*. *JBIC Journal of Biological Inorganic Chemistry*, *12*, 615-630. doi: 10.1007/s00775-007-0212-3

Trinquier, J.; Petti, S.; Feng, S.; Söding, J.; Steinegger, M.; Ovchinnikov, S. (2022). SWAMPNN: End-to-end protein structures alignment. *Machine Learning for Structural Biology Workshop*, NeurIPS, 2022. https://www.mlsb.io/papers_2022/SWAMPNN_End_to_end_protein_structures_alignment.pdf

Thorne JL, Goldman N, Jones DT. Combining protein evolution and secondary structure. *Mol Biol Evol*. 13(5), 666-673. doi: 10.1093/oxfordjournals.molbev.a025627.

van Kempen M, Kim SS, Tumescheit C, Mirdita M, Lee J, Gilchrist CLM, Söding J, Steinegger M. Fast and accurate protein structure search with Foldseek. *Nat Biotechnol*. doi: 10.1038/s41587-023-01773-0

Varadi M, Anyango S, Deshpande M, Nair S, Natassia C, Yordanova G, Yuan D, Stroe O, Wood G, Laydon A, Žídek A, Green T, Tunyasuvunakool K, Petersen S, Jumper J, Clancy E, Green R, Vora A, Lutfi M, Figurnov M, Cowie A, Hobbs N, Kohli P, Kleywegt G, Birney E, Hassabis D, Velankar S. AlphaFold Protein Structure Database: massively expanding the structural coverage of protein-sequence space with high-accuracy models. *Nucleic Acids Research*. 50(D1), D439-D444. doi: 10.1093/nar/gkab1061

Varadi M, Bertoni D, Magana P, Paramval U, Pidruchna I, Radhakrishnan M, Tsenkov M, Nair S, Mirdita M, Yeo J, Kovalevskiy O, Tunyasuvunakool K, Laydon A, Žídek A, Tomlinson H, Hariharan D, Abrahamson J, Green T, Jumper J, Birney E, Steinegger M, Hassabis D, Velankar S. (2023). AlphaFold Protein Structure Database in 2024: providing structure coverage for over 214 million protein sequences. *Nucleic Acids Research*, gkad1011. doi: 10.1093/nar/gkad1011

Waterhouse AM, Procter JB, Martin DMA, Clamp M, Barton GJ (2009). Jalview Version 2 - A multiple sequence alignment editor and analysis workbench. *Bioinformatics* 25 1189-1191. doi:10.1093/bioinformatics/btp03

Zhang C, Shine M, Pyle AM, Zhang Y. US-align: universal structure alignments of proteins, nucleic acids, and macromolecular complexes. *Nat Methods*. 19(9), 1109-1115. doi: 10.1038/s41592-022-01585-1.

Zhang Y, Skolnick J. 2005. TM-align: a protein structure alignment algorithm based on the TM-score. *Nucleic Acids Research*, 33(7), 2302-2309.

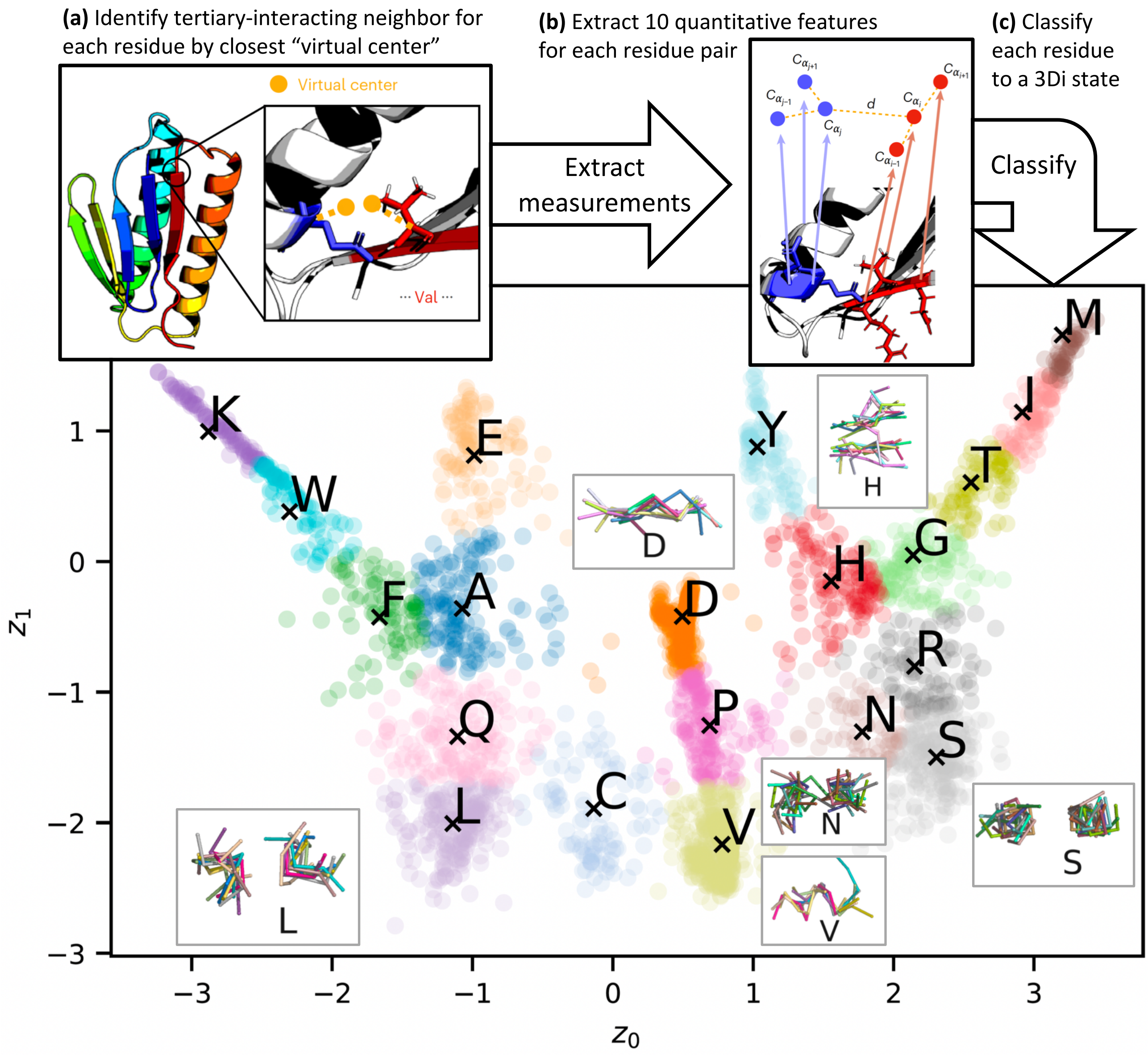

(a) Identify tertiary-interacting neighbor for each residue by closest "virtual center"
Virtual center
··· Val ···
(b) Extract 10 quantitative features for each residue pair
Extract measurements
d
(c) Classify each residue to a 3Di state
Classify
K
W
E
F
A
Q
L
C
D
P
V
Y
H
M
I
T
G
R
N
S
$z_1$
$z_0$

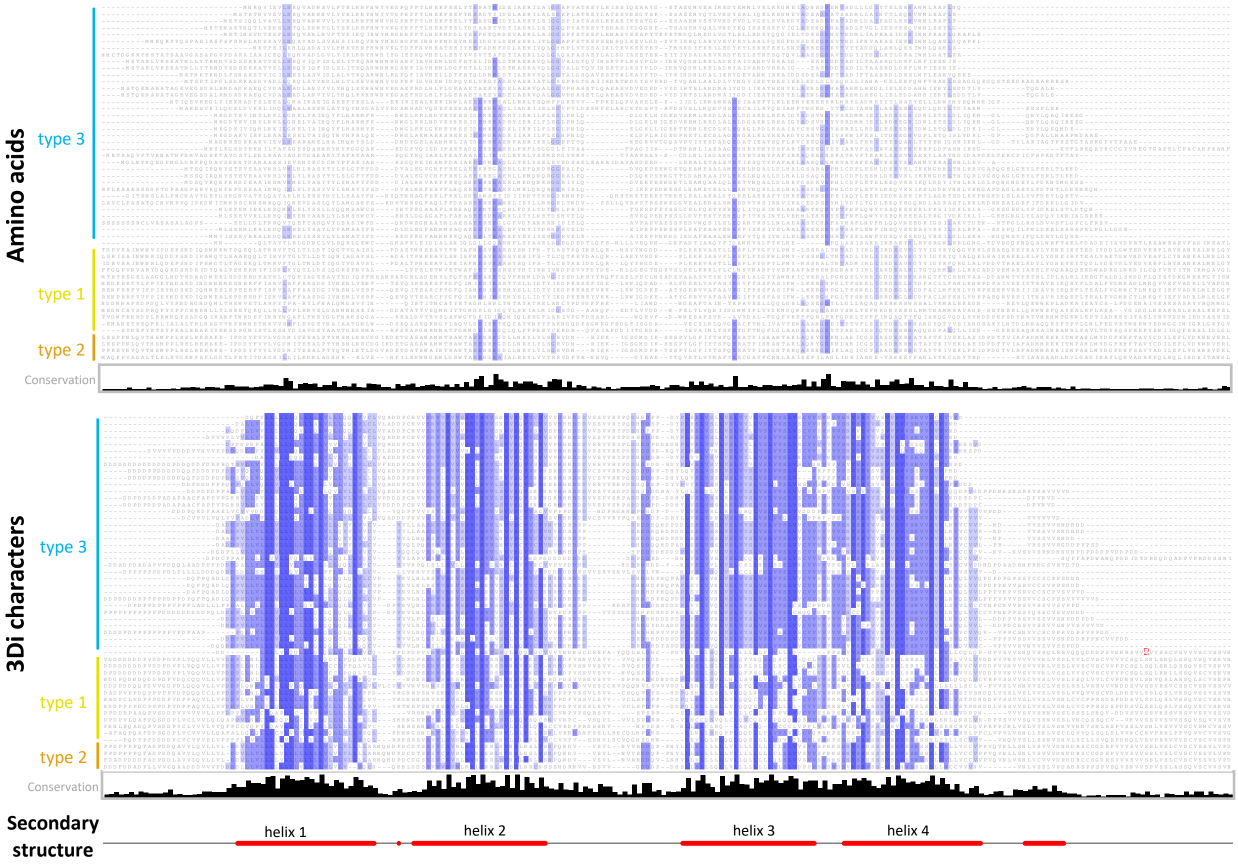

Amino acids
type 3
type 1
type 2
Conservation
3Di characters
type 3
type 1
type 2
Conservation
Secondary structure
helix 1
helix 2
helix 3
helix 4

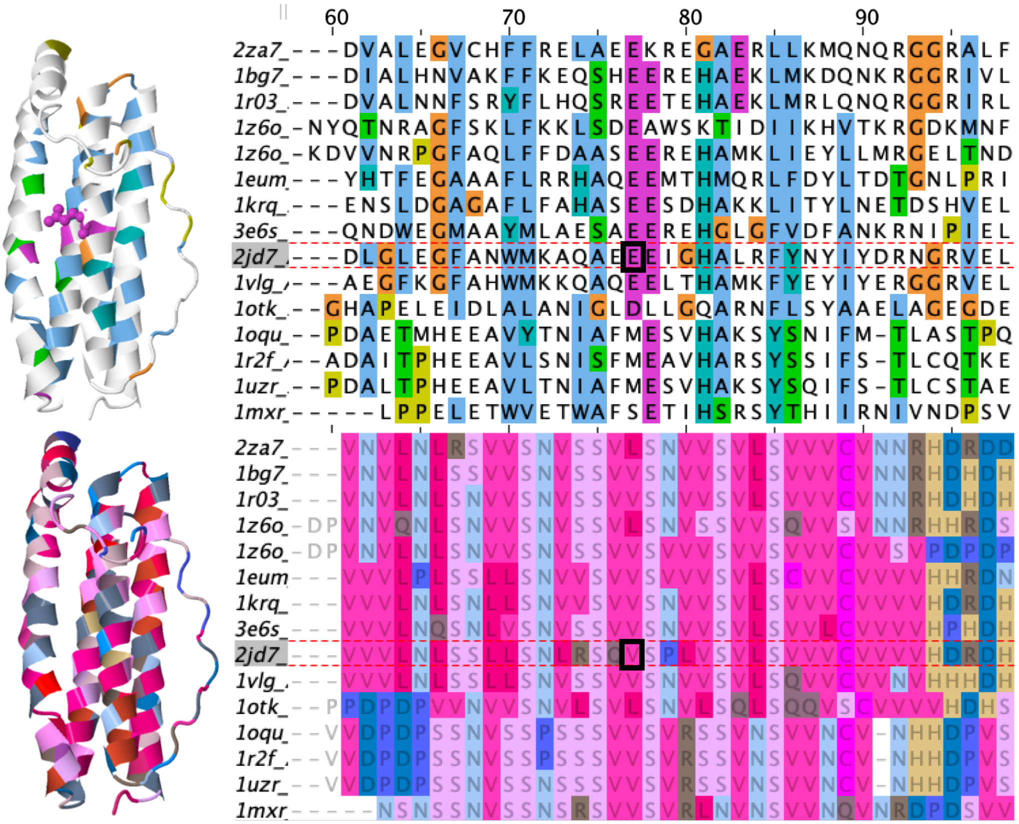

60 70 80 90
2za7_ ---DVALEGVCHFFRELAEEKREGAERLLKMQNQRGGRALF
1bg7_ ---DIALHNVAKFFKEQSHEEREHAEKLMKDQNKRGGRIVL
1r03_ ---DVALNNFSRYFLHQSREETEHAEKLMRLQNQRGGRIRL
1z6o_ -NYQTNRAGFSKLFKKLSDEAWSKTIDIIKHVTKRGDKMNF
1z6o_ -KDVVNRPGFAQLFFDAASEEREHAMKLIEYLLMRGELTND
1eum ---YHTFEGAAAFLRRHAQEEMTHMQRLFDYLTDTGNLPRI
1krq_ ---ENSLDGAGAFLFAHASEESDHAKKLITYLNETDSHVEL
3e6s_ ---QNDWEGMAAYMLAESAEEREHGLGFVDFANKRNIPIEL
2jd7_ ---DLGLEGFANWMKAQAEEEIGHALRFYNYIYDRNGRVEL
1vlg_ ---AEGFKGFAHWMKKQAQEELTHAMKFYEYIYERGGRVEL
1otk_ --GHAPELEIDLALANIGLDLLGQARNFLSYAAELAGEGDE
1oqu_ --PDAETMHEEAVYTNIAFMESVHAKSYSNIFM-TLASTPQ
1r2f_ --ADAITPHEEAVLSNISFMEAVHARSYSSIFS-TLCQTKE
1uzr_ --PDALTPHEEAVLTNIAFMESVHAKSYSQIFS-TLCSTAE
1mxr_ -----LPPELETWVETWAFSETIHSRSYTHIIRNIVNDPSV
2za7_
1bg7_
1r03_
1z6o_
1z6o_
1eum
1krq_
3e6s_
2jd7_
1vlg_
1otk_
1oqu_
1r2f_
1uzr_
1mxr_

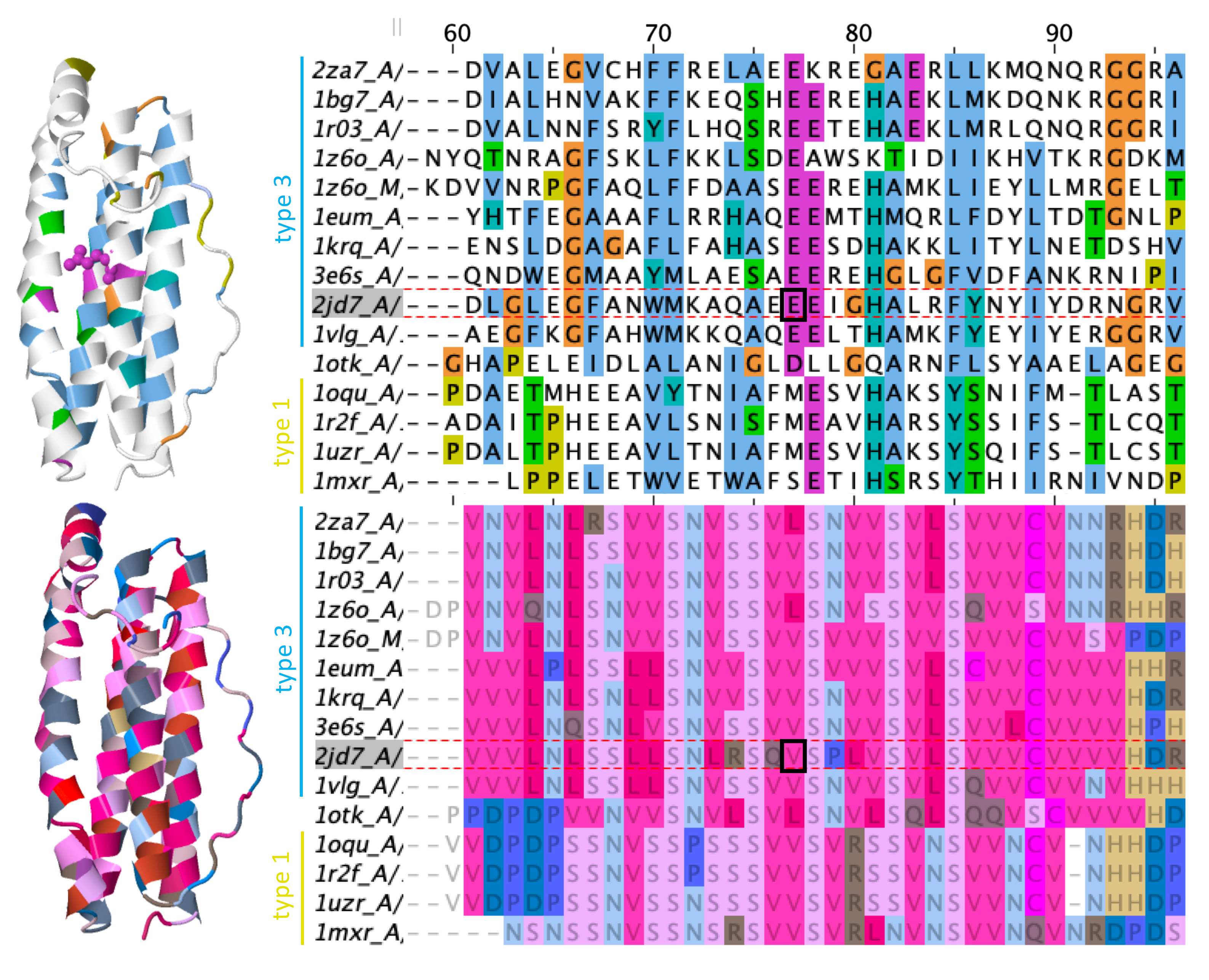
60
70
80
90
type 3
type 1
2za7_A/
1bg7_A/
1r03_A/
1z6o_A/
1z6o_M/
1eum_A
1krq_A/
3e6s_A/
2jd7_A/
1vlg_A/
1otk_A/
1oqu_A/
1r2f_A/
1uzr_A/
1mxr_A/

**Table 1.** Best-fit substitution models for IQtree runs, according to log-likelihood (lnL) and BIC. IQtree inferences were conducted using the BIC-best model. Runs are sorted by deep & shallow similarity to the M20 tree (first line). Results are shown only for IQtree runs using the custom model list (with partitioning for AA+3Di dataset) which included the 3Di substitution matrix. *k* represents the number of free parameters in the substitution model; all runs also included 103 free branch-length parameters in the BIC calculation. Note: lnL & BIC values can only be compared within lines, not between them, as they are only comparable for identical alignments.

| **Structure source** | **Alignment data** | **Alignment method** | **Phylogeny data** | **# bipartitions matched** | | | **Best log-likelihood models:** | | | **Best BIC models:** | | | |
|---|---|---|---|---|---|---|---|---|---|---|---|---|---|
| | | | | **all** | **deep** | **shal-low** | **lnL** | ***k*** | **Model** | **BIC** | **lnL** | ***k*** | **Model** |
| PDB | structures | Superpose | *Q* score | 51 | 22 | 29 | | | | | | | |
| alphafold | struct.+AAs | USalign | AA+3di, trim | 40 | 20 | 20 | -24477.7 | 27 | EX3+FU+G; 3DI+F+R3 | 49728.3 | -24477.7 | 27 | EX3+FU+G; 3DI+F+R3 |
| PDB | 3di | famsa3di | AA+3di | 39 | 20 | 19 | -39984.6 | 48 | Blosum62+F+G4; 3DI+F+R5 | 81081.6 | -39984.6 | 48 | Blosum62+F+G4; 3DI+F+R5 |
| alphafold | struct.+AAs | USalign | AA+3di | 39 | 20 | 19 | -43208.6 | 47 | Blosum62+F+R3; 3DI+F+R3 | 87609.2 | -43208.6 | 47 | Blosum62+F+R3; 3DI+F+R3 |
| PDB | 3di | famsa3di | AA+3di, trim | 37 | 20 | 17 | -29156.4 | 41 | Blosum62+F+G4; 3DI+F+G4 | 59179.9 | -29156.4 | 41 | Blosum62+F+G4; 3DI+F+G4 |
| PDB | struct.+AAs | USalign | AA | 37 | 19 | 18 | -15160.7 | 25 | Blosum62+F+R4 | 30976.0 | -15208.7 | 4 | Blosum62+R3 |
| alphafold | 3di | famsa3di | AA+3di | 36 | 19 | 17 | -42057.8 | 44 | Blosum62+F+G4; 3DI+F+R3 | 85195.4 | -42057.8 | 44 | Blosum62+F+G4; 3DI+F+R3 |
| PDB | struct.+AAs | USalign | AA+3di | 34 | 19 | 15 | -40816.9 | 47 | Blosum62+F+R3; 3DI+F+R3 | 82772.5 | -40816.9 | 47 | Blosum62+F+R3; 3DI+F+R3 |
| alphafold | struct.+AAs | USalign | AA, trim | 38 | 18 | 20 | -16270.9 | 25 | Blosum62+F+R4 | 33187.1 | -16315.2 | 3 | EX3+FU+G |
| PDB | 3di | famsa3di | AA | 36 | 18 | 18 | -24783.6 | 25 | Blosum62+F+R4 | 50404.9 | -24792.0 | 20 | Blosum62+F+G4 |
| alphafold | struct.+AAs | USalign | AA | 36 | 18 | 18 | -27095.9 | 25 | Blosum62+F+R4 | 55106.9 | -27107.4 | 20 | Blosum62+F+G4 |
| PDB | struct.+AAs | USalign | AA | 36 | 18 | 18 | -24877.6 | 25 | Blosum62+F+R4 | 50631.3 | -24881.1 | 23 | Blosum62+F+R3 |
| PDB | struct.+AAs | USalign | AA+3di, trim | 33 | 18 | 15 | -23679.1 | 25 | Blosum62+R3; 3DI+F+G4 | 48115.0 | -23679.1 | 25 | Blosum62+R3; 3DI+F+G4 |
| alphafold | 3di | famsa3di | AA+3di, trim | 36 | 17 | 19 | -30459.0 | 27 | EX3+FU+G; 3DI+F+R3 | 61715.6 | -30459.0 | 27 | EX3+FU+G; 3DI+F+R3 |
| alphafold | 3di | famsa3di | 3di | 32 | 17 | 15 | -14950.2 | 25 | 3DI+F+R4 | 30745.2 | -14953.6 | 23 | 3DI+F+R3 |
| alphafold | struct.+AAs | USalign | 3di | 31 | 16 | 15 | -15985.7 | 25 | 3DI+F+R4 | 32886.3 | -15986.2 | 23 | 3DI+F+R3 |
| alphafold | 3di | famsa3di | AA | 32 | 14 | 18 | -26932.0 | 27 | Blosum62+F+R5 | 54691.6 | -26993.3 | 3 | EX3+FU+G |
| alphafold | 3di | famsa3di | 3di, trim | 31 | 14 | 17 | -10259.3 | 27 | 3DI+F+R5 | 21218.1 | -10260.7 | 25 | 3DI+F+R4 |
| PDB | 3di | famsa3di | AA, trim | 30 | 14 | 16 | -18633.2 | 25 | Blosum62+F+R4 | 37938.1 | -18641.4 | 20 | Blosum62+F+G4 |
| PDB | struct.+AAs | USalign | 3di, trim | 23 | 14 | 9 | -8173.4 | 25 | 3DI+F+R4 | 17006.9 | -8174.6 | 23 | 3DI+F+R3 |
| PDB | struct.+AAs | USalign | 3di | 22 | 14 | 8 | -15612.5 | 23 | 3DI+F+R3 | 32094.1 | -15612.5 | 23 | 3DI+F+R3 |
| alphafold | struct.+AAs | USalign | 3di, trim | 28 | 13 | 15 | -8037.0 | 25 | 3DI+F+R4 | 16740.9 | -8039.5 | 23 | 3DI+F+R3 |
| PDB | 3di | famsa3di | 3di, trim | 25 | 13 | 12 | -10215.4 | 25 | 3DI+F+R4 | 21107.5 | -10218.1 | 23 | 3DI+F+R3 |
| none | AAs | famsa | AA, trim | 30 | 12 | 18 | -24136.9 | 23 | LG+F+R3 | 49084.3 | -24139.0 | 20 | LG+F+G4 |
| alphafold | 3di | famsa3di | AA, trim | 29 | 12 | 17 | -20019.9 | 25 | LG+F+R4 | 40665.8 | -20044.4 | 3 | EX3+FU+G |
| PDB | 3di | famsa3di | 3di | 24 | 12 | 12 | -14851.9 | 25 | 3DI+F+R4 | 30553.0 | -14856.1 | 23 | 3DI+F+R3 |
| none | AAs | famsa | AA | 29 | 10 | 19 | -18766.9 | 25 | LG+F+R4 | 38190.2 | -18813.5 | 1 | LG+G4 |

**Table 2.** Illustrative models compared on AA and 3Di datasets. Alignment source: USalign on alphafold structures+AAs, trimmed sites with <35% data.

| Data | Model | lnL | ΔlnL | *k* | BIC |
|---|---|---|---|---|---|
| amino acids (*n* =191 sites) | EX3+FU+G | -16315.2 | 35.9 | 106 | 33187.1 |
| | Blosum62+F+G4 | -16279.2 | 0.0 | 123 | 33204.5 |
| | LG+F+G4 | -16289.1 | 9.8 | 123 | 33224.2 |
| | EX3+F+G | -16299.1 | 19.8 | 125 | 33254.6 |
| | Poisson+F+G4 | -16888.9 | 609.7 | 123 | 34423.9 |
| | 3DI+F+G4 | -17950.8 | 1671.5 | 123 | 36547.6 |
| 3Di characters (*n* =191 sites) | 3DI+F+G4 | -8054.9 | 0.0 | 123 | 16755.9 |
| | Poisson+F+G4 | -8329.4 | 274.5 | 123 | 17304.9 |
| | Blosum62+F+G4 | -8485.0 | 430.0 | 123 | 17616.0 |
| | EX3+F+G | -8620.9 | 566.0 | 125 | 17898.3 |
| | LG+F+G4 | -8734.1 | 679.1 | 123 | 18114.1 |
| | EX3+FU+G | -10415.9 | 2361.0 | 106 | 21388.6 |

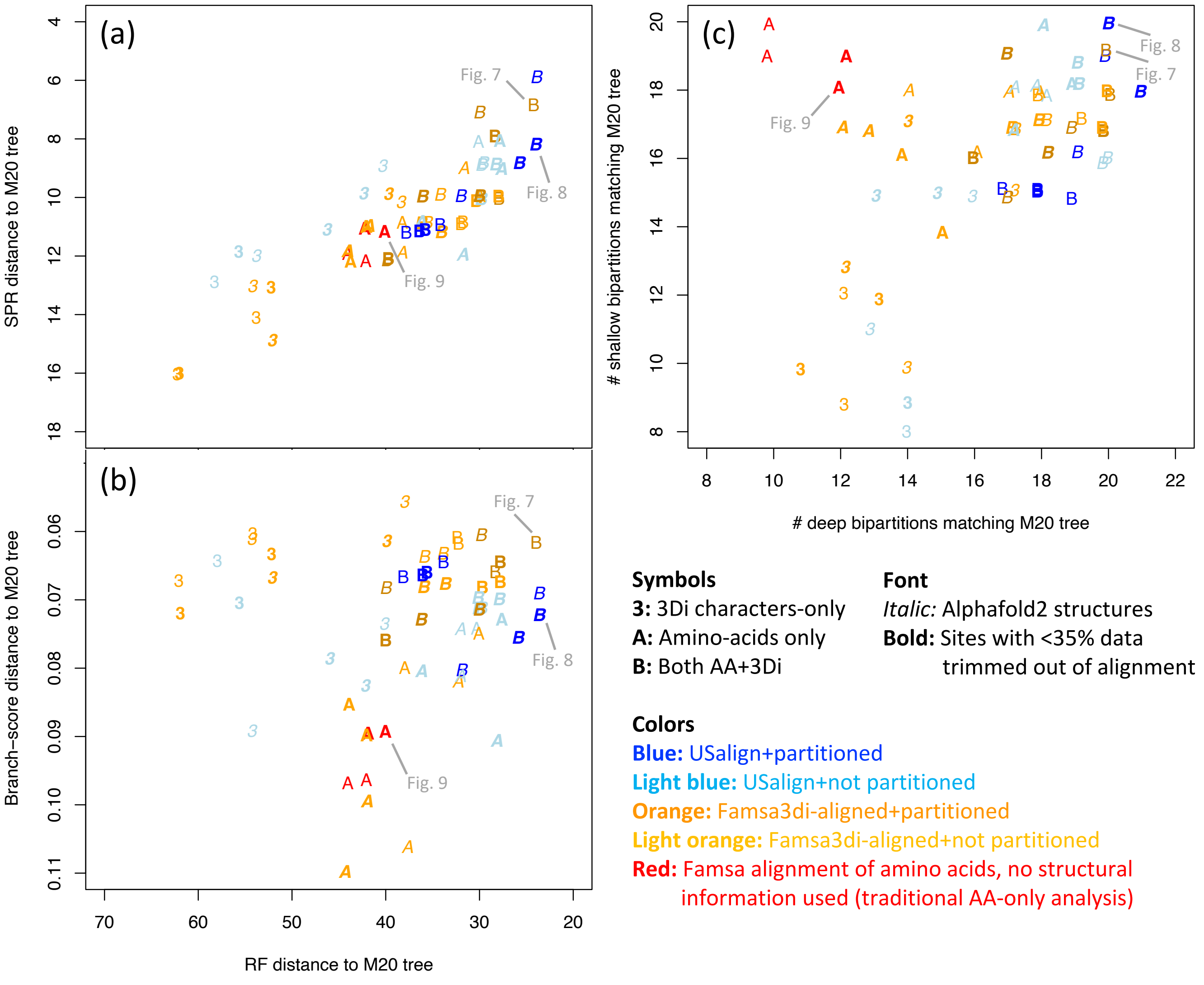
(a)
(b)
(c)
SPR distance to M20 tree
Branch−score distance to M20 tree
RF distance to M20 tree
### shallow bipartitions matching M20 tree
### deep bipartitions matching M20 tree
Fig. 7
Fig. 8
Fig. 9
Symbols
3: 3Di characters-only
A: Amino-acids only
B: Both AA+3Di
Font
Italic: Alphafold2 structures
Bold: Sites with <35% data trimmed out of alignment
Colors
Blue: USalign+partitioned
Light blue: USalign+not partitioned
Orange: Famsa3di-aligned+partitioned
Light orange: Famsa3di-aligned+not partitioned
Red: Famsa alignment of amino acids, no structural information used (traditional AA-only analysis)

**Nonmetric Multi−dimensional Scaling (NMDS)**
**RF distances between 61 trees**

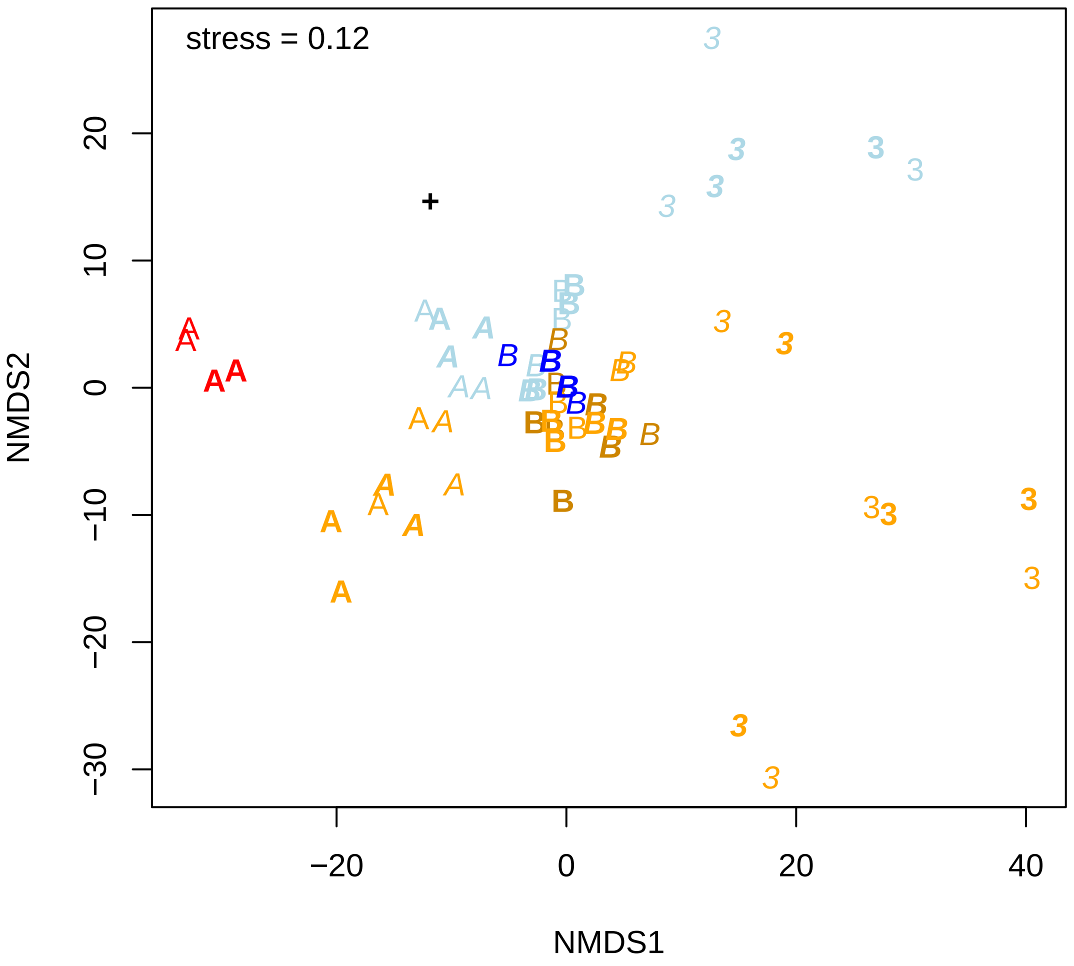

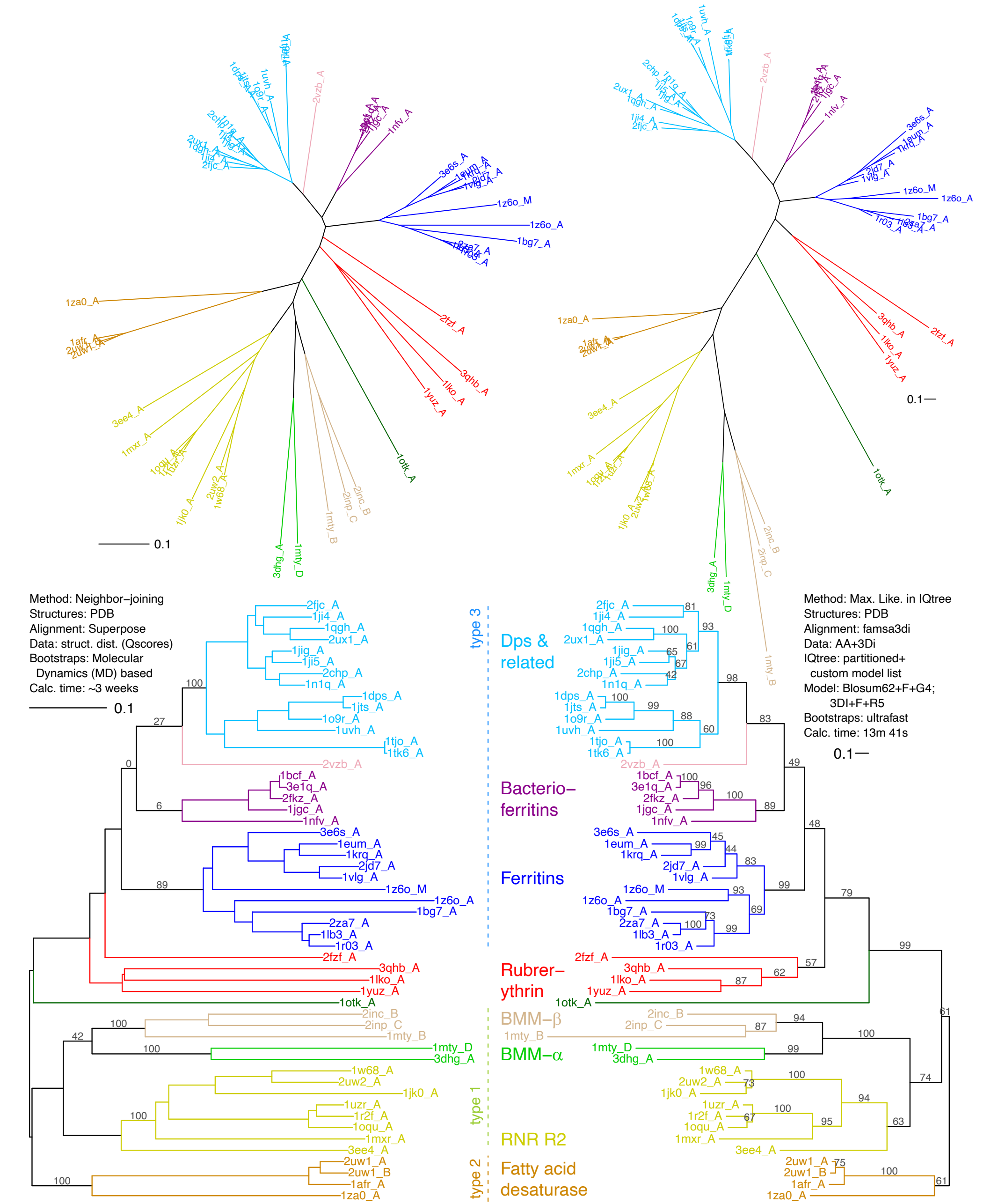

0.1
Method: Neighbor−joining
Structures: PDB
Alignment: Superpose
Data: struct. dist. (Qscores)
Bootstraps: Molecular Dynamics (MD) based
Calc. time: ~3 weeks
0.1
Method: Max. Like. in IQtree
Structures: PDB
Alignment: famsa3di
Data: AA+3Di
IQtree: partitioned+ custom model list
Model: Blosum62+F+G4; 3DI+F+R5
Bootstraps: ultrafast
Calc. time: 13m 41s
0.1
type 3
Dps & related
Bacterio−ferritins
Ferritins
Rubrer−ythrin
BMM−β
BMM−α
type 1
RNR R2
type 2
Fatty acid desaturase
2fjc_A
1ji4_A
1qgh_A
2ux1_A
1jig_A
1ji5_A
2chp_A
1n1q_A
1dps_A
1jts_A
1o9r_A
1uvh_A
1tjo_A
1tk6_A
2vzb_A
1bcf_A
3e1q_A
2fkz_A
1jgc_A
1nfv_A
3e6s_A
1eum_A
1krq_A
2jd7_A
1vlg_A
1z6o_M
1z6o_A
1bg7_A
2za7_A
1lb3_A
1r03_A
2fzf_A
3qhb_A
1lko_A
1yuz_A
1otk_A
2inc_B
2inp_C
1mty_B
1mty_D
3dhg_A
1w68_A
2uw2_A
1jk0_A
1uzr_A
1r2f_A
1oqu_A
1mxr_A
3ee4_A
2uw1_A
2uw1_B
1afr_A
1za0_A

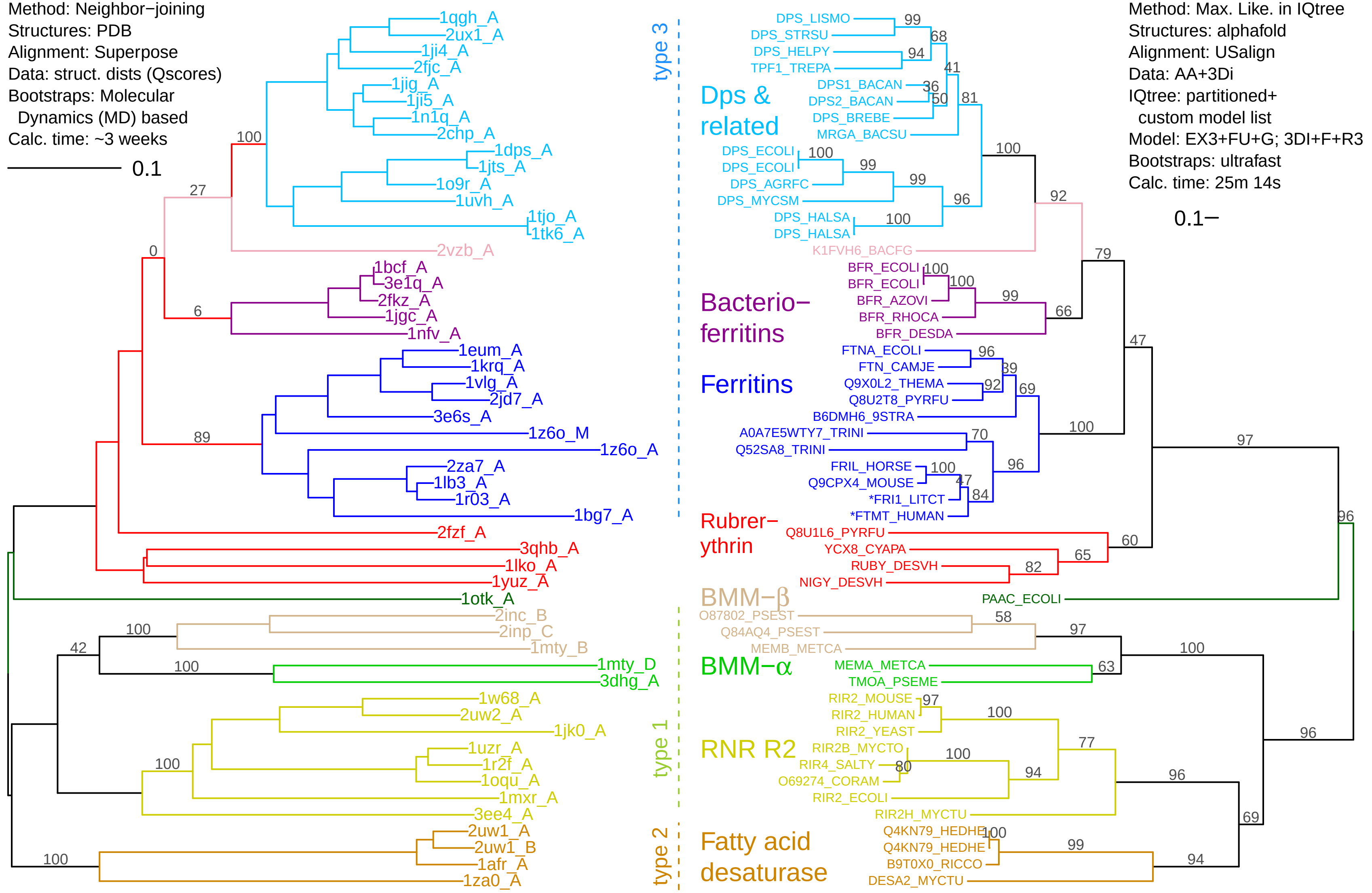
Method: Neighbor−joining
Structures: PDB
Alignment: Superpose
Data: struct. dists (Qscores)
Bootstraps: Molecular Dynamics (MD) based
Calc. time: ~3 weeks
0.1
1qgh_A
2ux1_A
1ji4_A
2fjc_A
1jig_A
1ji5_A
1n1q_A
2chp_A
1dps_A
1jts_A
1o9r_A
1uvh_A
1tjo_A
1tk6_A
2vzb_A
1bcf_A
3e1q_A
2fkz_A
1jgc_A
1nfv_A
1eum_A
1krq_A
1vlg_A
2jd7_A
3e6s_A
1z6o_M
1z6o_A
2za7_A
1lb3_A
1r03_A
1bg7_A
2fzf_A
3qhb_A
1lko_A
1yuz_A
1otk_A
2inc_B
2inp_C
1mty_B
1mty_D
3dhg_A
1w68_A
2uw2_A
1jk0_A
1uzr_A
1r2f_A
1oqu_A
1mxr_A
3ee4_A
2uw1_A
2uw1_B
1afr_A
1za0_A
100
27
0
6
89
100
42
100
100
100
type 3
type 1
type 2
Dps & related
Bacterio−ferritins
Ferritins
Rubrer−ythrin
BMM−β
BMM−α
RNR R2
Fatty acid desaturase
Method: Max. Like. in IQtree
Structures: alphafold
Alignment: USalign
Data: AA+3Di
IQtree: partitioned+ custom model list
Model: EX3+FU+G; 3DI+F+R3
Bootstraps: ultrafast
Calc. time: 25m 14s
0.1
DPS_LISMO
DPS_STRSU
DPS_HELPY
TPF1_TREPA
DPS1_BACAN
DPS2_BACAN
DPS_BREBE
MRGA_BACSU
DPS_ECOLI
DPS_ECOLI
DPS_AGRFC
DPS_MYCSM
DPS_HALSA
DPS_HALSA
K1FVH6_BACFG
BFR_ECOLI
BFR_ECOLI
BFR_AZOVI
BFR_RHOCA
BFR_DESDA
FTNA_ECOLI
FTN_CAMJE
Q9X0L2_THEMA
Q8U2T8_PYRFU
B6DMH6_9STRA
A0A7E5WTY7_TRINI
Q52SA8_TRINI
FRIL_HORSE
Q9CPX4_MOUSE
*FRI1_LITCT
*FTMT_HUMAN
Q8U1L6_PYRFU
YCX8_CYAPA
RUBY_DESVH
NIGY_DESVH
PAAC_ECOLI
O87802_PSEST
Q84AQ4_PSEST
MEMB_METCA
MEMA_METCA
TMOA_PSEME
RIR2_MOUSE
RIR2_HUMAN
RIR2_YEAST
RIR2B_MYCTO
RIR4_SALTY
O69274_CORAM
RIR2_ECOLI
RIR2H_MYCTU
Q4KN79_HEDHE
Q4KN79_HEDHE
B9T0X0_RICCO
DESA2_MYCTU
99
68
94
41
36
50
81
100
99
99
96
100
100
92
79
100
100
99
66
96
39
92
69
47
100
70
96
100
47
84
60
65
82
97
96
58
97
100
63
97
100
77
100
80
94
96
69
96
100
99
94

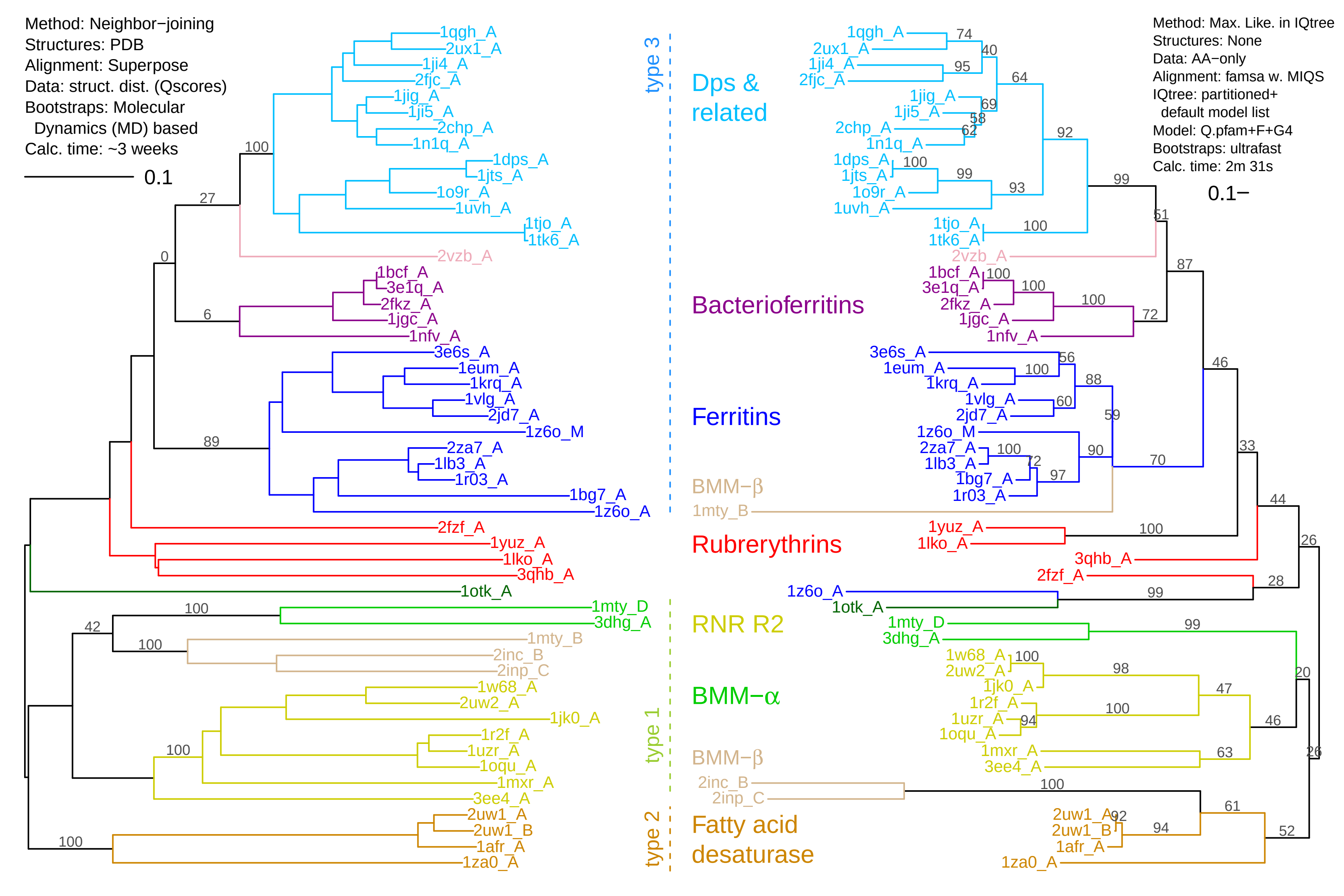

Method: Neighbor−joining
Structures: PDB
Alignment: Superpose
Data: struct. dist. (Qscores)
Bootstraps: Molecular Dynamics (MD) based
Calc. time: ~3 weeks
0.1
Method: Max. Like. in IQtree
Structures: None
Data: AA−only
Alignment: famsa w. MIQS
IQtree: partitioned+ default model list
Model: Q.pfam+F+G4
Bootstraps: ultrafast
Calc. time: 2m 31s
0.1
type 3
type 1
type 2
Dps & related
Bacterioferritins
Ferritins
BMM−β
Rubrerythrins
RNR R2
BMM−α
BMM−β
Fatty acid desaturase
1qgh_A
2ux1_A
1ji4_A
2fjc_A
1jig_A
1ji5_A
2chp_A
1n1q_A
1dps_A
1jts_A
1o9r_A
1uvh_A
1tjo_A
1tk6_A
2vzb_A
1bcf_A
3e1q_A
2fkz_A
1jgc_A
1nfv_A
3e6s_A
1eum_A
1krq_A
1vlg_A
2jd7_A
1z6o_M
2za7_A
1lb3_A
1r03_A
1bg7_A
1z6o_A
2fzf_A
1yuz_A
1lko_A
3qhb_A
1otk_A
1mty_D
3dhg_A
1mty_B
2inc_B
2inp_C
1w68_A
2uw2_A
1jk0_A
1r2f_A
1uzr_A
1oqu_A
1mxr_A
3ee4_A
2uw1_A
2uw1_B
1afr_A
1za0_A
