## Supplementary material for "Tertiary-interaction characters enable fast, model-based structural phylogenetics beyond the twilight zone": README.md

# 3diphy
Maximum likelihood structural phylogenetics by including Foldseek 3Di characters. Supporting Information for Puente-Lelievre et al. 2023.

The citation is:

Puente-Lelievre, Caroline; Malik, Ashar J.; Douglas, Jordan;  Ascher, David; Baker, Matthew; Allison, Jane; Poole, Anthony; Lundin, Daniel; Fullmer, Matthew; Bouckert, Remco; Steinegger, Martin; Matzke, Nicholas J. (2023). Tertiary-interaction characters enable fast, model-based structural phylogenetics beyond the twilight zone. *bioRxiv*, 571181. December 12, 2023. doi: https://www.biorxiv.org/content/10.1101/571181

For a README file / guide to the Supporting Information directory, please see: _00_README_SuppInf_files_v2.txt.
