## Supplementary material for "Tertiary-interaction characters enable fast, model-based structural phylogenetics beyond the twilight zone": parsing_iqtree_file_v1.R

```
get_numparams_from_AIC <- function(AIC, lnL)
	{
	'
	AIC = 92967.930
	lnL = -46353.965
	k = get_numparams_from_AIC(AIC, lnL)
	k

	iqtree_fn = "/GitHub/str2phy/ex/ferritins_M20af/03_USaln_v1/iqtree_3disTrim35/53fer.fasta.iqtree"
	iqtree_df = read_iqtree_model_scores(iqtree_fn, calc_extra=FALSE)
	k = get_numparams_from_AIC(AIC=iqtree_df$AIC, lnL=iqtree_df$LogL)
	k
	n = get_ndata_from_AICc(AICc=iqtree_df$AICc, AIC=iqtree_df$AIC, lnL=iqtree_df$LogL)
	n
	n = get_ndata_from_BIC(BIC=iqtree_df$BIC, AIC=iqtree_df$AIC, lnL=iqtree_df$LogL)
	n	
	'
	
	# Number of parameters
	k = round(-1*((AIC / -2) - lnL), digits=2)
	k
	return(k)
	} # END get_numparams_from_AIC <- function(AIC, lnL)


get_ndata_from_AICc <- function(AICc, AIC, lnL)
	{
	cmds='
	AICc = 16831.15
	AIC = 16331.08
	lnL = -8039.542
	k = get_numparams_from_AIC(AIC, lnL)
	k
	n = get_ndata_from_AICc(AICc, AIC, lnL)
	n

	iqtree_fn = "/GitHub/str2phy/ex/ferritins_M20af/03_USaln_v1/iqtree_3disTrim35/53fer.fasta.iqtree"
	iqtree_df = read_iqtree_model_scores(iqtree_fn, calc_extra=FALSE)
	k = get_numparams_from_AIC(AIC=iqtree_df$AIC, lnL=iqtree_df$LogL)
	k
	n = get_ndata_from_AICc(AICc=iqtree_df$AICc, AIC=iqtree_df$AIC, lnL=iqtree_df$LogL)
	n
	n = get_ndata_from_BIC(BIC=iqtree_df$BIC, AIC=iqtree_df$AIC, lnL=iqtree_df$LogL)
	n	
	' # END cmds
	derivation='
	# Number of data
	AICc = AIC + 2k(k+1) / (n-k-1)

	# Solve for n
	AICc - AIC = 2k(k+1) / (n-k-1)

	2k(k+1) / (AICc - AIC) = n-k-1

	(2k(k+1) / (AICc - AIC)) + k + 1 = n
	
	# BIC
	BIC = -2 * loglikelihood + d * log(N)
	BIC = -2lnL + k * log(n)
	exp((BIC + 2*lnL) / k) = n
	' # END derivation
	k = get_numparams_from_AIC(AIC, lnL)
	n = ( (2*k*(k+1)) / (AICc - AIC)) + k + 1
	n = round(n, digits=2)
	n
	return(n)
	} # END get_ndata_from_AICc <- function(AICc, AIC, lnL)

get_ndata_from_BIC <- function(BIC, AIC, lnL)
	{
	cmds='
	iqtree_fn = "/GitHub/str2phy/ex/ferritins_M20af/03_USaln_v1/iqtree_3disTrim35/53fer.fasta.iqtree"
	

	BIC = 16740.87
	AIC = 16331.08
	lnL = -8039.542
	k = get_numparams_from_AIC(AIC, lnL)
	k
	n = get_ndata_from_BIC(BIC, AIC, lnL)
	n
	
	iqtree_fn = "/GitHub/str2phy/ex/ferritins_M20af/03_USaln_v1/iqtree_3disTrim35/53fer.fasta.iqtree"
	iqtree_df = read_iqtree_model_scores(iqtree_fn, calc_extra=FALSE)
	k = get_numparams_from_AIC(AIC=iqtree_df$AIC, lnL=iqtree_df$LogL)
	k
	n = get_ndata_from_AICc(AICc=iqtree_df$AICc, AIC=iqtree_df$AIC, lnL=iqtree_df$LogL)
	n
	n = get_ndata_from_BIC(BIC=iqtree_df$BIC, AIC=iqtree_df$AIC, lnL=iqtree_df$LogL)
	n	
	' # END cmds
	derivation='
	# Number of data
	AICc = AIC + 2k(k+1) / (n-k-1)

	# Solve for n
	AICc - AIC = 2k(k+1) / (n-k-1)
	2k(k+1) / (AICc - AIC) = n-k-1
	(2k(k+1) / (AICc - AIC)) + k + 1 = n
	
	# BIC, solved for n
	BIC = -2 * loglikelihood + d * log(N)
	BIC = -2lnL + k * log(n)
	exp((BIC + 2*lnL) / k) = n
	' # END derivation
	k = get_numparams_from_AIC(AIC, lnL)
	n = exp((BIC + 2*lnL) / k)
	n = round(n, digits=2)
	n
	return(n)
	} # END get_ndata_from_AICc <- function(AICc, AIC, lnL)


read_iqtree_model_scores <- function(iqtree_fn, start_txt = "List of models sorted by BIC scores: ", calc_extra=TRUE, partitioned_TF=FALSE)
	{
	ex='
	iqtree_fn = "/GitHub/str2phy/ex/ferritins_M20af/03_USaln_v1/iqtree_3disTrim35/53fer.fasta.iqtree"
	start_txt = "List of models sorted by BIC scores: "
	calc_extra = TRUE  # back-calculate k and n
	partitioned_TF=FALSE

	iqtree_df = read_iqtree_model_scores(iqtree_fn)
	head(iqtree_df)
	iqtree_df = read_iqtree_model_scores(iqtree_fn, calc_extra=FALSE)
	head(iqtree_df)
	'	

	# Get the model results table
	lines = readLines(iqtree_fn)
	startline = 0
	endline = 0
	for (i in 1:length(lines))
		{
		if (startsWith(x=lines[i], prefix=start_txt) == TRUE)
			{
			startline = i+2
		
			for (j in startline:length(lines))
				{
				if (lines[j] == "")
					{
					endline = j-1
					break()
					}
				} # END for (j in startline:length(lines))
			break()
			} # END if (startsWith(x=lines[i], prefix=start_txt) == TRUE)
		} # END for (i in 1:length(lines))

	# Edit the header so it can be read, including the "+" "-" model-weights
	lines[startline] = stringi::stri_replace_first_fixed(str=lines[startline], pattern="AIC", replacement="AIC AICp")
	lines[startline]

	lines[startline] = stringi::stri_replace_first_fixed(str=lines[startline], pattern="AICc", replacement="AICc AICcp")
	lines[startline]

	lines[startline] = stringi::stri_replace_first_fixed(str=lines[startline], pattern="BIC", replacement="BIC BICp")
	lines[startline]

	writeLines(text=lines[startline:endline], con="tmpdf.txt")

	iqtree_df = read.table(file="tmpdf.txt", header=TRUE, sep="", stringsAsFactors=FALSE)
	head(iqtree_df)


	# Get the single-best-fit ML tree
	start_txt = "Tree in newick format:"
	startline = 0
	endline = 0
	for (i in 1:length(lines))
		{
		if (startsWith(x=lines[i], prefix=start_txt) == TRUE)
			{
			startline = i+2
		
			for (j in startline:length(lines))
				{
				if (lines[j] == "")
					{
					endline = j-1
					break()
					}
				} # END for (j in startline:length(lines))
			break()
			} # END if (startsWith(x=lines[i], prefix=start_txt) == TRUE)
		} # END for (i in 1:length(lines))
	writeLines(text=lines[startline:endline], con="tmp_MLtr.newick")
	tr = ape::read.tree("tmp_MLtr.newick")
	tr
	
	num_brlens = length(tr$edge.length)


	if (partitioned_TF == FALSE)
		{
		# Back-calculate some things
		if (calc_extra == TRUE)
			{
			k = get_numparams_from_AIC(AIC=iqtree_df$AIC, lnL=iqtree_df$LogL)
			k_wo_brlens = k - num_brlens
			ndata_AICc = get_ndata_from_AICc(AICc=iqtree_df$AICc, AIC=iqtree_df$AIC, lnL=iqtree_df$LogL)
			ndata_BIC = get_ndata_from_BIC(BIC=iqtree_df$BIC, AIC=iqtree_df$AIC, lnL=iqtree_df$LogL)
			iqtree_df = cbind(iqtree_df, k, k_wo_brlens, ndata_AICc, ndata_BIC)
			}
		}
	
	if (partitioned_TF == TRUE)
		{
		# Back-calculate some things
		if (calc_extra == TRUE)
			{
			iqtree_df_summed = iqtree_df[1,]
			iqtree_df_summed$Model = paste(iqtree_df$Model, collapse="; ")
			iqtree_df_summed$LogL = sum(iqtree_df$LogL)
			iqtree_df_summed$AIC = sum(iqtree_df$AIC)
			iqtree_df_summed$AICc = sum(iqtree_df$AICc)
			iqtree_df_summed$BIC = sum(iqtree_df$BIC)
			
			# These don't really work, so get directly...
			# total lnL
			TF = startsWith(x=lines, prefix="Log-likelihood of the tree: ")
			words = strsplit(lines[TF], split=":")[[1]]
			num = strsplit(gdata::trim(words[2]), split=" ")[[1]][1]
			iqtree_df_summed$LogL = as.numeric(num)

			TF = startsWith(x=lines, prefix="Number of free parameters")
			words = strsplit(lines[TF], split=":")[[1]]
			num = strsplit(gdata::trim(words[2]), split=" ")[[1]][1]
			k = as.numeric(num)
			k_wo_brlens = k - num_brlens

			# total AIC
			TF = startsWith(x=lines, prefix="Akaike information criterion (AIC) score:")
			words = strsplit(lines[TF], split=":")[[1]]
			num = strsplit(gdata::trim(words[2]), split=" ")[[1]][1]
			iqtree_df_summed$AIC = as.numeric(num)

			# total AICc
			TF = startsWith(x=lines, prefix="Corrected Akaike information criterion (AICc) score:")
			words = strsplit(lines[TF], split=":")[[1]]
			num = strsplit(gdata::trim(words[2]), split=" ")[[1]][1]
			iqtree_df_summed$AICc = as.numeric(num)

			# total AICc
			TF = startsWith(x=lines, prefix="Bayesian information criterion (BIC) score:")
			words = strsplit(lines[TF], split=":")[[1]]
			num = strsplit(gdata::trim(words[2]), split=" ")[[1]][1]
			iqtree_df_summed$BIC = as.numeric(num)

			ndata_AICc = get_ndata_from_AICc(AICc=iqtree_df_summed$AICc, AIC=iqtree_df_summed$AIC, lnL=iqtree_df_summed$LogL)
			ndata_BIC = get_ndata_from_BIC(BIC=iqtree_df_summed$BIC, AIC=iqtree_df_summed$AIC, lnL=iqtree_df_summed$LogL)
			iqtree_df = cbind(iqtree_df_summed, k, k_wo_brlens, ndata_AICc, ndata_BIC)
			}
		}
	
	return(iqtree_df)
	} # END read_iqtree_model_scores <- function(iqtree_fn)
```
