## Supplementary material for "Tertiary-interaction characters enable fast, model-based structural phylogenetics beyond the twilight zone": str2phy_v1.R

```
#######################################################
# Check tree for a bipartition
#######################################################
check_bipartition <- function(tr2, bipartition_tipnames, remove_txt="\\.pdb", tr2table=NULL)
	{
	junk='
	remove_txt="\\.pdb"
	
	trfn = "/GitHub/str2phy/ex/ferritins_M20/iqtree/full/iqtree_mset_FAMSAln_v1/partitions_v1.raxml.contree"
	tr2 = read.tree(trfn)
	bipartition_tipnames = c("1lko_A","1yuz_A","3qhb_A")
	
	source("/GitHub/str2phy/Rsrc/str2phy_v1.R")
	res = check_bipartition(tr2, bipartition_tipnames, remove_txt="\\.pdb")
	res
	'
	# Remove annoying text from tip labels
	orig_tr2_tiplabels = tr2$tip.label
	tr2_tiplabels = gsub(pattern=remove_txt, replacement="", x=tr2$tip.label)
	tr2_tiplabels = gsub(pattern="'", replacement="", x=tr2_tiplabels)
	tr2_tiplabels = gsub(pattern="`", replacement="", x=tr2_tiplabels)
	tr2_tiplabels
	
	tr2$tip.label = tr2_tiplabels
		
	bipartition_tipnums = match(x=bipartition_tipnames, table=tr2_tiplabels)
	# Sort, so that tip numbers match tip numbers in tree being searched
	bipartition_tipnums = sort(bipartition_tipnums)
	bipartition_tipnums

	numtips = length(bipartition_tipnums)
	tipnames = paste0(bipartition_tipnames, collapse=",")
	
	if (is.null(tr2table) == TRUE)
		{
		tr2table = prt(tr2, printflag=FALSE, get_tipnames=FALSE)
		}
	
	# bipartitions in 1 tree
	bipartitions_found = ape::prop.part(tr2)
	
	#  
	TF = sapply(X=bipartitions_found, FUN=identical, y=bipartition_tipnums)
	sum(TF)
	
	if (sum(TF) == 1)
		{
		bipart = 1
		} else {
		bipart = 0
		}

	
	# If the bipartition exists in the searched tree, get the branch length and support value, if present
	# Node in the searched tree
	nodenum = getMRCA(phy=tr2, tip=bipartition_tipnames)
	
	# Check if the node matches the number of tips
	daughter_tips = get_daughter_tipnames(t=tr2, node=nodenum)

	if (identical(sort(daughter_tips), sort(bipartition_tipnames)) == TRUE)
		{
		edgenum = which(tr2$edge[,2]==nodenum)
		brlen2 = tr2$edge.length[edgenum]
		if (length(brlen2) == 0)
			{
			brlen2 = NA
			}
		# Get the node label, if any
		# (you have to subtract the number of tips)
		nodenum2 = nodenum - length(tr2$tip.label)
		bootstrap2 = tr2$node.label[nodenum2]
		if (length(bootstrap2) == 0)
			{
			bootstrap2 = NA
			}

		# Record node_ht above midpoint root, as well as proportional height relative to highest tip
		node_ht = tr2table$node_ht[nodenum]
		rel_ht = tr2table$node_ht[nodenum] / max(tr2table$node_ht)
		} else {
		brlen2 = NA
		bootstrap2 = NA
		node_ht = NA
		rel_ht = NA
		}
	
	res = c(numtips, bipart, brlen2, bootstrap2, node_ht, rel_ht, tipnames)
	names(res) = c("numtips", "bipart", "brlen2", "bootstrap2", "node_ht", "rel_ht", "tipnames")
	return(res)
	}


check_bipartitions_in_tree2 <- function(tr1, tr2, remove_txt="\\.pdb")
	{
	bipartitions_list_in_distance_tr = ape::prop.part(tr1)
	orig_tr_tiplabels = tr1$tip.label
	tr1_tiplabels = gsub(pattern=remove_txt, replacement="", x=tr1$tip.label)
	tr1_tiplabels = gsub(pattern="'", replacement="", x=tr1_tiplabels)
	tr1_tiplabels = gsub(pattern="`", replacement="", x=tr1_tiplabels)
	tr1_tiplabels
	
	tr2table = prt(tr2, printflag=FALSE, get_tipnames=FALSE)
	
	dtf = NULL
	for (i in 1:length(bipartitions_list_in_distance_tr))
		{
		bipartition_tipnums = bipartitions_list_in_distance_tr[[i]]
		bipartition_tipnames = sort(tr1_tiplabels[bipartition_tipnums])
		
		res = check_bipartition(tr2, bipartition_tipnames, remove_txt=remove_txt, tr2table=tr2table)
		dtf = rbind(dtf, res)
		}
	
	dtf = as.data.frame(dtf, stringsAsFactors=FALSE)
	names(dtf) = c("numtips", "bipart", "brlen2", "bootstrap2", "node_ht", "rel_ht", "tipnames")

	dtf = dfnums_to_numeric(dtf)
	dtf = unlist_df3(dtf)
	return(dtf)	
	}


#######################################################
# Align each 3di sequence to the US-aligned version
#######################################################
cmds='
library(msa)
library(seqinr)
library(BioGeoBEARS)
sourceall("/GitHub/bioinfRhints/Rsrc")
source("/GitHub/str2phy/Rsrc/str2phy_v1.R")


wd = "/GitHub/str2phy/ex/CDhits_subset/"
setwd(wd)

# Specify file locations
fasta_fn = "/Users/nickm/GitHub/bioinfRhints/flag/AQB_classification/1283_AQBs.fasta"
usaln_fn = "usalign_test1_aa.fasta"
di3s_dir = "/GitHub/str2phy/aln_3di_to_aln"

# Read in files
raw_seqs = as.character(ape::read.FASTA(fasta_fn, type="AA"))
raw_names = unname(sapply(X=names(raw_seqs), FUN=firstword))

usaln_seqs = as.character(ape::read.FASTA(usaln_fn, type="AA"))
usaln_names = names(usaln_seqs)
usaln_names = gsub(pattern="WP_", replacement="WP|", x=tmpnames)
for (i in 1:length(usaln_names))
	{
	words = strsplit(usaln_names[i], split="_")[[1]]
	usaln_names[i] = words[1]
	}
usaln_names = gsub(pattern="WP\\|", replacement="WP_", x=usaln_names)
usaln_names

di3s_fns = list.files(di3s_dir, pattern="*_3di.fasta", full.names=TRUE)
di3s_fns

#######################################################
# Align each 3di sequence to the US-aligned version
#######################################################
i=1
aln_seq = paste0(usaln_seqs[[i]], collapse="")
raw_seq = paste0(raw_seqs[[i]], collapse="")
conversion_table = aln_2nd_seq_to_1st(aln_seq, raw_seq)
conversion_table
	
'


aln_2nd_seq_to_1st <- function(aln_seq, raw_seq)
	{
	pairwise_aln = msa(inputSeqs=c(aln_seq, raw_seq), method="ClustalW", type="protein", order="input")
	pairwise_aln2 = msaConvert(pairwise_aln, type="ape::AAbin")
	pairwise_aln3 = add_labels_to_AAbin(pairwise_aln2)
	names(pairwise_aln2)
	names(pairwise_aln3)
	seqs_list = as.character(pairwise_aln3)

	#######################################################
	# Convert the non-blank positions of the 2nd sequence into positions in the 1st sequence
	#######################################################
	observed_positions = 1:length(seqs_list[[2]])

	TF = seqs_list[[2]] != "-"
	nonblank_positions = rep(NA, times=length(seqs_list[[2]]))
	nonblank_positions[TF] = 1:sum(TF)

	aligned_seq_pos = observed_positions
	unaligned_seq_pos = nonblank_positions
	
	conversion_table = rbind(aligned_seq_pos, unaligned_seq_pos)
	#conversion_table[,400:410]
	return(conversion_table)
	}


# Writes to FASTA to add labels, reads back in
add_labels_to_AAbin <- function(aln, outfn = "tmp_addlabels.fasta")
	{
	cmds='
	library(msa)
	library(seqinr)
	library(BioGeoBEARS)

a="LSVWGMYQHADIVVKCVMIGLILASVVTWAIFFSKSVEFFNQKRRLKREQQLLAEARSLNQANDIAADFGSKSLSLHLLNEAQNELELSEGSDDNEGIKERTSFRLERRVAAVGRQMGRGNGYLATIGAISPFVGLFGTVWGIMNSFIGIAQTQTTNLAVVAPGIAEALLATAIGLVAAIPAVVIYNVFARQIGGFKAMLGDVAAQVLLLQSRDLDLEASAAAHP"
	b="DLSIWGMYQHADAVVKAVMIGLVLASIVTWTILFAKGSELLRAKRRLRREQLALAEARSLDEASELAQNFSPESVSAVLLNDAQNELELSAESNDNNGIKERTGFRLERRVAAYSRNMGRGNGFLATIGAISPFVGLFGTVWGIMNSFIGIAHSQTTNLAVVAPGIAEALLATAMGLVAAIPAVVIYNIFARVISGHRAQVGDVAAQVLLLQGRDLDLAATAEAKRSQHAHQLRAG"

	# Align with ClustalW
	aln = msa(inputSeqs=c(a,b), method="ClustalW", type="protein", order="input")

	aln2 = msaConvert(aln, type="ape::AAbin")
	aln2
	names(aln2)
	labels(aln2)

	outfn = "tmp_addlabels.fasta"
	aln3 = add_labels_to_AAbin <- function(aln, outfn = "tmp_addlabels.fasta")
	names(aln3)
	labels(aln3)

	'
	ape::write.FASTA(x=aln, file=outfn, header=NULL, append=FALSE)
	aln = ape::read.FASTA(file=outfn, type="AA")
	return(aln)
	}

fasta_to_alphafold <- function(fasta_fn, seqnum=1)
	{
	seqs = ape::read.FASTA(fasta_fn, type="AA")
	seqs

	header = names(seqs)[[seqnum]]
	gid = firstword(header)

	html_fn = paste0(gid, "_alphafold.html")
	cif_fn = paste0(gid, "_alphafold.cif")
	table_fn = paste0(gid, "_alphafold_table.html")

	if (file.exists(html_fn) == TRUE)
		{
		# Check for "no hit"
		TF = grepl(pattern="No AlphaFold model with similar sequence", x=readLines(html_fn))
		if (sum(TF) > 0)
			{
			fns = c("NONE", NA, 0.0, 0.0, NA, html_fn, NA, NA, NA, NA, NA, NA)
			outdf = as.data.frame(matrix(fns, nrow=1), stringsAsFactors=FALSE)
			row.names(outdf) = NULL
			names(outdf) = c("AlphaFold Model", "Query Sequence", "Identity %", "Coverage %", "cif_fn", "html_fn", "table_fn", "dump_fn", "chain_names_fn", "di3_fn", "aa_fn", "both_fn")
			outdf = cbind(seqnum, outdf)
			return(outdf)
			}
		
		
		alphafold_df = alphafold_html_to_df(html_fn, table_fn)
		alphafold_df
	
		# Structure to 3di and fasta
		dump_fn = gsub(pattern="\\.cif", replacement="_3di.dump", x=cif_fn)
		chain_names_fn = gsub(pattern="\\.cif", replacement="_chains.txt", x=cif_fn)
		di3_fn = gsub(pattern="\\.cif", replacement="_3di.fasta", x=cif_fn)
		aa_fn = gsub(pattern="\\.cif", replacement="_aa.fasta", x=cif_fn)
		both_fn = gsub(pattern="\\.cif", replacement="_both.fasta", x=cif_fn)

		fns = c(cif_fn, html_fn, table_fn, dump_fn, chain_names_fn, di3_fn, aa_fn, both_fn)
		fns_df = as.data.frame(matrix(fns, nrow=1), stringsAsFactors=FALSE)
		row.names(fns_df) = NULL
		names(fns_df) = c("cif_fn", "html_fn", "table_fn", "dump_fn", "chain_names_fn", "di3_fn", "aa_fn", "both_fn")
	
		outdf = cbind(alphafold_df, fns_df)
		outdf = cbind(seqnum, outdf)
		} else {
		chars = as.character(seqs)[[seqnum]]
		outdf = aa_to_alphafold(chars, gid)
		outdf = cbind(seqnum, outdf)
		} # END if (file.exists(html_fn) == TRUE)
	return(outdf)
	}
	
aa_to_alphafold <- function(chars, gid, timeout=30)
	{
	aastr = toupper(paste0(chars, collapse="", sep=""))
	aastr
	
	html_fn = paste0(gid, "_alphafold.html")
	cif_fn = paste0(gid, "_alphafold.cif")
	table_fn = paste0(gid, "_alphafold_table.html")
	
	# Download closest-match alphafold structure
	keep_going = TRUE
	start_time = Sys.time()
	first_time = TRUE
	while(keep_going == TRUE)
		{
		if (first_time == TRUE)
			{
			cmd_txt = paste0('ChimeraX --cmd "alphafold match ', aastr, '; log save ', html_fn, ' executableLinks true; save ', cif_fn, ' relModel #1; close #1; quit;"')
			cmd_txt
			#result = try(system(cmd_txt))
			first_time = FALSE
			system(cmd_txt, wait=TRUE, timeout=timeout)
			}
		
		if (file.exists(cif_fn) == TRUE)
			{
			keep_going = FALSE
			break()
			}
		
		current_time = Sys.time()
		elapsed_time = as.numeric(current_time - start_time)
		if (elapsed_time > (timeout+5))
			{
			outdf = rep(NA, times=12)
			return(outdf)
			keep_going = FALSE
			}
		} # END while-loop
				

	
	# Wait until cif_fn exists
	keep_going_TF = TRUE
	cat("\nRunning: ", cmd_txt)
	cat("\n")
	while (keep_going_TF == TRUE)
		{
		if ( (file.exists(cif_fn) == TRUE) && (file.exists(html_fn) == TRUE) )
			{
			keep_going_TF = FALSE
			}
		}
	cat("...done.\n")
	
	alphafold_df = alphafold_html_to_df(html_fn, table_fn)
	alphafold_df
	
	# Structure to 3di and fasta
	dump_fn = gsub(pattern="\\.cif", replacement="_3di.dump", x=cif_fn)
	chain_names_fn = gsub(pattern="\\.cif", replacement="_chains.txt", x=cif_fn)
	di3_fn = gsub(pattern="\\.cif", replacement="_3di.fasta", x=cif_fn)
	aa_fn = gsub(pattern="\\.cif", replacement="_aa.fasta", x=cif_fn)
	both_fn = gsub(pattern="\\.cif", replacement="_both.fasta", x=cif_fn)
	
	cmd_txt = paste0("foldseek structureto3didescriptor ", cif_fn, " ", dump_fn)
	system(cmd_txt)
	
	cmd_txt = paste0("awk -F'\t' '{print $1; next}{print}' ", dump_fn, " | tee ", chain_names_fn)
	system(cmd_txt)
	
	cmd_txt = paste0("awk -F'\t' '{print ", '">"', '$1"\\n"', "$3; next}{print}' ", dump_fn, " | tee ", di3_fn)
	system(cmd_txt)

	#cmd_txt2 = paste0("-F'\t' '{print ", '">"', '$1"\\n"', "$3; next}{print}' ", dump_fn, " | tee ", di3_fn)
	#system2(command="awk", args=cmd_txt2)

	cmd_txt = paste0("awk -F'\t' '{print ", '">"', '$1"\\n"', "$2; next}{print}' ", dump_fn, " | tee ", aa_fn)
	system(cmd_txt)
	
	cmd_txt = paste0("awk -F'\t' '{print ", '">"', '$1"\\n"', '$2"\\n"', "$3; next}{print}' ", dump_fn, " | tee ", both_fn)
	system(cmd_txt)
	
	
	# fns
	fns = c(cif_fn, html_fn, table_fn, dump_fn, chain_names_fn, di3_fn, aa_fn, both_fn)
	fns_df = as.data.frame(matrix(fns, nrow=1), stringsAsFactors=FALSE)
	row.names(fns_df) = NULL
	names(fns_df) = c("cif_fn", "html_fn", "table_fn", "dump_fn", "chain_names_fn", "di3_fn", "aa_fn", "both_fn")
	
	outdf = cbind(alphafold_df, fns_df)
	outdf
	
	return(outdf)
	}


alphafold_html_to_df <- function(html_fn, table_fn)
	{
	R_cmds='
	wd = "/GitHub/str2phy/aln_3di_to_aln/"
	setwd(wd)
	html_fn = "AAQ60658.1_alphafold.html"
	table_fn = "AAQ60658.1_alphafold_table.html"
	alphafold_df = alphafold_html_to_df(html_fn, table_fn)
	alphafold_df
	'
	
	
	txt_cmds="
	cd /GitHub/str2phy/aln_3di_to_aln/
	tr '\n' ' ' < AAQ60658.1_alphafold.html | awk -F 'table' '$2{print $2}' > AAQ60658.1_alphafold_table.html
	"	
	
	# Extract just the table to a text file
	cmdtxt = paste0("tr '\n' ' ' < ", html_fn, " | awk -F 'table' '$2{print $2}' > ", table_fn)
	cmdtxt
	system(cmdtxt)
	
	# Parse the text
	tmpline = readLines(table_fn)
	gsub(pattern='<td style=\\"text-align:center\\">', replacement="\t", x=tmpline)


	tmpline = gsub(pattern=' border=1 cellpadding=4 cellspacing=0>', replacement="\t", x=tmpline)

	tmpline = gsub(pattern='<td style=\\"text-align:center\\">', replacement="\t", x=tmpline)
	tmpline = gsub(pattern="<thead>", replacement="\t", x=tmpline)
	tmpline = gsub(pattern="</thead>", replacement="\t", x=tmpline)
	tmpline = gsub(pattern="<tbody>", replacement="\t", x=tmpline)
	tmpline = gsub(pattern="</tbody>", replacement="\t", x=tmpline)
	tmpline = gsub(pattern="<th colspan=4>", replacement="\t", x=tmpline)
	tmpline = gsub(pattern="<th>", replacement="\t", x=tmpline)
	tmpline = gsub(pattern="<tr>", replacement="\t", x=tmpline)
	tmpline = gsub(pattern="td", replacement="\t", x=tmpline)
	tmpline = gsub(pattern="</tr>", replacement="\t", x=tmpline)
	tmpline = gsub(pattern="</td>", replacement="\t", x=tmpline)
	tmpline = gsub(pattern="</", replacement="\t", x=tmpline)
	
	tmpline = gsub(pattern="\t\t", replacement="\t", x=tmpline)
	tmpline = gsub(pattern="\t\t", replacement="\t", x=tmpline)
	tmpline = gsub(pattern="\t\t", replacement="\t", x=tmpline)
	tmpline = gsub(pattern="\t\t", replacement="\t", x=tmpline)
	tmpline = gsub(pattern="\t\t", replacement="\t", x=tmpline)
	tmpline = gsub(pattern="\t\t", replacement="\t", x=tmpline)
	tmpline = gsub(pattern="\t\t", replacement="\t", x=tmpline)
	tmpline = gsub(pattern="\t\t", replacement="\t", x=tmpline)
	tmpline = gsub(pattern="\t\t", replacement="\t", x=tmpline)
	tmpline = gsub(pattern="\t\t", replacement="\t", x=tmpline)
	tmpline = gsub(pattern="\t\t", replacement="\t", x=tmpline)
	tmpline = gsub(pattern="\t\t", replacement="\t", x=tmpline)
	tmpline = gsub(pattern="\t\t", replacement="\t", x=tmpline)
	tmpline = gsub(pattern="\t\t", replacement="\t", x=tmpline)
	tmpline
	
	words = strsplit(tmpline, split="\t")[[1]]
	words = gdata::trim(words)
	words = words[words != ""]
	words = words[2:9]
	
	tmpmat = matrix(data=words[5:8], nrow=1)
	alphafold_df = as.data.frame(tmpmat, stringsAsFactors=FALSE)
	row.names(alphafold_df) = NULL
	names(alphafold_df) = words[1:4]
	return(alphafold_df)
	}


#######################################################
# Process a CD-hits (conserved domain) xlxs table
# (which has been manually processed to have a max of 2 domain hits)
#######################################################
cmds='
wd = "/GitHub/bioinfRhints/flag/AQB_classification/batch_CD_CDART/"
setwd(wd)

library(openxlsx)			# for read.xlsx


xlsfn = "/GitHub/bioinfRhints/flag/AQB_classification/groupTax_1282_mafftConstr_2023-08-07_edit.xlsx"
xls = read.xlsx(xlsfn)


cdhits_fn = "1283_AQBs_gids_hitdata_v2merge34.xlsx"
cdxls = read.xlsx(cdhits_fn)
numhits_by_gid = rev(sort(table(cdxls$Query)))

core_domains = c("MotA_ExbB superfamily", "TolQ superfamily", "MotA superfamily")
process_CDhits(cdxls, xls, max_core_length=325, core_domains=core_domains)

'

process_CDhits <- function(cdxls, xls, max_core_length=325, core_domains=c("MotA_ExbB superfamily", "TolQ superfamily", "MotA superfamily"))
	{
	#core_domains = c("MotA_ExbB superfamily", "TolQ superfamily", "MotA superfamily")

	uniq_CD_names = names(rev(sort(table(cdxls$Short.name))))

	gids_in_xls = rep("", times=nrow(xls))
	core_start = rep(NA, times=nrow(xls))
	core_stop = rep(NA, times=nrow(xls))

	alt1_start = rep(NA, times=nrow(xls))
	alt1_stop = rep(NA, times=nrow(xls))
	alt2_start = rep(NA, times=nrow(xls))
	alt2_stop = rep(NA, times=nrow(xls))
	alt3_start = rep(NA, times=nrow(xls))
	alt3_stop = rep(NA, times=nrow(xls))
	alt4_start = rep(NA, times=nrow(xls))
	alt4_stop = rep(NA, times=nrow(xls))

	num_domains = rep(0, times=nrow(xls))
	protein_lens = rep(0, times=nrow(xls))
	core_dom_names = rep("", times=nrow(xls))
	alt1_dom_names = rep("", times=nrow(xls))
	alt2_dom_names = rep("", times=nrow(xls))
	alt3_dom_names = rep("", times=nrow(xls))
	alt4_dom_names = rep("", times=nrow(xls))

	i = 1
	uniq_gids = unique(cdxls$Query)
	
	cat("\nProcessing CD-hits for ", length(uniq_gids), " unique gids in ", nrow(cdxls), " domain hits in cdxls table...\n")
	for (i in 1:length(uniq_gids))
		{
		cat(i, ",", sep="")
		gid = uniq_gids[i]

		TF = xls$tipnames3_uniq == gid
		xlsrownum = (1:length(TF))[TF]
		xlsrow = xls[xlsrownum,]
		gids_in_xls[xlsrownum] = gid
		
		
		TF = cdxls$Query == gid
		cdrows = cdxls[TF,]

		TF = cdrows$Short.name %in% core_domains
		core_rows = cdrows[TF,]
		noncore_rows = cdrows[TF==FALSE,]

		# Initialize each loop
		protein_len = xls$len[xlsrownum]
		core_hit_start = 0
		core_hit_stop = 0
		alt1_hit_start = 0
		alt1_hit_stop = 0
		core_dom_name = ""
		alt1_dom_name = ""

		if (nrow(core_rows) < 1)
			{
			num_domains[xlsrownum] = 0
			core_hit_start = 1
			core_hit_stop = protein_len
			core_dom_name = NA
			alt1_dom_name = NA
			}

		if ((nrow(core_rows) == 1) && (nrow(cdrows)==1))
			{
			core_dom_name = core_rows$Short.name[1]
			num_domains[xlsrownum] = 1

			core_hit_start = core_rows$From[1]
			core_hit_stop = core_rows$To[1]
			core_length = core_hit_stop - core_hit_start + 1
	
			# Split a very-long protein
			if ((protein_len) > 1.5*max_core_length)
				{
				num_domains[xlsrownum] = 2
				alt1_dom_name = "unknown noncore sequence"
		
				# Which side of the protein is the core on?
				middle_of_protein = round(protein_len/2)
				middle_of_core = (core_hit_start + core_hit_stop) / 2

				if (middle_of_core >= middle_of_protein)
					{
					core_hit_start_option1 = core_hit_start
					core_hit_start_option2 = protein_len - max_core_length + 1
					
					core_hit_start = min(core_hit_start_option1, core_hit_start_option2)
					core_hit_stop = protein_len
					alt1_hit_start = 1
					alt1_hit_stop = core_hit_start - 1
					} else {
					core_hit_stop_option1 = core_hit_stop
					core_hit_stop_option2 = core_hit_stop
					
					core_hit_start = 1
					core_hit_stop = max_core_length
					alt1_hit_start = core_hit_stop + 1
					alt1_hit_stop = protein_len	
					}
				} else {
				alt1_dom_name = ""
				core_hit_start = 1
				core_hit_stop = protein_len
				} # END if ((protein_len) > max_core_length)
			} # END if ((nrow(core_rows) == 1) && (nrow(cdrows)==1))


		if ((nrow(core_rows) == 2) && (nrow(cdrows)==2))
			{
			core_dom_name = core_rows$Short.name[1]
			num_domains[xlsrownum] = 1
			core_start[xlsrownum] = 1
			core_stop[xlsrownum] = protein_len
			next()
			} # END if ((nrow(core_rows) == 2) && (nrow(cdrows)==2))

		if ((nrow(core_rows) == 1) && (nrow(cdrows)==2))
			{
			core_dom_name = core_rows$Short.name[1]
			alt1_dom_name = noncore_rows$Short.name[1]
			num_domains[xlsrownum] = 2
	
			core_hit_start = core_rows$From[1]
			core_hit_stop = core_rows$To[1]
	
			alt1_hit_start = noncore_rows$From[1]
			alt1_hit_stop = noncore_rows$To[1]
	
	
			# Which side of the protein is the core on?
			middle_of_protein = round(protein_len/2)
			middle_of_core = (core_hit_start + core_hit_stop) / 2
	
			if (middle_of_core >= middle_of_protein)
				{
				start_of_core_option1 = core_hit_start
				start_of_core_option2 = core_hit_stop - max_core_length
				if (start_of_core_option2 <= alt1_hit_stop)
					{
					start_of_core_option2 = alt1_hit_stop + 1
					core_hit_start = max(c(start_of_core_option1, start_of_core_option2))
					core_hit_stop = protein_len
			
					alt1_hit_start = 1
					alt1_hit_stop = core_hit_start - 1
					} else {
					core_hit_start = min(c(start_of_core_option1, start_of_core_option2))
					core_hit_stop = protein_len
			
					alt1_hit_start = 1
					alt1_hit_stop = core_hit_start - 1
					}
				} # END if (middle_of_core >= middle_of_protein)
		
			if (middle_of_core < middle_of_protein)
				{
				stop_of_core_option1 = core_hit_stop
				stop_of_core_option2 = max_core_length
				if (start_of_core_option2 <= alt1_hit_start)
					{
					stop_of_core_option2 = alt1_hit_start - 1
					core_hit_start = 1
					core_hit_stop = min(c(stop_of_core_option1, stop_of_core_option2))
			
					alt1_hit_start = core_hit_stop + 1
					alt1_hit_stop = protein_len
					} else {
					core_hit_start = 1
					core_hit_stop = max(c(stop_of_core_option1, alt1_hit_start-1))
			
					alt1_hit_start = core_hit_stop + 1
					alt1_hit_stop = protein_len
					}
				} # END if (middle_of_core < middle_of_protein)
			} # END if ((nrow(core_rows) == 1) && (nrow(cdrows)==2))

		protein_lens[xlsrownum] = protein_len
		core_dom_names[xlsrownum] = core_dom_name
		alt1_dom_names[xlsrownum] = alt1_dom_name
		alt2_dom_names[xlsrownum] = ""
		alt3_dom_names[xlsrownum] = ""
		alt4_dom_names[xlsrownum] = ""
		core_start[xlsrownum] = core_hit_start
		core_stop[xlsrownum] = core_hit_stop
		alt1_start[xlsrownum] = alt1_hit_start
		alt1_stop[xlsrownum] = alt1_hit_stop
		} # END for (i in 1:length(uniq_gids))
	cat("\n...done.\n")
	
	mat = cbind(gids_in_xls, num_domains, protein_lens, core_dom_names, core_start, core_stop, alt1_dom_names, alt1_start, alt1_stop)
	cdhits_df = as.data.frame(mat, stringsAsFactors=FALSE)
	row.names(cdhits_df) = NULL
	names(cdhits_df) = c("gid_for_CD", "num_domains", "protein_lens", "core_dom_names", "core_start", "core_stop", "alt1_dom_names", "alt1_start", "alt1_stop")
	return(cdhits_df)
	} # END process_CDhits


#######################################################
# Horizontally cat fasta_fn2 to the end of fasta_fn1
# (fasta_fn2 names must be grepl matches to names of fasta_fn1
#######################################################
hcat_fastas <- function(fasta_fn1, fasta_fn2, outfn=NULL, type="AA")
	{
	junk='
	wd = "/GitHub/str2phy/ex/ferritins_M20/align/02_FAMSA_aln_v1/"
	setwd(wd)

	fasta_fn1 = "53_ferritin_AAs.fasta.aln"
	fasta_fn2 = "53_ferritin_3dis.fasta.aln"

	outfn=NULL
	type="AA"
	
	aa3di_alignment = hcat_fastas(fasta_fn1, fasta_fn2, outfn=NULL, type="AA")
	aa3di_alignment
	'
	
	
	aln_aaBin = ape::read.FASTA(fasta_fn1, type=type)
	aa_chars = as.character(aln_aaBin)

	aln_aaBin2 = ape::read.FASTA(fasta_fn2, type=type) 
	chars_aa_to_add = as.character(aln_aaBin2)

	#######################################################
	# Align each 3di sequence to the US-aligned version
	#######################################################
	i=1

	# Go through the USaligned-alignment
	# find the 3di FASTA file, load, parse, align, add to alignment
	aa3di_alignment_list = list()
	for (i in 1:length(aa_chars))
		{
		aln_name = names(aa_chars)[i]
		aln_seq = aa_chars[[i]]

		TF = grepl(pattern=aln_name, x=names(chars_aa_to_add))

		if (sum(TF) != 1)
			{
			txt = paste0("STOP ERROR in hcat_fastas(): no matching name found for alignment entry aln_name='", aln_name, "', in names(chars_aa_to_add). Please fix and re-run.")
			cat("\n")
			cat(txt)
			cat("\n")
			stop(txt)
			}
		
		listnum = (1:length(chars_aa_to_add))[TF]
		add_chars = chars_aa_to_add[[listnum]]

		aa3di_alignment_list[[aln_name]] = c(aa_chars[[aln_name]], add_chars)
		} # END for (i in 1:length(aa_chars))

	if (type == "AA")
		{
		aa3di_alignment = ape::as.AAbin(aa3di_alignment_list)
		} else {
		aa3di_alignment = ape::as.DNAbin(aa3di_alignment_list)
		}
	aa3di_alignment
	
	if (is.null(outfn))
		{
		outfn = gsub(pattern="\\.fasta", replacement="", x=outfn)
		outfn = paste0(outfn, "_plus.fasta")
		}
	ape::write.FASTA(aa3di_alignment, file=outfn)
	return(aa3di_alignment)
	}


#######################################################
# Reorder fasta_fn to match names order in tip_txt
# (fasta_fn names must be grepl matches to names of tip_txt
#######################################################
reorder_fasta <- function(fasta_fn, tip_txt, outfn=NULL, type="AA")
	{
	# By hand:
	# tr$tip.label = gsub(pattern="'", replacement="", x=tr$tip.label)
	
	junk='
	library(ape)
	library(seqinr)
	library(BioGeoBEARS)
	sourceall("/GitHub/bioinfRhints/Rsrc")
	source("/GitHub/str2phy/Rsrc/str2phy_v1.R")


	wd = "/GitHub/str2phy/ex/ferritins_M20/align/02_FAMSA_aln_v1/"
	setwd(wd)

	fasta_fn = "53_ferritin_AAs.fasta.aln"

	outfn=NULL
	type="AA"
	
	trfn = "53_ferritin_both.fasta.aln.contree.newick"
	tr = read.tree(trfn)
	# By hand:
	# #NOTE: PASTE MANUAL ABOVE!!
	tip_txt = tr$tip.label

	outfn = gsub(pattern="\\.fasta", replacement="", x=fasta_fn)
	outfn = paste0(outfn, "_reord.fasta")
	
	alignment = reorder_fasta(fasta_fn, tip_txt=tip_txt, outfn=outfn, type="AA")
	alignment
	moref(outfn)
	' # END
	
	aln_aaBin = ape::read.FASTA(fasta_fn, type=type)
	aa_chars = as.character(aln_aaBin)
	aln_names = names(aa_chars)
	
	#######################################################
	# Align each 3di sequence to the US-aligned version
	#######################################################
	i=1
	alignment_list = list()
	for (i in 1:length(tip_txt))
		{
		TF = grepl(pattern=tip_txt[i], x=aln_names)
		seqnum = (1:length(aa_chars))[TF]
		add_chars = aa_chars[[seqnum]]
		aln_name = aln_names[seqnum]
		alignment_list[[aln_name]] = add_chars
		}
	
	if (type == "AA")
		{
		alignment = ape::as.AAbin(alignment_list)
		} else {
		alignment = ape::as.DNAbin(alignment_list)
		}
	alignment
	
	if (is.null(outfn))
		{
		outfn = gsub(pattern="\\.fasta", replacement="", x=outfn)
		outfn = paste0(outfn, "_reord.fasta")
		}
	ape::write.FASTA(alignment, file=outfn)
	return(alignment)
	}


#######################################################
# Align a 3di FASTA file to a matching pre-aligned AA FASTA entry
#######################################################
align_3di_to_aa <- function(chars_aa_aligned, chars_3di)
	{
	di3_aligned = rep("-", times=length(chars_aa_aligned))
	colnums_TF = chars_aa_aligned != "-"
	colnums = (1:length(colnums_TF))[colnums_TF]
	di3_aligned[colnums] = chars_3di
	return(di3_aligned)
	}


cmds='
library(ape)
library(seqinr)
library(BioGeoBEARS)
sourceall("/GitHub/bioinfRhints/Rsrc")
source("/GitHub/str2phy/Rsrc/str2phy_v1.R")

wd = "/GitHub/str2phy/ex/ferritins_M20/align/02_FAMSA_aln_v1/"
setwd(wd)

aligned_fn = "53_ferritin_3dis.fasta.aln"
unaligned_fn = "53_ferritin_AAs.fasta"
outfn = paste0(unaligned_fn, ".aln")
di3_alignment = align_3dis_to_AAs(aligned_fn, unaligned_fn, outfn)
'

align_3dis_to_AAs <- function(aligned_fn, unaligned_fn, outfn)
	{
	aln_aaBin = ape::read.FASTA(aligned_fn, type="AA")
	chars_aa_aligned = as.character(aln_aaBin)

	aln_aaBin2 = ape::read.FASTA(unaligned_fn, type="AA") 
	chars_aa_unaligned = as.character(aln_aaBin2)
	
	#######################################################
	# Align each 3di sequence to the US-aligned version
	#######################################################
	i=1

	# Go through the USaligned-alignment
	# find the 3di FASTA file, load, parse, align, add to alignment
	di3_alignment_list = list()
	for (i in 1:length(chars_aa_aligned))
		{
		aln_name = names(chars_aa_aligned)[i]
		aln_seq = chars_aa_aligned[[i]]
		aln_positions = 1:length(aln_seq)
		aln_positions = aln_positions[aln_seq != "-"]
		aln_positions

		TF = grepl(pattern=aln_name, x=names(chars_aa_unaligned))

		if (sum(TF) != 1)
			{
			txt = paste0("STOP ERROR in align_3dis_to_AAs(): no matching name found for alignment entry aln_name='", aln_name, "', in names(chars_aa_unaligned). Please fix and re-run.")
			cat("\n")
			cat(txt)
			cat("\n")
			stop(txt)
			}
		
		listnum = (1:length(chars_aa_unaligned))[TF]
		unaln_seq = chars_aa_unaligned[[listnum]]
		
		di3_aligned = align_3di_to_aa(aln_seq, unaln_seq)

		di3_alignment_list[[aln_name]] = di3_aligned
		}

	di3_alignment_list

	di3_alignment = ape::as.AAbin(di3_alignment_list)
	di3_chars = as.character(di3_alignment)
	
	if (is.null(outfn))
		{
		outfn = paste0(unaligned_fn, ".aln")
		}
	cat("\nalign_3dis_to_AAs() is writing aligned characters to: '", outfn, "'.\n", sep="")
	ape::write.FASTA(di3_alignment, file=outfn)

	return(di3_alignment)
	}


align_3dis <- function(usaln_fn, di3s_dir, di3_outfn, pattern_3di_fn="*_3di.fasta")
	{
	ex='
	usaln_fn = "alphafolds/usalign_9_MotBs3.fasta"
	di3s_dir = "/GitHub/str2phy/ex/MotB/alphafolds/"
	di3_outfn = "motB_seqs_first10_3di_aln.fasta"
	pattern_3di_fn="*_3di.fasta"
	di3_alignment = align_3dis(usaln_fn, di3s_dir, di3_outfn, pattern_3di_fn="*_3di.fasta")
	'


	# Read in files
	usaln_aaBin = ape::read.FASTA(usaln_fn, type="AA")
	numcols = max(as.numeric(summary(usaln_aaBin)[,"Length"]))
	usaln_seqs = as.character(usaln_aaBin)
	usaln_names = names(usaln_seqs)
	usaln_names = gsub(pattern="WP_", replacement="WP.", x=usaln_names)
	usaln_names = gsub(pattern="VER_", replacement="VER.", x=usaln_names)
	usaln_names_orig = usaln_names

	for (i in 1:length(usaln_names))
		{
		words = strsplit(usaln_names[i], split="_")[[1]]
		usaln_names[i] = words[1]
		words = strsplit(usaln_names[i], split="\\|")[[1]]
		usaln_names[i] = words[1]
		}
	usaln_names = gsub(pattern="WP\\.", replacement="WP_", x=usaln_names)
	usaln_names = gsub(pattern="VER\\.", replacement="VER_", x=usaln_names)
	usaln_names

	di3_fns = slashslash(list.files(di3s_dir, pattern=pattern_3di_fn, full.names=TRUE))
	di3_fns


	#######################################################
	# Align each 3di sequence to the US-aligned version
	#######################################################
	i=1

	# Go through the USaligned-alignment
	# find the 3di FASTA file, load, parse, align, add to alignment
	di3_alignment_list = list()
	for (i in 1:length(usaln_seqs))
		{
		usaln_name = usaln_names[i]
		usaln_seq = usaln_seqs[[i]]
		usaln_positions = 1:length(usaln_seq)
		usaln_positions = usaln_positions[usaln_seq != "-"]
		usaln_positions

	
		TF = grepl(pattern=usaln_name, x=di3_fns)
		if (sum(TF) != 1)
			{
			txt = paste0("STOP ERROR in align_3dis(): no single *_3di.fasta found for alignment entry usaln_name='", usaln_name, "'. Please fix and re-run.")
			cat("\n")
			cat(txt)
			cat("\n")
			stop(txt)
			}

		di3_fn = di3_fns[TF]
		
		chars_aa_aligned = usaln_seq
		chars_3di = as.character(ape::read.FASTA(di3_fn, type="AA"))[[1]]	
		di3_aligned = align_3di_to_aa(chars_aa_aligned, chars_3di)

		di3_alignment_list[[usaln_name]] = di3_aligned
		}

	di3_alignment_list

	di3_alignment = ape::as.AAbin(di3_alignment_list)
	di3_chars = as.character(di3_alignment)

	cat("\nalign_3di() is writing 3di alignment to: '", di3_outfn, "'.\n", sep="")
	ape::write.FASTA(di3_alignment, file=di3_outfn)

	return(di3_alignment)
	}


str_to_3di <- function(str_fn, suffix="\\.cif")
	{	
	if (length(str_fn) > 1)
		{
		fns_df = NULL
		for (i in 1:length(str_fn))
			{
			tmpdf = str_to_3di(str_fn[i], suffix=suffix)
			fns_df = rbind(fns_df, tmpdf)
			}
		return(fns_df)
		}
	
	# Structure to 3di and fasta
	dump_fn = gsub(pattern=suffix, replacement="_3di.dump", x=str_fn)
	chain_names_fn = gsub(pattern=suffix, replacement="_chains.txt", x=str_fn)
	di3_fn = gsub(pattern=suffix, replacement="_3di.fasta", x=str_fn)
	aa_fn = gsub(pattern=suffix, replacement="_aa.fasta", x=str_fn)
	both_fn = gsub(pattern=suffix, replacement="_both.fasta", x=str_fn)
	
	cmd_txt = paste0("foldseek structureto3didescriptor ", str_fn, " ", dump_fn)
	system(cmd_txt)
	
	cmd_txt = paste0("awk -F'\t' '{print $1; next}{print}' ", dump_fn, " | tee ", chain_names_fn)
	system(cmd_txt)
	
	cmd_txt = paste0("awk -F'\t' '{print ", '">"', '$1"\\n"', "$3; next}{print}' ", dump_fn, " | tee ", di3_fn)
	system(cmd_txt)

	#cmd_txt2 = paste0("-F'\t' '{print ", '">"', '$1"\\n"', "$3; next}{print}' ", dump_fn, " | tee ", di3_fn)
	#system2(command="awk", args=cmd_txt2)

	cmd_txt = paste0("awk -F'\t' '{print ", '">"', '$1"\\n"', "$2; next}{print}' ", dump_fn, " | tee ", aa_fn)
	system(cmd_txt)
	
	cmd_txt = paste0("awk -F'\t' '{print ", '">"', '$1"\\n"', '$2"\\n"', "$3; next}{print}' ", dump_fn, " | tee ", both_fn)
	system(cmd_txt)
	
	
	# fns
	fns = c(str_fn, dump_fn, chain_names_fn, di3_fn, aa_fn, both_fn)
	fns_df = as.data.frame(matrix(fns, nrow=1), stringsAsFactors=FALSE)
	row.names(fns_df) = NULL
	names(fns_df) = c("str_fn", "dump_fn", "chain_names_fn", "di3_fn", "aa_fn", "both_fn")
	
	return(fns_df)
	}


#######################################################
# Get the indices of the PDB structure residues with 
# confidence below a cutoff
#
# It should look at the same field, "b" or "B_iso_or_equiv",
# which represents the B-factor measure of structural uncertainty.
# 
# foldseek structureto3didescriptor 1afr_A_AF-B9T0X0-F1-model_v4.cif 1afr_A_AF-B9T0X0-F1-model_v4.dump --mask-bfactor-threshold 70
# awk -F'\t' '{print ">"$1"\n"$3; next}{print}' 1afr_A_AF-B9T0X0-F1-model_v4.dump
# 
# 
#######################################################
get_hiconf_residues <- function(pdb, mask_bfactor_threshold=70)
	{
	junk='
	library(bio3d)

	wd = "/GitHub/str2phy/ex/EXAMPLE_alphafold_to_3di/"
	setwd(wd)

	pdbfn = "1afr_A_AF-B9T0X0-F1-model_v4.pdb"
	pdb = read.pdb(pdbfn)
	
	mask_bfactor_threshold = 70
	
	names(pdb)
	names(pdb$atom)
	names(pdb$xyz)
	names(pdb$seqres)
	names(pdb$calpha)
	names(pdb$call)


	# B-factor / B_iso confidence is:
	head(pdb$atom$b)
	
	keep_resnums = get_hiconf_residues(pdb, mask_bfactor_threshold=70)
	' # END JUNK
	
	residues = pdb$atom[pdb$calpha,]
	
	# B-factors cutoff
	keepTF = residues$b >= mask_bfactor_threshold
	sum(keepTF)
	
	residues_kept = residues[keepTF,]
	
	keep_resnums = residues_kept$resno
	keep_resnums
	
	return(keep_resnums)
	} # END get_hiconf_residues <- function(pdb, mask_bfactor_threshold=70)


get_hiconf_atoms_by_residue <- function(pdb, mask_bfactor_threshold=70)
	{
	junk='
	library(bio3d)
	source("/GitHub/str2phy/Rsrc/str2phy_v1.R")

	wd = "/GitHub/str2phy/ex/EXAMPLE_alphafold_to_3di/"
	setwd(wd)

	pdbfn = "1afr_A_AF-B9T0X0-F1-model_v4.pdb"
	pdb = read.pdb(pdbfn)
	
	mask_bfactor_threshold = 70
	
	names(pdb)
	names(pdb$atom)
	names(pdb$xyz)
	names(pdb$seqres)
	names(pdb$calpha)
	names(pdb$call)


	# B-factor / B_iso confidence is:
	head(pdb$atom$b)
	
	keep_resnums = get_hiconf_residues(pdb, mask_bfactor_threshold=70)
	atom_nums = get_hiconf_atoms_by_residue(pdb, mask_bfactor_threshold=70)
	
	' # END JUNK
	
	residues = pdb$atom[pdb$calpha,]
	
	# B-factors cutoff
	keepTF = residues$b >= mask_bfactor_threshold
	sum(keepTF)
	
	residues_kept = residues[keepTF,]
	
	# Get the atoms, from residues with 
	atoms_to_keep_TF = pdb$atom$resno %in% residues_kept$resno 
	
	atom_nums = (1:length(atoms_to_keep_TF))[atoms_to_keep_TF]
	
	return(atom_nums)
	} # END get_hiconf_residues <- function(pdb, mask_bfactor_threshold=70)


trim_pdb_bfactor <- function(pdb, mask_bfactor_threshold=70)
	{
	#atom_nums = get_hiconf_atoms_by_residue(pdb, mask_bfactor_threshold=mask_bfactor_threshold)
	keep_resnums = get_hiconf_residues(pdb, mask_bfactor_threshold=mask_bfactor_threshold)
	subset_pdb = bio3d::trim.pdb(pdb, inds=bio3d::atom.select(pdb, resno=keep_resnums))
	subset_pdb
	
	return(subset_pdb)
	} # END trim_pdb_bfactor <- function(pdb, mask_bfactor_threshold=70)


trim_pdbfn_bfactor <- function(pdbfn, mask_bfactor_threshold=70, outfn="")
	{
	#atom_nums = get_hiconf_atoms_by_residue(pdb, mask_bfactor_threshold=mask_bfactor_threshold)
	pdb = bio3d::read.pdb(pdbfn)
	keep_resnums = get_hiconf_residues(pdb, mask_bfactor_threshold=mask_bfactor_threshold)
	subset_pdb = bio3d::trim.pdb(pdb, inds=bio3d::atom.select(pdb, resno=keep_resnums))
	subset_pdb
	
	# What to return
	if (outfn == "")
		{
		return(subset_pdb)
		} else if (outfn == "auto")
		{
		if (endsWith(x=pdbfn, suffix=".pdb") == TRUE)
			{
			outfn = gsub(pattern="\\.pdb", replacement="_sub.pdb", x=pdbfn)
			} else if (endsWith(x=pdbfn, suffix=".cif") == TRUE)
			{
			outfn = gsub(pattern="\\.cif", replacement="_sub.cif", x=pdbfn)
			} else if (endsWith(x=pdbfn, suffix=".mmcif") == TRUE)
			{
			outfn = gsub(pattern="\\.mmcif", replacement="_sub.mmcif", x=pdbfn)
			} else {
			outfn = paste0(pdbfn, "_sub")
			}
		}
	# Write the file
	bio3d::write.pdb(pdb=subset_pdb, file=outfn)
	
	return(outfn)
	} # END trim_pdb_bfactor <- function(pdb, mask_bfactor_threshold=70)


iqtree_file_to_model_df <- function(fn)
	{
	
	}
```
