## Supplementary material for "Tertiary-interaction characters enable fast, model-based structural phylogenetics beyond the twilight zone": 00_plot_xlsx_tree_comparisons_v1.R

```
library(openxlsx)
library(psych)

# Set working director
wd = "/GitHub/str2phy/ex/ferritins_M20/tree_comparison/"
setwd(wd)

# 
xlsfn = "60iqtree_runs_compared_v4subset_sumMatches_order.xlsx"
tmpxls = openxlsx::read.xlsx(xlsfn, startRow=3)

tmpxls$symbol[tmpxls$symbol == "d"] = "3"


tmpxls$sum_matches[tmpxls$sum_matches == 52] = 51 # unrooted tree has 51 bipartitions
#xls = xls[c(-1:-2),] # Remove the "+" symbol
tmpxls = tmpxls[c(-1),] # Remove the "+" symbol
orig_xls = tmpxls
xls = orig_xls
head(xls)
dim(xls)


#######################################################
# Pairs plot: all-against-all scatterplots
#######################################################
tmpdf = xls[,13:25]

pdffn = "pairs_13x13_v2.pdf"
pdf(file=pdffn, width=20, height=20)

pairs(tmpdf, pch=".")

psych::pairs.panels(tmpdf, 
             method = "pearson", # correlation method
						 hist.col = "#00AFBB",
             density = TRUE,  # show density plotspch=".")
						 lm = TRUE,
						 pch=symbols,
						 col=colors)
						 
dev.off()
cmdstr = paste0("open ", pdffn)
system(cmdstr)


#######################################################
# Legend
#######################################################
legend_lines = NULL
ll=0
legend_lines[(ll=ll+1)] = "3: 3Di characters-only"
legend_lines[(ll=ll+1)] = "A=amino-acids only"
legend_lines[(ll=ll+1)] = "B=Both AA+3Di characters used"
legend_lines[(ll=ll+1)] = "Blue: USalign+partitioned."
legend_lines[(ll=ll+1)] = "Light blue: USalign+not partitioned."
legend_lines[(ll=ll+1)] = "Orange: Famsa3di-aligned+partitioned."
legend_lines[(ll=ll+1)] = "Light orange: Famsa3di-aligned+not partitioned."
legend_lines[(ll=ll+1)] = "Red: No structural information used (AA-only)"
legend_lines[(ll=ll+1)] = "Italics: Structures came from Alphafold."
legend_lines[(ll=ll+1)] = "Bold: Sites with <35% data trimmed out."

xstart = 1
ystart = 1
yinc = -1
for (i in 1:ll)
	{
	
	}


#######################################################
# Choose a particular plot & annotate
#######################################################
fonts = rep(1, nrow(xls))
#base_pch = 0.8
#sizes = rep(base_pch, nrow(xls))
for (i in 1:nrow(xls))
	{
	if (xls$accent[i] == "italics")
		{
		fonts[i] = 3
		if (xls$size[i] == "S")
			{
			# Trimmed
			fonts[i] = 4
			}
		} else {
		fonts[i] = 1
		if (xls$size[i] == "S")
			{
			# Trimmed
			fonts[i] = 2
			}
		}
	}


minmax <- function(x, prettyTF=TRUE, na.rm=TRUE)
	{
	if (prettyTF == TRUE)
		{
		pretty_vals = pretty(x)
		minval = min(pretty_vals, na.rm=na.rm)
		maxval = max(pretty_vals, na.rm=na.rm)
		} else {
		minval = min(x, na.rm=na.rm)
		maxval = max(x, na.rm=na.rm)
		}
	lims = c(minval, maxval)
	return(lims)
	}

pdffn = "M20_bipartitions_vs_SPR_distance.pdf"
pdf(file=pdffn, width=6, height=6)

x = jitter(xls$sum_matches)
y = jitter(xls$spr)
xlab = "# bipartitions matching M20 tree"
ylab = "SPR distance to M20 tree"
xlim = minmax(x)
ylim = rev(minmax(y))

# Blank plot (white dots)
plot(x, y, pch=".", col="white", xlab=xlab, ylab=ylab, xlim=xlim, ylim=ylim)
# Add text
text(x, y, labels=xls$symbol, col=xls$color, cex=1, font=fonts)

dev.off()
cmdstr = paste0("open ", pdffn)
system(cmdstr)


pdffn = "M20_RF_symDiff_vs_SPR_distance.pdf"
pdf(file=pdffn, width=6, height=6)

x = jitter(xls$symDiff)
y = jitter(xls$spr)
xlab = "RF distance to M20 tree"
ylab = "SPR distance to M20 tree"
xlim = rev(minmax(x))
ylim = rev(minmax(y))

# Blank plot (white dots)
plot(x, y, pch=".", col="white", xlab=xlab, ylab=ylab, xlim=xlim, ylim=ylim)
# Add text
text(x, y, labels=xls$symbol, col=xls$color, cex=1, font=fonts)

dev.off()
cmdstr = paste0("open ", pdffn)
system(cmdstr)


rr = -1  # rr = remove rows
xls = orig_xls[rr,]
pdffn = "M20_RF_symDiff_vs_SPR_distance_v2.pdf"
pdf(file=pdffn, width=6, height=6)

x = jitter(xls$symDiff)
y = jitter(xls$spr)
xlab = "RF distance to M20 tree"
ylab = "SPR distance to M20 tree"
xlim = rev(minmax(x))
ylim = rev(minmax(y))

# Blank plot (white dots)
plot(x, y, pch=".", col="white", xlab=xlab, ylab=ylab, xlim=xlim, ylim=ylim)
# Add text
text(x, y, labels=xls$symbol, col=xls$color, cex=1, font=fonts[rr])

dev.off()
cmdstr = paste0("open ", pdffn)
system(cmdstr)


pdffn = "M20_RF_symDiff_vs_branchScoreDiff.pdf"
pdf(file=pdffn, width=6, height=6)

x = jitter(xls$symDiff)
y = jitter(xls$branchscoreDiff_s1)
xlab = "RF distance to M20 tree"
ylab = "Branch-score distance to M20 tree"
xlim = rev(minmax(x))
ylim = rev(minmax(y))

# Blank plot (white dots)
plot(x, y, pch=".", col="white", xlab=xlab, ylab=ylab, xlim=xlim, ylim=ylim)
# Add text
text(x, y, labels=xls$symbol, col=xls$color, cex=1, font=fonts[rr])

dev.off()
cmdstr = paste0("open ", pdffn)
system(cmdstr)


#######################################################
# Matches: deep vs. shallow
#######################################################
shallow_matches = xls$sum_matches - xls$sum_deepnodes
hist(shallow_matches)

pdffn = "deep_vs_shallow_bipartitions.pdf"
pdf(file=pdffn, width=6, height=6)

x = jitter(xls$sum_deepnodes)
y = jitter(shallow_matches)
xlab = "# deep bipartitions matching M20 tree"
ylab = "# shallow bipartitions matching M20 tree"
xlim = minmax(x)
ylim = minmax(y)

# Blank plot (white dots)
plot(x, y, pch=".", col="white", xlab=xlab, ylab=ylab, xlim=xlim, ylim=ylim)
# Add text
text(x, y, labels=xls$symbol, col=xls$color, cex=1, font=fonts[rr])

dev.off()
cmdstr = paste0("open ", pdffn)
system(cmdstr)


#######################################################
# Do an NMMDS plot
#######################################################
```
