## Supplementary material for "Tertiary-interaction characters enable fast, model-based structural phylogenetics beyond the twilight zone": 00_tree-tree_compare_v1.R

```
library(ape)
library(BioGeoBEARS)

wd = "/GitHub/str2phy/ex/ferritins_M20/tree_comparison/"
setwd(wd)

distance_trfn = "Malik_2020_Fig10b_FerritinData_distance_matrix_BIONJtree.newick"
distance_tr = read.tree(distance_trfn)

plot(distance_tr)

cat(sort(distance_tr$tip.label), sep="\n")

length(distance_tr$tip.label)

# Yay, they match the Excel file!
```
