## Supplementary material for "Tertiary-interaction characters enable fast, model-based structural phylogenetics beyond the twilight zone": 01_all-against-all_tree_distances_v1.R

```
#######################################################
# All-against-all tree-to-tree distances
#######################################################

library(ape)
library(BioGeoBEARS)
library(Quartet)   # for Quartet distances
library(gdata)		# for trim
library(stringr)		# for ?
library(stringi)		# for fancy string search/replace
library(phangorn)		# for treedist
library(psych)			# for psych::pairs.panels
wd = "/GitHub/str2phy/ex/ferritins_M20/treedists_all-against-all/"
setwd(wd)

distance_trfn = "/GitHub/str2phy/ex/ferritins_M20/iqtree/Malik_2020_Fig10b_FerritinData_distance_matrix_BIONJtree.newick"
distance_tr = read.tree(distance_trfn)


#######################################################
# Calculate bipartition matches to distances tree
#######################################################

fn = "/GitHub/str2phy/ex/ferritins_M20/tree_comparison/output_filenames_v1.txt"
lines = readLines(fn, skipNul=TRUE)

TF = startsWith(x=lines, prefix="#")
lines = lines[TF==FALSE]
TF = lines == ""
lines = lines[TF==FALSE]
length(lines)

trfns = c(distance_trfn,
lines)


analysis_names = c("M20_distances_tr",
"iqtree_FAMSA_AAonly_trim35",
"iqtree_FAMSA_AAonly_trim35_allmodels",
"iqtree_FAMSA_AAonly_full",
"iqtree_FAMSA_AAonly_full_allmodels",
"iqtree_FAMSA3di_3diFull",
"iqtree_FAMSA3di_3diFull_allmodels",
"iqtree_FAMSA3di_3diTrim35",
"iqtree_FAMSA3di_3diTrim35_allmodels",
"iqtree_FAMSA3di_AAfull",
"iqtree_FAMSA3di_AAfull_allmodels",
"iqtree_FAMSA3di_AA_trim35",
"iqtree_FAMSA3di_AA_trim35_allmodels",
"iqtree_FAMSA3di_BothFull",
"iqtree_FAMSA3di_BothFull_allmodels",
"iqtree_FAMSA3di_BothFullNonPart",
"iqtree_FAMSA3di_BothFullNonPart_allmodels",
"iqtree_FAMSA3di_BothTrim35",
"iqtree_FAMSA3di_BothTrim35_allmodels",
"iqtree_FAMSA3di_BothTrim35NonPart",
"iqtree_FAMSA3di_BothTrim35NonPart_allmodels",
"iqtree_USaln_3di_Full",
"iqtree_USaln_3di_Trim",
"iqtree_USaln_AAs_Full",
"iqtree_USaln_AAs_Trim",
"iqtree_USaln_Both_Full",
"iqtree_USaln_Both_Full_allmodels",
"iqtree_USaln_Both_Trim",
"iqtree_USaln_Both_Trim_allmodels",
"iqtrAF_FAMSA3di_3diFull",
"iqtrAF_FAMSA3di_3diFull_allmodels",
"iqtrAF_FAMSA3di_3diTrim35",
"iqtrAF_FAMSA3di_3diTrim35_allmodels",
"iqtrAF_FAMSA3di_AAfull",
"iqtrAF_FAMSA3di_AAfull_allmodels",
"iqtrAF_FAMSA3di_AA_trim35",
"iqtrAF_FAMSA3di_AA_trim35_allmodels",
"iqtrAF_FAMSA3di_BothFull",
"iqtrAF_FAMSA3di_BothFull_allmodels",
"iqtrAF_FAMSA3di_BothFullNonPart",
"iqtrAF_FAMSA3di_BothFullNonPart_allmodels",
"iqtrAF_FAMSA3di_BothTrim35",
"iqtrAF_FAMSA3di_BothTrim35_allmodels",
"iqtrAF_FAMSA3di_BothTrim35NonPart",
"iqtrAF_FAMSA3di_BothTrim35NonPart_allmodels",
"iqtrAF_USaln_3diFull",
"iqtrAF_USaln_3diFull_allmodels",
"iqtrAF_USaln_3diTrim35",
"iqtrAF_USaln_3diTrim35_allmodels",
"iqtrAF_USaln_AAfull",
"iqtrAF_USaln_AAfull_allmodels",
"iqtrAF_USaln_AA_trim35",
"iqtrAF_USaln_AA_trim35_allmodels",
"iqtrAF_USaln_BothFull",
"iqtrAF_USaln_BothFull_allmodels",
"iqtrAF_USaln_BothFullNonPart",
"iqtrAF_USaln_BothFullNonPart_allmodels",
"iqtrAF_USaln_BothTrim35",
"iqtrAF_USaln_BothTrim35_allmodels",
"iqtrAF_USaln_BothTrim35NonPart",
"iqtrAF_USaln_BothTrim35NonPart_allmodels")

partitioned_modelnames = c("iqtree_FAMSA3di_BothFull",
"iqtree_FAMSA3di_BothFull_allmodels",
"iqtree_FAMSA3di_BothTrim35",
"iqtree_FAMSA3di_BothTrim35_allmodels",
"iqtree_USaln_Both_Full",
"iqtree_USaln_Both_Full_allmodels",
"iqtree_USaln_Both_Trim",
"iqtree_USaln_Both_Trim_allmodels",
"iqtrAF_FAMSA3di_BothFull",
"iqtrAF_FAMSA3di_BothFull_allmodels",
"iqtrAF_FAMSA3di_BothTrim35",
"iqtrAF_FAMSA3di_BothTrim35_allmodels",
"iqtrAF_USaln_BothFull",
"iqtrAF_USaln_BothFull_allmodels",
"iqtrAF_USaln_BothTrim35",
"iqtrAF_USaln_BothTrim35_allmodels")


trs_list = list()

txt = paste0("\nCalculating statistics for ", length(trfns), " tree files. Calc #: ")
cat(txt)
for (i in 1:length(trfns))
	{
	cat(i, ",", sep="")
	
	# Make branchlengths relative so that all tree-lengths = 1.0
	tmptr = read.tree(trfns[i])
	tmptr$edge.length = tmptr$edge.length / sum(tmptr$edge.length)
	tmptr$tip.label = gsub(pattern="\\.pdb", replacement="", x=tmptr$tip.label)
	tmptr = phytools::midpoint.root(tmptr)
	tmptr = ladderize(tmptr)
	# Read back in
	tmptr = read.tree(file="", text=write.tree(tmptr))
	
	trs_list[[i]] = tmptr
	} # END 

cat("...done!\n")
class(trs_list) = c("list", "multiPhylo")

#######################################################
# Calculate all-against-all tree-to-tree distances
#######################################################
n = length(trfns)
RFdistmat = matrix(data=0, nrow=n, ncol=n)
SPRdistmat = matrix(data=0, nrow=n, ncol=n)
BranchScore_distmat = matrix(data=0, nrow=n, ncol=n)

txt = paste0("\nCalculating all-versus-all distances for ", n", trees. i=")
cat(txt)
for (i in 1:n)
	{
	cat(i, ",", sep="")
	for (j in i:n)
		{
		tr1 = trs_list[[i]]
		tr2 = trs_list[[j]]
		
		# Tree-to-tree distances
		tdists = phangorn::treedist(tree1=tr1, tree2=tr2, check.labels=TRUE)
		RFdistmat[i,j] = tdists["symmetric.difference"]
		RFdistmat[j,i] = tdists["symmetric.difference"]
		BranchScore_distmat[i,j] = tdists["branch.score.difference"]
		BranchScore_distmat[j,i] = tdists["branch.score.difference"]
		sprdist = phangorn::sprdist(tree1=tr1, tree2=tr2)
		SPRdistmat[i,j] = sprdist["spr"]
		SPRdistmat[j,i] = sprdist["spr"]
		} # END for (j in 1:n)
	} # END for (i in 1:n)
cat("\n...done")

# Nonmetric Multidimensional Scaling (NMMDS)
library(vegan)
RF_nmmds = vegan::metaMDS(comm=RFdistmat)
SPR_nmmds = vegan::metaMDS(comm=SPRdistmat)
BranchScore_nmmds = vegan::metaMDS(comm=BranchScore_distmat)


# Load plotting info
xlsfn = "/GitHub/str2phy/ex/ferritins_M20/tree_comparison/60iqtree_runs_compared_v4subset_sumMatches_order.xlsx"
tmpxls = openxlsx::read.xlsx(xlsfn, startRow=3)
tmpxls$symbol[tmpxls$symbol == "d"] = "3"
tmpxls$sum_matches[tmpxls$sum_matches == 52] = 51 # unrooted tree has 51 bipartitions
#xls = xls[c(-1:-2),] # Remove the "+" symbol
tmpxls = tmpxls[c(-1),] # Remove the "+" symbol
orig_xls = tmpxls
xls = orig_xls
head(xls)
dim(xls)

fonts = rep(1, nrow(xls))
#base_pch = 0.8
#sizes = rep(base_pch, nrow(xls))
for (i in 1:nrow(xls))
	{
	if (xls$accent[i] == "italics")
		{
		fonts[i] = 3
		if (xls$size[i] == "S")
			{
			# Trimmed
			fonts[i] = 4
			}
		} else {
		fonts[i] = 1
		if (xls$size[i] == "S")
			{
			# Trimmed
			fonts[i] = 2
			}
		}
	}

# default plot
plot(RF_nmmds)


# NMDS plot of RF distances
pdffn = "RF_nmmds_v1.pdf"
pdf(file=pdffn, width=6, height=6)

x = RF_nmmds$points[,"MDS1"]
y = RF_nmmds$points[,"MDS2"]
xlab = "NMDS1"
ylab = "NMDS2"

# Blank plot (white dots)
plot(x, y, pch=".", col="white", xlab=xlab, ylab=ylab)
# Add text
text(x, y, labels=xls$symbol, col=xls$color, cex=1, font=fonts)
title("Nonmetric Multi-dimensional Scaling (NMDS)\nRF distances between 61 trees", cex=0.8)

stress_txt = paste0("stress = ", round(RF_nmmds$stress, digits=2))
text(x=min(x), y=max(y), label=stress_txt, adj=0)

dev.off()
cmdstr = paste0("open ", pdffn)
system(cmdstr)


# NMDS plot of SPR distances
pdffn = "SPR_nmmds_v1.pdf"
pdf(file=pdffn, width=6, height=6)

x = SPR_nmmds$points[,"MDS1"]
y = SPR_nmmds$points[,"MDS2"]
xlab = "NMDS1"
ylab = "NMDS2"

# Blank plot (white dots)
plot(x, y, pch=".", col="white", xlab=xlab, ylab=ylab)
# Add text
text(x, y, labels=xls$symbol, col=xls$color, cex=1, font=fonts)
title("Nonmetric Multi-dimensional Scaling (NMDS)\nSPR distances between 61 trees", cex=0.8)

stress_txt = paste0("stress = ", round(SPR_nmmds$stress, digits=2))
text(x=min(x), y=max(y), label=stress_txt, adj=0)

dev.off()
cmdstr = paste0("open ", pdffn)
system(cmdstr)


# NMDS plot of BranchScore distances
pdffn = "BranchScore_nmmds_v1.pdf"
pdf(file=pdffn, width=6, height=6)

x = BranchScore_nmmds$points[,"MDS1"]
y = BranchScore_nmmds$points[,"MDS2"]
xlab = "NMDS1"
ylab = "NMDS2"

# Blank plot (white dots)
plot(x, y, pch=".", col="white", xlab=xlab, ylab=ylab)
# Add text
text(x, y, labels=xls$symbol, col=xls$color, cex=1, font=fonts)
title("Nonmetric Multi-dimensional Scaling (NMDS)\nBranchScore distances between 61 trees", cex=0.8)

stress_txt = paste0("stress = ", round(BranchScore_nmmds$stress, digits=2))
text(x=min(x), y=max(y), label=stress_txt, adj=0)

dev.off()
cmdstr = paste0("open ", pdffn)
system(cmdstr)


# Add together RF and Branchscore
tmp1 = RFdistmat / sum(RFdistmat)
tmp2 = BranchScore_distmat / sum(BranchScore_distmat)
sum_distmat = tmp1 + tmp2
sumDists_nmmds = vegan::metaMDS(comm=sum_distmat)

# NMDS plot of BranchScore distances
pdffn = "RF+BranchScore_nmmds_v1.pdf"
pdf(file=pdffn, width=6, height=6)

x = sumDists_nmmds$points[,"MDS1"]
y = sumDists_nmmds$points[,"MDS2"]
xlab = "NMDS1"
ylab = "NMDS2"

# Blank plot (white dots)
plot(x, y, pch=".", col="white", xlab=xlab, ylab=ylab)
# Add text
text(x, y, labels=xls$symbol, col=xls$color, cex=1, font=fonts)
title("Nonmetric Multi-dimensional Scaling (NMDS)\nSum of normalized RF+BS distances between 61 trees", cex=0.8)

stress_txt = paste0("stress = ", round(sumDists_nmmds$stress, digits=2))
text(x=min(x), y=max(y), label=stress_txt, adj=0)

dev.off()
cmdstr = paste0("open ", pdffn)
system(cmdstr)
```
