## Supplementary figures and images for "Tertiary-interaction characters enable fast, model-based structural phylogenetics beyond the twilight zone"

### Fig1_foldseek_3di_alphabet_v1.pdf

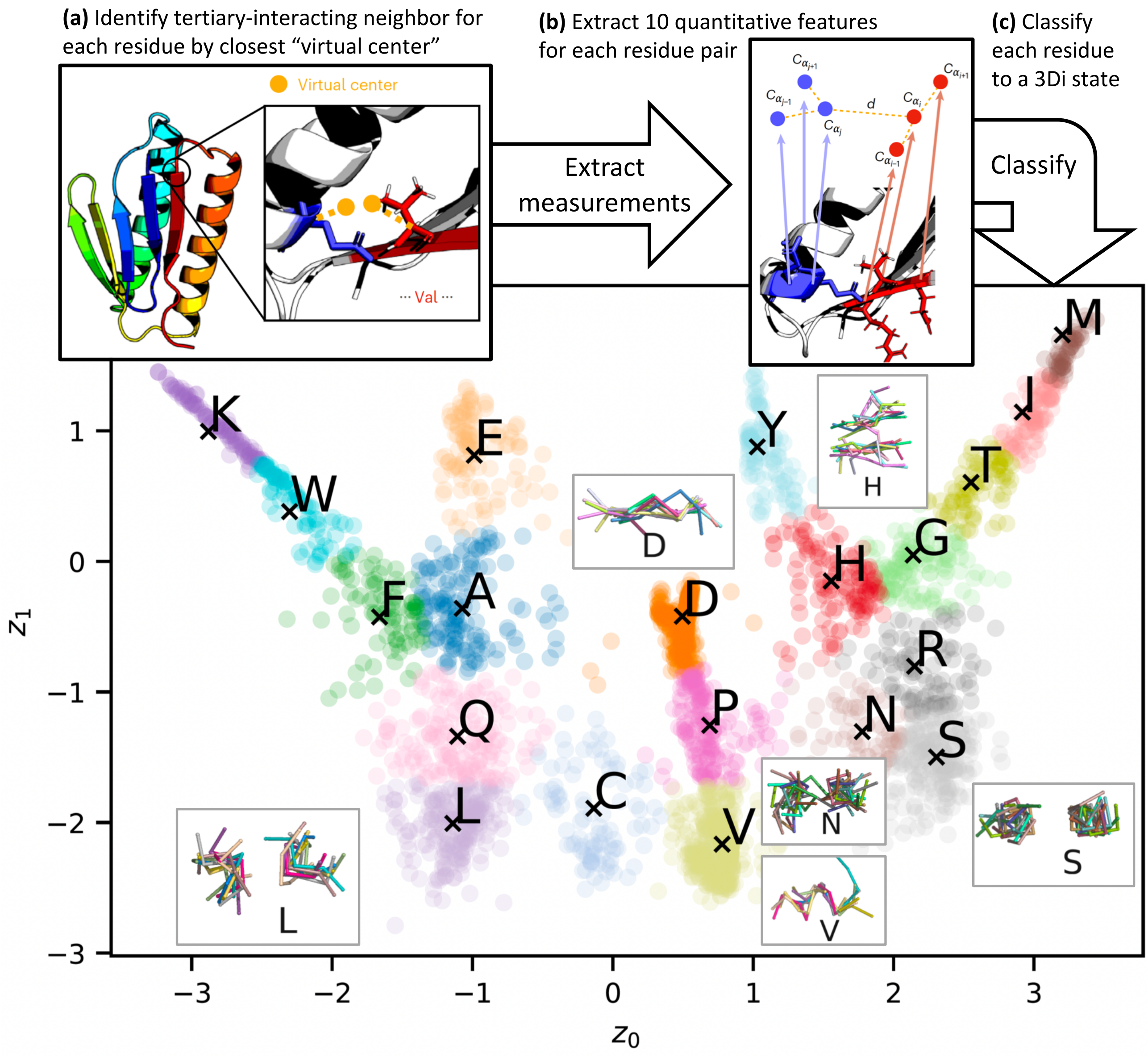

### Fig2_AA_3di_compared_v3.pdf

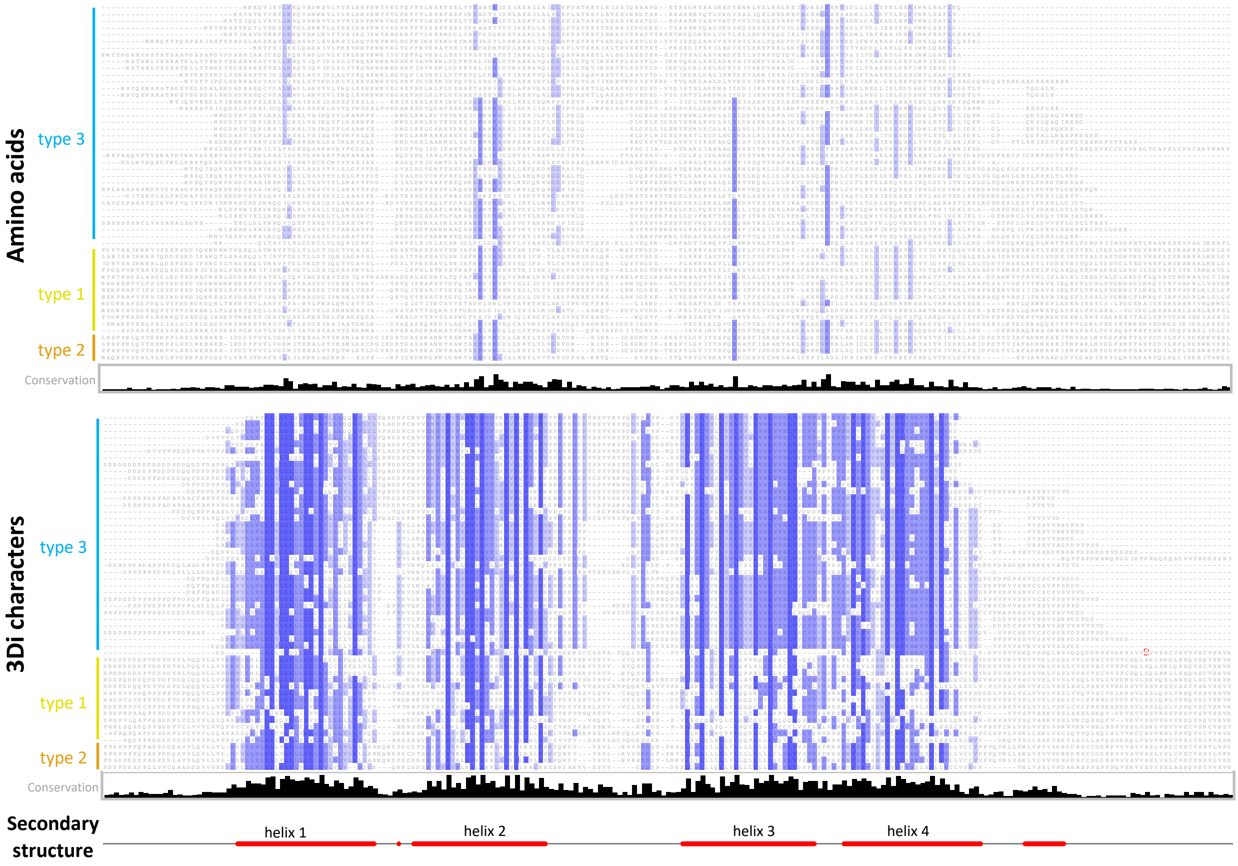

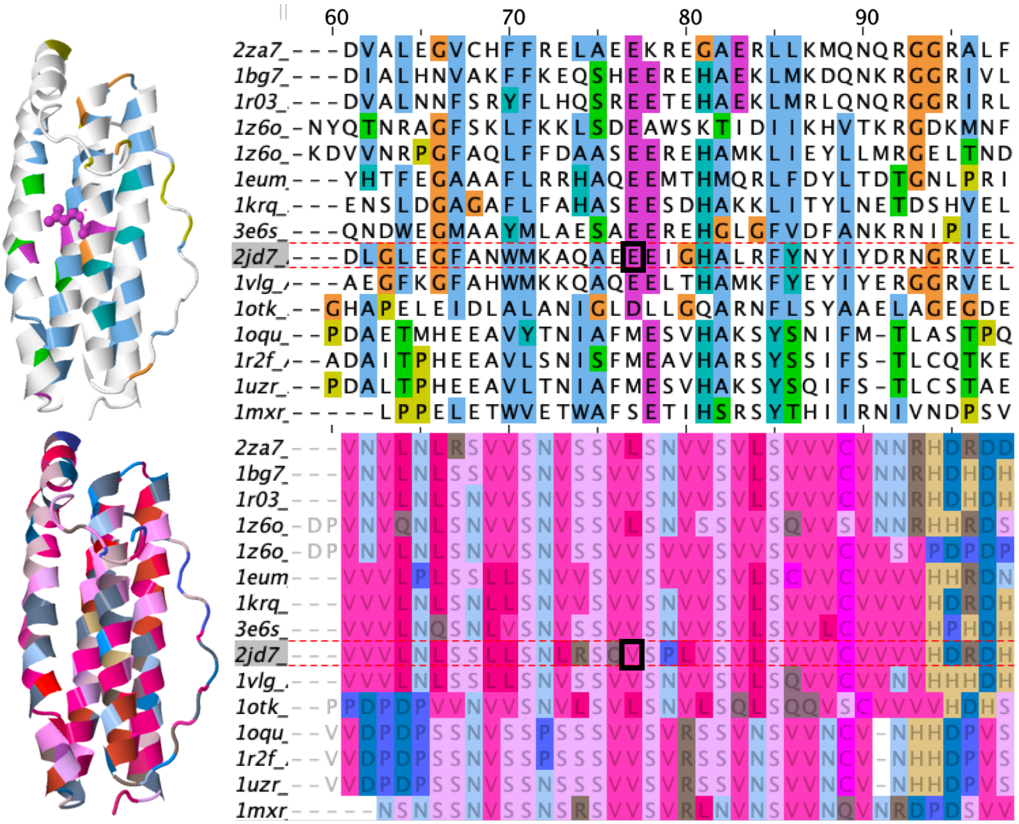

### Fig3_AA_3di_compared_Zoomed_v4.pdf

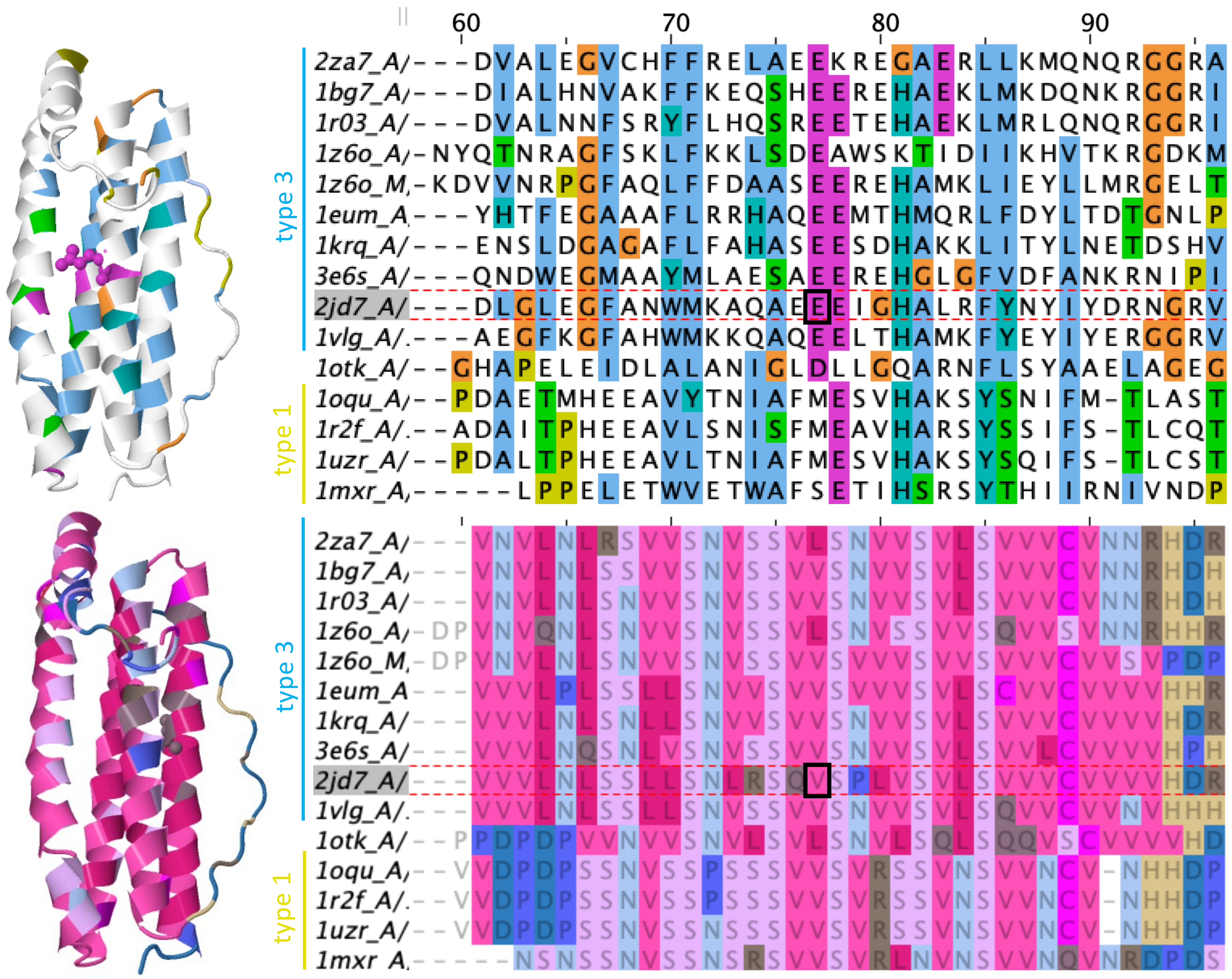

### Fig3_AA_3di_compared_Zoomed_v4.pptx

#### Slide 1
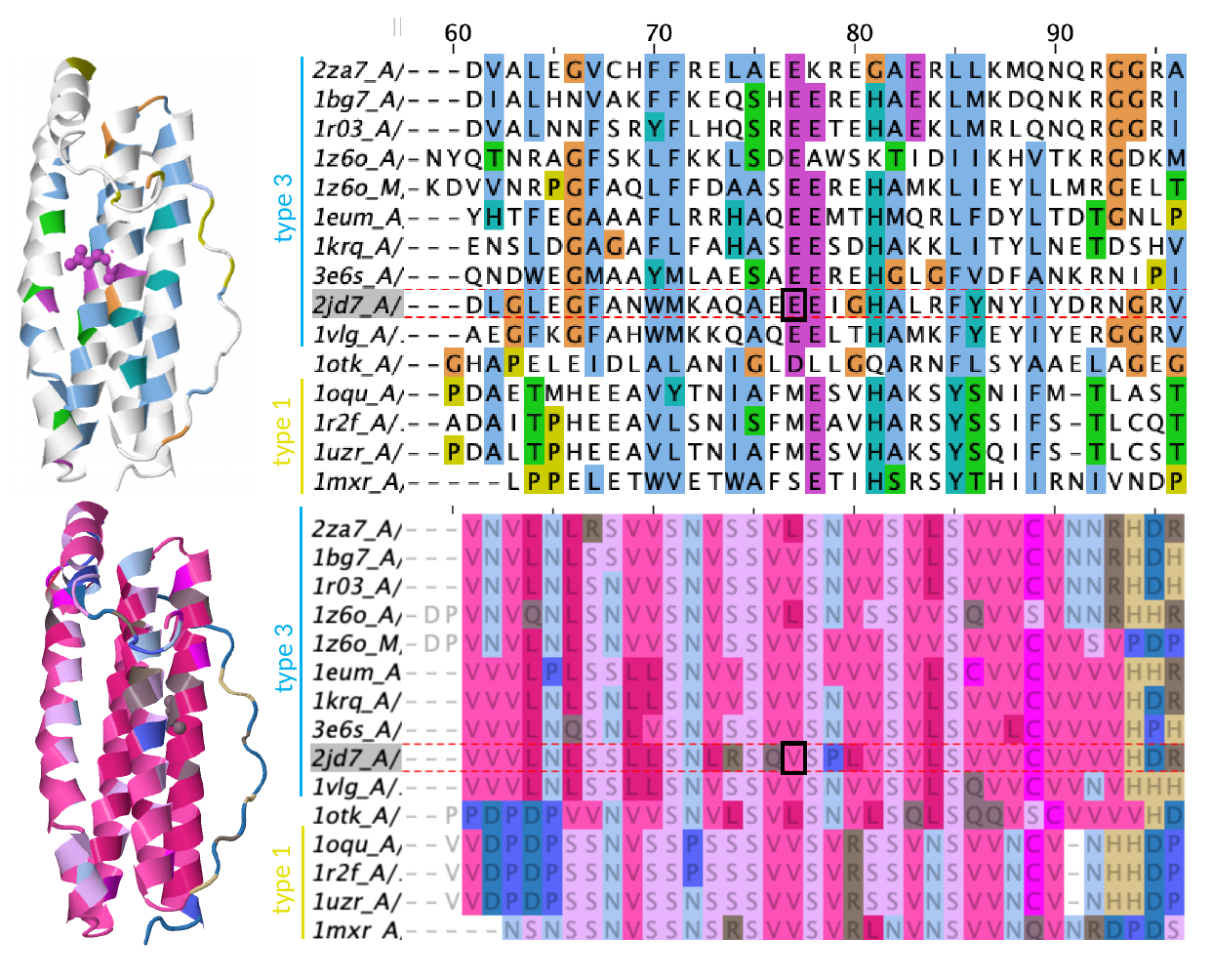

type 3
type 1
type 3
type 1

### VanKempen_etal_2023_SuppFig2_color_scheme.png

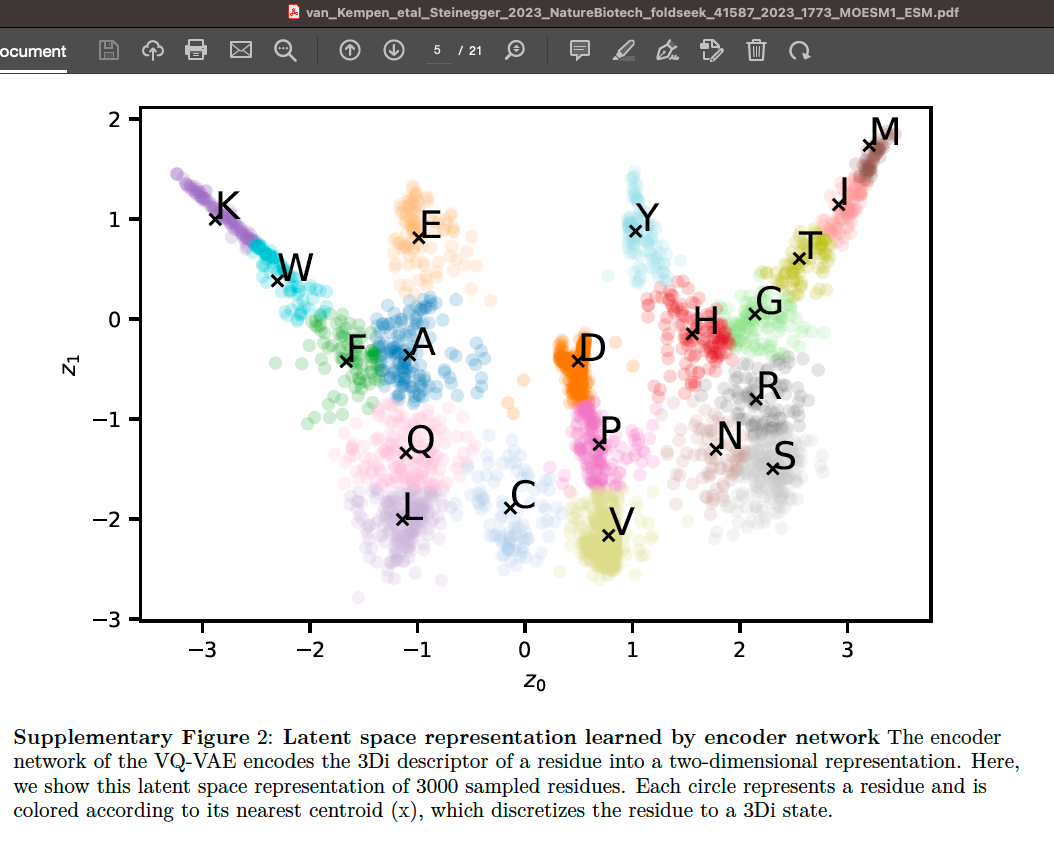
